# Hippocampal-medial prefrontal event segmentation and integration contribute to episodic memory formation

**DOI:** 10.1101/2020.03.14.990002

**Authors:** Wei Liu, Yingjie Shi, James N. Cousins, Nils Kohn, Guillén Fernández

**Affiliations:** School of Psychology, Central China Normal University (CCNU), Wuhan, China; Donders Institute for Brain, Cognition and Behaviour, Radboud University Medical Centre, Nijmegen, The Netherlands

**Keywords:** subsequent memory effect, hippocampus, medial prefrontal cortex, event segmentation, event integration

## Abstract

How do we encode our continuous life experiences for later retrieval? Theories of event segmentation and integration suggest that the hippocampus binds separately represented events into an ordered narrative. Using a functional Magnetic Resonance Imaging (fMRI) movie watching-recall dataset, we quantified two types of neural similarities (i.e., *activation pattern* similarity and within-region voxel-based *connectivity pattern* similarity) between separate events during movie watching and related them to subsequent retrieval of events as well as retrieval of sequential order. We demonstrated that compared to forgotten events, successfully remembered events were associated with distinct *activation patterns* in the hippocampus and medial prefrontal cortex. By contrast, similar *connectivity patterns* between events were associated with memory formation and were also relevant for retaining events in the correct order. We applied the same approaches to an independent movie watching fMRI dataset as validation and highlighted again the role of hippocampal activation pattern and connectivity pattern in memory formation. We propose that distinct *activation patterns* represent neural segmentation of events while similar *connectivity patterns* encode context information, and therefore integrate events into a narrative. Our results provide novel evidence for the role of hippocampal-medial prefrontal event segmentation and integration in episodic memory formation of real-life experience.

## Introduction

How we form memories of our life experiences is a fundamental scientific question with broad implications. In the past two decades, human neuroimaging and electrophysiology studies using the subsequent memory effect paradigm have implicated a distinct set of brain regions involved in successful memory formation (Brewer et al. 1998; Wagner et al. 1998; Fernández et al. 1999; Kim 2011). In these subsequent memory studies, increased neural activity of the hippocampus, parahippocampal gyrus, and the prefrontal cortex during memory encoding is associated with successful subsequent retrieval. However, real-world memories are formed based on a continuous stream of information rather than the sequentially presented, isolated items used in most subsequent memory studies (Kim 2011). Potentially, continuous sensory experience is segmented into distinct events (i.e., *event segmentation*) (Baldassano et al. 2017; Zacks 2020) that are then bound together into a coherent narrative, preserving their sequential relationships (i.e., *event integration*) (Griffiths and Fuentemilla 2020). To examine episodic memory formation of real-life-like experiences in humans, we analysed brain activity using functional Magnetic Resonance Imaging (fMRI) while participants were watching a movie. Based on subsequent memory recall, we aimed at identifying brain regions and neural representational processes underlying event segmentation and integration during episodic memory formation.

Thanks to recent advances in statistical analysis of ongoing neural activity (Hermans et al. 2011; Cohen et al. 2017; Xue 2018; Nastase et al. 2019), naturalistic stimuli (e.g., *movie, spoken narratives, music*) have been increasingly used in neuroscience (Hasson et al. 2004; Hermans et al. 2011; Huk et al. 2018; Sonkusare et al. 2019). This is especially valuable for memory research because naturalistic stimuli can greatly enhance the ecological validity of experimental studies (Hasson et al. 2008; Baldassano et al. 2017; Chen et al. 2017; Montchal et al. 2019). Hasson and colleagues first investigated memory formation with cinematographic stimuli and demonstrated that brain activity was more correlated among participants for later remembered than forgotten events (Hasson et al. 2008). While that study uncovered regions that encode continuous experiences, the nature of representations in those regions remained unclear, particularly with regard to how episodes are segmented into separate events and then integrated into a coherent sequence.

Event segmentation theory suggests that continuous experiences need to be segmented into discrete event representations, and thereafter they can be better understood and encoded (Zacks et al. 2001, 2007; Zacks 2020). Two recent studies provided novel perspectives into segmentation theory. Using Multi-Voxel Pattern Analysis (MVPA) and a movie watching-recall dataset, Chen and colleagues showed similar *activation patterns* of the same events across individuals and event-specific reinstatements of *activation patterns* between encoding and retrieval (Chen et al. 2017). Following this, Baldassano and colleagues demonstrated a nested processing hierarchy of events (‘*hierarchical memory system*’, (Hasson et al. 2015)) from fine-grained segmentation in early sensory regions to coarse segmentation in regions of the higher-order default-mode network (e.g., *medial prefrontal cortex (mPFC) and posterior medial cortex (PMC)*). Importantly, boundaries of long events at the top of the hierarchy matched with event boundaries annotated by human observers and were coupled to increased hippocampal activity (Baldassano et al. 2017). These results demonstrated that human brains spontaneously used different *activation patterns* to represent events during continuous movie watching, and how these *activation patterns* reactivated during recall. Also, it may suggest that regions such as mPFC, PMC, and hippocampus encode events at the same level that we consciously perceive boundaries between events. However, it remains unclear how exactly this event segmentation at the neural level relates to subsequent memory recall.

Event segmentation alone is not sufficient for episodic memory formation of continuous real-life experiences. Event perception theory suggests that it is essential to also integrate segmented events into a coherent narrative via time, meaning, or other abstract features (Bauer and Varga 2017; Griffiths and Fuentemilla 2020). Therefore, a non-exhaustive list of questions are: (1) what are the neural underpinnings of event integration during continuous memory formation, (2) does integration occur in the same brain regions as segmentation, and (3) how does integration relate to subsequent memory recall. A promising approach to answer these questions is to examine local *connectivity patterns* (also called *multi-voxel correlation structure*), which may represent a brain signal that integrates events (Tambini and Davachi 2019). This method was derived from rodent electrophysiology (Qin et al. 1997; Kudrimoti et al. 1999; Lansink et al. 2008) and has been used in human fMRI studies (Tambini and Davachi 2013; Hermans et al. 2017) to quantify distributed memory representations in neuronal assemblies. Recently, Tambini and Davachi (Tambini and Davachi 2019) proposed that *activation patterns* are the representations of specific perceptual inputs (e.g., *stimuli*), while local *connectivity patterns* reflect particular encoding contexts or states. However, the different mnemonic functions of *activity patterns* and *connectivity patterns* have yet to be compared empirically within a single study. If local *connectivity patterns* represent encoding context, they may facilitate integration across events. Examination of *connectivity patterns* alongside *activation patterns* would help to characterise how the brain simultaneously performs event segmentation and integration.

Recently, a hippocampal neural code that simultaneously tracked subdivisions of a continuous experience (i.e., *events*) and their sequential relationship was described in rodents’ CA1 region (Sun et al. 2020). This neural code could be a fundamental neural correlate by which episodic experience is segmented and integrated, but has yet to be revealed in humans. Univariate hippocampal activity was found to increase at the boundaries between two events during continuous experience (Ben-Yakov and Dudai 2011; Ben-Yakov et al. 2013; DuBrow and Davachi 2013; Baldassano et al. 2017; Ben-Yakov and Henson 2018), but what these hippocampal signals represent in terms of event segmentation and integration is not clear. Theoretical models proposed that increased hippocampal signal may reflect a rapid shift in mental representations (e.g., *temporal and/or contextual information of an event*) (Ranganath and Ritchey 2012; DuBrow and Davachi 2016; DuBrow et al. 2017). Therefore, it can be regarded as the neural signature of event segmentation. Alternatively, this increase may link to the integration of episodic memories across event boundaries, as suggested by scalp electrocorticography (EEG) studies (Sols et al. 2017; Silva et al. 2019) and the event conjunction framework (Griffiths and Fuentemilla 2020). Previous fMRI studies of associative inference paradigms (mainly the AB-AC paradigm) suggested that integration of separate elements in memory may rely on coordinated neural activity between the hippocampus and mPFC (Zeithamova et al. 2012; Preston and Eichenbaum 2013; Schlichting et al. 2014; Tompary and Davachi 2017). Additional fMRI evidence came from tasks in which artificial boundaries were created within sequentially presented sentences and/or pictures: boundary-related activation of the lateral PFC and hippocampus, measured by fMRI, is involved in bridging neighboring elements at recall (Ezzyat and Davachi 2011; DuBrow and Davachi 2016). Moreover, Chen and colleagues showed hippocampal involvement in bridging narrative events across days (Chen et al. 2016). However, fMRI evidence for the role of hippocampal signals in online event integration when participants process continuous experience remains limited.

The current study aimed to reveal the neural underpinnings of the two processes in question – event segmentation and event integration - during memory formation of naturalistic experiences. To that end, we first analyzed an existing dataset (Baldassano et al. 2017; Chen et al. 2017) where participants watched a movie while being scanned (**Figure 1A**) and afterwards were instructed to freely recall the story of the movie (**Figure 1B**). This design allowed us to associate different neural measures during episodic encoding with subsequent memory retrieval (**Figure 1C-D**). We extracted voxel-wise Blood Oxygenation Level Dependent (BOLD) time courses during movie watching (encoding) from six predefined regions-of-interest (ROI) in the ‘*hierarchical memory system*’ (Hasson et al. 2015) including early auditory and visual areas, posterior medial cortex, medial prefrontal cortex, hippocampus, and posterior parahippocampal gyrus (**Figure 2A**; **Figure S1**). To probe the role of a broader set of regions in event segmentation and integration, we repeated all analyses in each region of a neocortical parcellation (Schaefer et al. 2018) (**Figure 2B**). We first examined the relationship between ROI-based activity time courses and subsequent memory recall and replicated the classical subsequent memory effects (i.e., *greater activation for remembered compared to forgotten events*) in regions including the hippocampus as well as the posterior parahippocampal gyrus (**Figure S2-3**, details in **Supplementary Materials**). To dissociate the two event processes, we used voxel-wise activity (**Figure 2C**) from each ROI to quantify the similarity between neural representations of events by two different multivariate methods (i.e., *activation* and *connectivity patterns)* (**Figure 2D-E**). Before linking neural pattern similarities with subsequent memory, we first compared *between-event* and *within-event* pattern similarities. We predicted that if our multivariate methods capture event representations, *within-event* pattern shifts should be smaller than *between-event* pattern shifts. Then we reasoned that if the neural representation (*activation* or *connectivity pattern*) shows a large transition (i.e., *lower neural similarity value*) between two adjacent events, and if this dissimilarity associates with better subsequent memory for events, then this representation might be involved in event segmentation (**Figure 2E**). By contrast, if the neural representation remains stable (i.e., *higher similarity*) across two or more neighboring events, and this stability relates to event memory as well as retention of the correct order for those events (i.e., *order memory*), then this representation may underlie event integration (**Figure 2F**). The relationship between neural event processing (i.e., *segmentation and event integration*) and memory formation was further cross-validated in an independent movie watching-recall dataset (*replication dataset*) that used a different experimental protocol with alternative movie stimuli.

**Figure 1.**
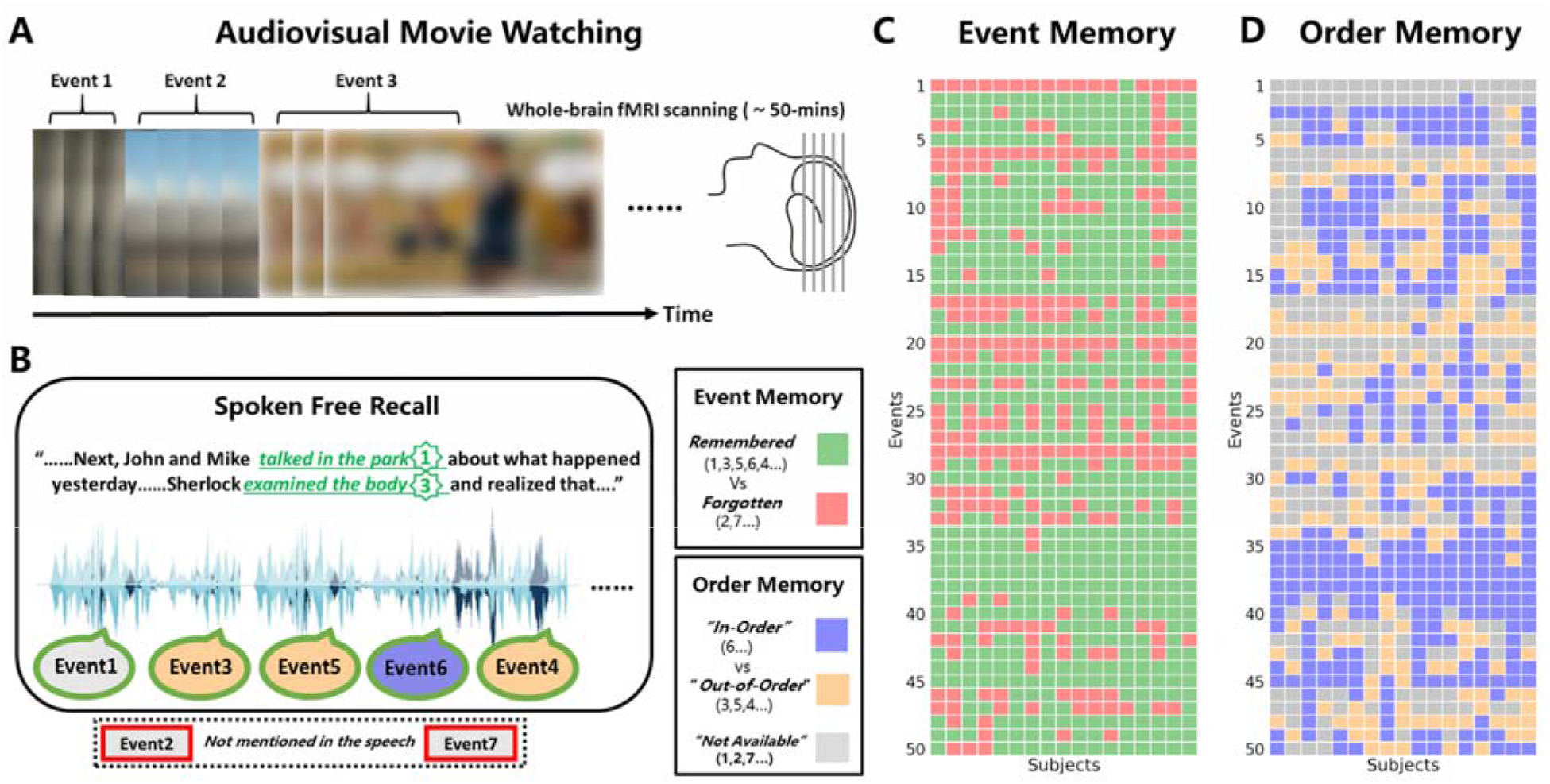
Experimental procedure and behavioural performance in the discovery dataset. **(A)** Each participant watched a 50-min audiovisual movie, BBC’s Sherlock (season 1, episode 1), while brain activity was recorded with fMRI. The movie was divided into 50 events based on major narrative shifts. Blurred images are shown here due to copyright reasons. However, the movie was shown in high resolution during the experiment. **(B)** Immediately after movie-watching, participants verbally recalled the movie content in as much detail as possible without any visual or auditory cues. Speech was recorded using a microphone and then transcribed. Critically, speech was also segmented into events and matched with the events segmented from the movie. All events mentioned in the speech were labelled as *remembered* while missing events were labelled as *forgotten*. In addition, among those remembered events, the ones that were recalled in the correct sequential order were labelled as *in-order* events (e.g., *event 6* was recalled after *event 5*). *Out-of-order* events were those that were recalled in an incorrect sequential order (e.g., *event 4* was recalled after *event 6*). We labelled the first recalled event and all *forgotten* events as *not available* because no sequential information can be accessed. **(C)** Illustration of all *remembere*d and *forgotten* events during movie-watching in all participants**. (D)** Illustration of all *in-orde*r and *out-of-order* events during movie watching in all participants. Each row of the heatmap is a different event, and each column represents a participant.

**Figure 2.**
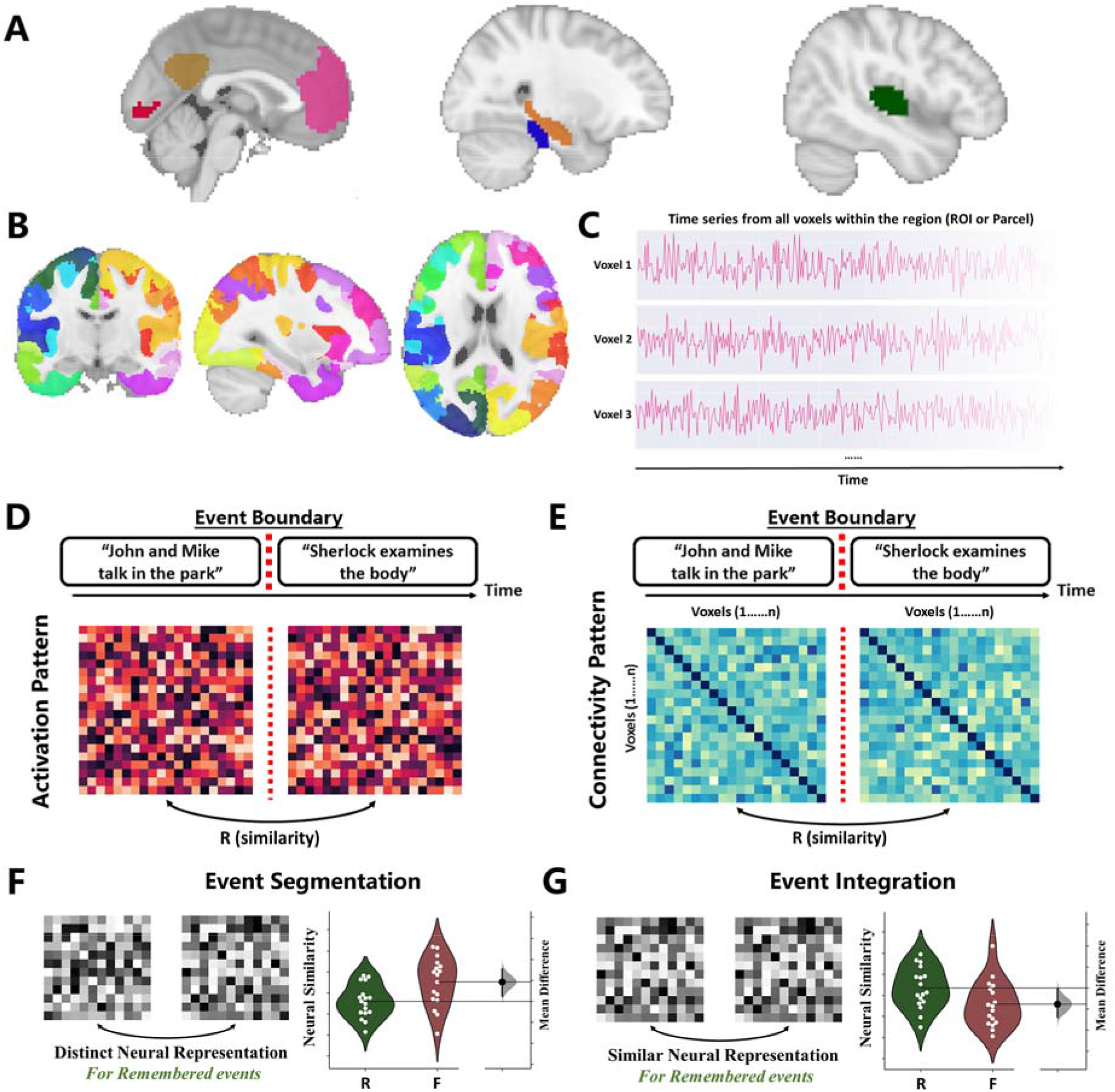
Neural similarities between separate events and their link with subsequent memory recall. **(A)** Six predefined regions-of-interest (ROIs): early auditory (green) and visual area (red), posterior medial cortex (brown), medial prefrontal cortex (pink), hippocampus (blue), and posterior parahippocampal gyrus (orange). See also Supplementary Figure 1. **(B)** Neocortical parcellation (1000 parcels) used in searchlight analysis. **(C)** For each region (ROI or parcel), voxel-wise signal during movie watching was extracted and then segmented into 50 events based on the event annotations. **(D)** We first generated event-specific *activation patterns* by averaging over all time points in that event. Then *activation pattern* similarity was calculated by Pearson’s correlation between *activation patterns* of two sequential events. If a region encodes two events separately, we expect two distinct neural representations and therefore a lower pattern similarity. (E) Event-specific within-region *connectivity patterns* were represented by voxel-by-voxel pairwise correlation matrices. *Connectivity pattern* similarity across event boundaries was also calculated using Pearson’s r between two sequential events. Stable neural representations across two events should yield a higher pattern similarity in the corresponding region. **(F)** fMRI evidence for event segmentation. For a certain multivariate neural measure, if it can be found that two distinct neural representations are used to encode the adjacent events while the neural patterns for remembered *(‘R’)* events are more dissimilar compared to forgotten *(‘F’)* events, this measure is likely to be associated with event segmentation. **(G)** fMRI evidence for event integration. If the multivariate neural measure remains stable across the boundary of two neighboring events and remembered *(‘R’)* events have higher neural similarity compared to forgotten *(‘F’)* events, this measure may relate to event integration.

## Methods

### 1. Participants and procedure

#### 1.1 Participants

##### Discovery dataset

Twenty-two healthy young adults (10 female, age range 18-26, mean age 20.8 years) participated in the experiment. All participants were native English speakers and naïve to the BBC crime drama *Sherlock.* Data were discarded from participants with excessive motion (> 1 voxel; n = 2), low recall duration (< 10 min; n = 2), or sleeping during the experiment (n = 1). This leaves 17 participants in total for our analyses. Due to a technical problem, one participant (s5) is missing data for the last 75 s (part of event 49 and all of event 50) and the affected two events were excluded in the analyses.

##### Replication dataset

In total 52 healthy adults (40 older adults (mean age=69 years) and 12 young adults (mean age=23 years)) from the St. Louis community or Washington University’s Psychology Department participant pool participated in this study. No participant reported current physical or mental health disorders. All of them were right-handed and had normal or corrected to normal vision. This research was approved by the Human Research Protection Office at Washington University

#### 1.2 Procedure

##### Discovery dataset

All our analyses are based on the Sherlock Movie Dataset (Baldassano et al. 2017; Chen et al. 2017); see *Data availability* below) acquired and pre-processed at Princeton Neuroscience Institute. No similar analysis or results (excluding behavioural results of recall accuracy) have been reported in previous studies using this dataset.

Participants were informed that they would watch a movie and would later be required to recall its content. They were then presented with a 48-min segment of the first episode of the *Sherlock* series (encoding phase), split into two parts of approximately equal length (23 min and 25 min) and presented in two consecutive blocks. A 30 s introductory cartoon clip was prepended before each block. Immediately after the movie presentation, participants were instructed to verbally describe the movie in as much detail as they could and for as long as they wished (recall phase). They were asked to recall the episode in the correct sequential order but were permitted to return to earlier points in the narrative if they remembered further content. Audio was simultaneously recorded by a customized MR-compatible recording system throughout the recall phase.

##### Replication dataset

A detailed description of the procedure can be found in the previous publication (Kurby and Zacks 2018). Participants finished two sessions with an interval of around 3.4 days (SD=2.3 days, min=0, max=14, mode=2). In session 1 (fMRI session), participants watched five movie clips in the same order. They were instructed to remember movie contents as much as possible. In session 2 (i.e., *Behavioral session*), two kinds of behavioral testing were performed. The first was the segmentation task, where participants were instructed to “*press a button to press a button when, in their opinion, one meaningful unit of activity ended and another began*” (Kurby and Zacks 2018). They produced both coarse (i.e., *largest meaningful units*) and fine (i.e., *smallest meaningful units*) segmentation for the same clip and we used the coarse segmentation as event boundaries in our neuroimaging analyses. Second, they performed memory tests. Recognition, recall, and order memory was tested for each movie clip. Recognition memory was tested using a 20-item two-alternative forced-choice test where participants were instructed to choose the visual image they saw in the movie instead of the distracter. Recall memory was assessed by asking participants to describe the movie content in as much detail as possible. Order memory was tested by reordering 12 visually distinctive images from the movie according to when they appeared.

### 2. Behavioural data analysis

#### Discovery dataset

##### Event annotations of the movie and verbal speech recording

The movie was segmented into 48 events by an independent observer who was blind to the experimental purpose, design, or results, following major shifts in the narrative (e.g., *changes in location, topic, and/or time*). Each event was given a descriptive label (e.g., *“press conference”*). Including the two introductory cartoon clips, 50 scenes were analysed in total. The timestamps for both the onset and offset of identified scenes were recorded and aligned across all participants. Both the onset and offset are referred to as the boundaries of the respective event. This is a widely used method for event segmentation and has been validated by a data-driven approach (Baldassano et al., 2017). The length of the events ranges from 11 to 180s (Mean ± SD: 57.5 ± 41.7 s). The distribution of event length is visualized in Figure S1, and the duration of each event is presented in **Table S1**. Each subject’s verbal speech was transcribed, segmented, and matched to the events that were recalled from the movie.

##### Situational variables of movie events

For each movie event, several situational variables including both semantic (e.g., *location*) and affective features (e.g. *arousal*) were analyzed together with the subsequent recall of that event in both correlational analyses and mixed-effect modeling (*see validation analysis below*). Firstly, the entire movie was divided into 1000 time segments (mean duration=3.0s, s.d.=2.2s) by a human rater. Each of the 1000 segments was then labeled for variables including arousal (*excitement/engagement/activity level*), music (*whether or not there is music playing*), location (*whether the location is indoor or outdoor*), and valence (*Positive or negative mood*). For subjective rating (i.e., *arousal* and *valence*), assessments were collected from four different raters (*arousal*: Cronbach’s α = 0.75; *valence*: Cronbach’s α = 0.81). In the event-specific analyses, a score was derived for each of the 50 events for each of the variables mentioned by averaging/adding up ratings across time segments. The event-level scores are displayed in **Table S1**. These analyses were performed and data were shared by Chen and colleagues (Chen et al. 2017). We correlated these variables (along with *event duration*) with the mean recall rate in all participants (**Figure S5**) and found that arousal and event duration positively correlated with memory recall (p<0.05). Other situational variables did not show significant correlations with memory (p>0.05).

##### Low-level perceptual features of movie events

Perceptual changes around event boundaries may influence memory formation and event processing in the brain, therefore we extracted perceptual features for each second of the video. We chose to average these features at 1s resolution to match the resolution of boundary data, while also limiting the computation that would be necessary to analyse the video frame-by-frame. Three kinds of perceptual change were estimated based on the second immediately before and after a boundary: (1) *corrs*: visual correlations calculated using IMage Normalized Cross-Correlation (IMNCC) (Nakhmani and Tannenbaum 2013); (2) *dist*: visual distances calculated by using IMage Euclidean distance (IMED) (Wang et al. 2005); (3) *lum*: global lighting changes calculated by first averaging luminance over all pixels, and then taking the absolute values of the differences. We did not find significant correlations of these measures with situational variables of the preceding event, memory performance, or neural pattern similarity across boundaries (all ps>0.05). These measures were further modeled as fixed effects in our mixed-effects model analyses (see below).

##### Event and order memory

For each participant, we first asked whether events were successfully recalled or not, as in the classical subsequent memory paradigm (Brewer et al. 1998; Wagner et al. 1998; Fernández et al. 1999). An event was labeled as ‘remembered’ if any part of the event was described during the recall. ‘Forgotten’ events are the ones that were not mentioned throughout the recall phase.

Secondly, *out-of-order* events were identified as a measure of order memory. Among all remembered events, an event was labelled as *out-of-order* if it was not described immediately after its preceding event in the original movie. For example, if *event 3* is described immediately after *event 1* without mentioning *event 2*, then *event 3* is an *out-of-order* event. By contrast, if a participant described *event 4, 5, 6* sequentially during the recall phase, since *event 5, 6* correctly followed their preceding event, *event 5, 6* were counted as *in-order* events. The first event verbally described in the recall phase was always labelled as ‘not available’ in the order memory analysis since it is not preceded by any event. It was possible that a single scene was mentioned multiple times (in different parts) during the recall, in which case the position of its first recall was used in the event and order memory analyses.

##### Replication dataset

All of the behavioral results were performed and shared by authors (C.K and J.Z) of the original publication (Kurby and Zacks 2018). Although there are both coarse and fine-grained event boundaries available in the dataset, coarse event boundaries were used for our neural pattern similarity analyses to better match the event duration in the discovery dataset. Recognition memory was measured as the percent of correct responses during the forced-choice test. Action recall memory was scored by how many of each type of action was mentioned (e.g., man drinking coffee). This scoring procedure was based on the Action Coding System (ACS: (Schwartz et al. 1991)). Raw order memory was assessed as the number of errors made in the reordering test. To generate the (final) order memory measure, we multiplied each raw order memory measure by −1 and Z-normalized across all participants. This transformation made the measure intuitive (i.e., *larger values indicate the better order memory*).

### 3. fMRI data analysis

#### 3.1 fMRI data acquisition and pre-processing

##### Discovery dataset

fMRI data were acquired using a T2*-weighted EPI sequence on a 3T Siemens Skyra scanner (20-channel head coil; TR 1,500 ms; TE 28 ms; flip angle 64, spatial resolution 3*3*4 mm^3^). Only data from the encoding phase were analysed and reported in the current study.

A standard pre-processing pipeline was followed using FSL (Jenkinson et al. 2012), which includes slice timing correction, motion correction, linear detrending, high-pass filtering (140 s cutoff), co-registration, and affine transformation into 3 mm MNI standard space (Chen et al. 2017). The time series were shifted 3 TRs (4.5 s) to account for the Haemodynamic response function (HRF). Data were z-scored across time at every voxel and a 6 mm smoothing kernel was applied.

All subsequent analyses were performed on the pre-processed voxel-wise BOLD signal in units of functional volume (TR = 1.5 s). Custom MatLab (R2018b, The Mathworks, Natick, MA) and Python (version 3.6) scripts were used for both Region of Interest and parcellation-based searchlight analysis.

##### Replication dataset

Neuroimaging data were acquired with a Siemens Trio 3T scanner. Functional data (i.e., *movie watching data*) were acquired in five runs using a T2* weighted EPI sequence (TR =2000 ms, TE =27 ms) in 35 transverse slices (voxel size=4.0 mm). We re-ran the preprocessing according to the pipeline used in the *discovery dataset*. Subsequent neuroimaging data analyses were performed on the preprocessed voxel-vise BOLD signal for each ROI.

#### 3.2 Region of interest (ROI) selection

The six ROIs used in this study were independently defined by Chen and colleagues in correspondence to the timescale hierarchy of the event segmentation model (Hasson et al. 2015; Baldassano et al. 2017). Early visual and early auditory cortex were functionally defined based on inter-subject correlation during an audio-visual movie and an audio narrative, respectively (Chen et al. 2016; Simony et al. 2016). ROIs for the medial prefrontal cortex (mPFC) and posterior medial cortex (PMC) were taken from the functional atlas derived from resting-state default mode network (https://findlab.stanford.edu/functional_ROIs.html) from FIND lab at Stanford University (Shirer et al. 2012). The hippocampus and posterior parahippocampal gyrus were anatomically defined from the probabilistic Harvard-Oxford Subcortical Structural Atlas (Desikan et al. 2006). Chen and colleagues manually adjust the threshold of around 50% to ensure better anatomical coverage during the visual check.

#### 3.3 Whole-brain parcellation

Alongside the ROI-based analysis, we performed a parcel-based searchlight analysis on the basis of 1000 functionally parcellated cerebral regions (https://github.com/ThomasYeoLab/CBIG/tree/master/stable_projects/brain_parcellation/Schaefer2018_LocalGlobal). The parcellation was based on a gradient-weighted Markov Random Field (gwMRF) model, which integrated local gradient and global similarity approaches (Schaefer et al., 2018). Using both task and resting-state fMRI acquired from 1489 participants, parcels with functional and connectional homogeneity within the cerebral cortex were generated (hippocampus and subcortical regions were not included). In this fashion, each of these biologically meaningful and non-overlapping parcels can be treated in the same way as an independent region similar to an ROI in the following analyses. The parcellation was provided in both volume and surface space, and the volume-based parcellation space was used in our searchlight analyses.

#### 3.4 fMRI-based neural responses to event boundaries

##### 3.4.1 Univariate response

BOLD signals were first averaged for each TR across all voxels in an ROI. Then the time series were z-scored and segmented based on the event annotations mentioned above. We selected the time window of the univariate response analysis based on the shortest event duration. Among all events, the shortest event was 7 volumes (10.5 s), therefore we focused on BOLD signals 6 volumes before and after the event boundaries (i.e., *in total 13 volumes around event boundaries*).

##### 3.4.2 Activation patterns

Voxel-wise BOLD time series from separate events were first extracted based on the onset and offset timestamps derived from the movie. Multivariate patterns of brain activation were generated for each event by averaging across all volumes within this event. To assess the similarity between two neighboring events, the *activation pattern* for each event of interest was correlated with its following event. The resulting Pearson’s correlation coefficient depicted the extent to which similar representational activity patterns were elicited by neighboring scenes. The lower similarity between two events represented a greater change in neural patterns across the event boundary.

##### 3.4.3 Connectivity patterns

Intra-regional *connectivity pattern* analyses were conducted based on a method originally used in rodent electrophysiology studies to quantify the reactivation of sparsely distributed neuron assemblies (Qin et al. 1997; Lansink et al. 2008) and recently used in human fMRI (Tambini and Davachi 2013, 2019; Hermans et al. 2017). For each event within each brain region, Pearson’s correlations were performed on the extracted m*n (volumes*voxels) BOLD-fMRI time series between each of the n voxel time series. This yielded an n-by-n pairwise correlation matrix (containing p values indicating the significance of the Pearson’s correlations), representing the within-region connectivity structure for each scene. For two neighboring events, Pearson’s correlation coefficient of their correlation matrices was calculated to quantify the similarity for *connectivity patterns*. The lower similarity between two *connectivity patterns* represented a greater change in the intra-region connectivity patterns across the event boundary.

##### 3.4.4 Activation/Connectivity pattern similarity within and between movie events

We compared *within-event* and *between-event* activation/connectivity pattern similarities to reveal the effects of event boundaries on neural pattern shifts. First, we generated “*middle-point boundaries*” for within-event neural similarities calculation. Specifically, for each of the fifty original events, an additional *middle-point boundary* was located at the middle points of the corresponding time series. Thus, one original event can be divided into two “half events” with equal duration. This created 100 “half events” defined by both human-annotated boundaries and “*middle-point boundaries*.” Then, we quantified neural similarities of activation/connectivity patterns between two neighbouring “half events.” If two “half events” were segmented by a human-annotated boundary, then the similarity was defined as the “*between-event*” pattern similarity, whereas if the two “half events” were segmented by a *middle-point boundary,* the similarity was defined as the “*within-event*” pattern similarity.

### 3.5 Relationship between neural responses during encoding and subsequent memory

#### Discovery dataset

##### 3.5.1 Remembered and forgotten events comparisons

We first compared our neural pattern similarities (i.e., *activation pattern* similarity and *connectivity pattern* similarity) at the single-subject level explained above for each brain region (ROI or brain parcel). The similarity indices (Pearson’s r between two matrices) for both *activation* and *connectivity patterns* were averaged for the two types of event pairs (*remembered* and *forgotten*) for each participant. If the first event of the pair was retrieved during the recall phase, the event pair was labeled as *remembered*. *Remembered* and *forgotten* event pairs were then compared in two separate *t*-tests for *activity* and *connectivity pattern* transitions (indexed by pattern similarity).

We further examined the relationship between *connectivity pattern* transitions and order memory (i.e., *temporal order of event recall*). More specifically, *connectivity patterns* were averaged for another two types of event pairs (i.e., *In-order* or *Out-of-order*) for each participant. If the second event of the pair was recalled in an incorrect sequential order (e.g., *event 4* was recalled immediately after *event 6*), the event pair was labeled as *Out-of-order. Connectivity pattern* transitions for *In-order* and *Out-of-order* event pairs were then compared with *t*-tests.

##### 3.5.2 Validation analyses of subsequent memory effects

Beyond the paired t-tests between neural similarities of Remembered and Forgotten events, in total six additional statistical tests were performed to further validate reported subsequent memory effects (Detailed methods and results can be found in *Supplementary Materials*). In brief, (1) neighboring event pairs were divided into four categories based on memory for both the first and second event of the event pair (i.e., *both Forgotten (FF), first Forgotten and second Remembered (FR), first remembered and second forgotten (RF), and both remembered (RR)*). We then compared neural similarities across these four categories. (2) To confirm that the subsequent memory effect on pattern similarity was only present for actual event boundaries but not shuffled boundaries, we generated a null distribution of subsequent memory effects using the event boundary permutation analysis. During each permutation, event boundaries were re-located within events to create the same number of pseudo-events. The sequence of memory labels remain unchanged, and subsequent memory effects were qualified based on pseudo-events and corresponding memory labels. We asked whether subsequent memory effects based on the actual boundaries were larger than effects based on shuffled boundaries. (3) Memory labels (i.e., *R and F*) were shuffled across events, then neural pattern similarities were compared between shuffled memory labels instead of real labels. We aggregated p-values of null R-F difference from 1000 permutations as the null distribution. The real R-F difference (i.e., the *p-value of the paired t-test*) was compared to this null distribution of p-values to generate the permutation-based p-value. To account for the number of ROIs involved, the permutation-based p-value was multiplied by 6 to produce the corrected p-value. (4) For each event, we examined whether the likelihood of an event being remembered among all participants correlated with its mean neural pattern similarity with the previous event. (5) We also performed a cross-participant correlation: we asked whether participants who demonstrated better memory showed lower/higher activation/connectivity pattern similarity during movie watching. (6) We further used a mixed-effects model to examine the relationship between neural similarity and memory, considering both participants and events as random effects and incorporating multiple event-specific situational variables (e.g., *event duration, location, music, arousal*) and low-level perceptual features (i.e., *corrs, dist, and lum*).

#### Replication dataset

Subsequent memory for each movie clip was assessed by three different memory measures (i.e., *recognition, recall, and order memory*). Their relationships with event processing during encoding were investigated as a conceptual replication of findings from the *discovery dataset*. We calculated neural pattern similarities using the “*within-movie*” and “*between-movie*” method separately. For the “*within-movie*” method, participant-specific boundaries generated during the segmentation task were used as event boundaries to calculate event-specific fMRI activation patterns and connectivity patterns during movie watching. Similarities of these activation/connectivity patterns were calculated across boundaries and then averaged for each movie clip at the participant level. To probe the memory relevances of these mean similarities, they were correlated with all three memory measures. For the connectivity pattern similarity analyses, we additionally used the “*between-movie*” method to enable meaningful connectivity analyses with enough TRs. Each movie clip was regarded as an event, and the connectivity pattern was estimated within the entire clip. Then connectivity pattern similarities were calculated across different movie clips and correlated with memory measures.

### 3.6 Relationship between hippocampal pattern similarity and event distance

The above analyses focused on neural pattern similarities between two neighboring events. Here, we examined the hippocampal pattern similarities between events with variable distances. Event distance was defined as the number of event boundaries between two events (the event distance between event 1 and event 3 is 2). For each event, we first calculated its *activation* and *connectivity pattern*. Then, we calculated the *activation* and *connectivity pattern* similarity between all possible combinations of event A-B pairs (‘Event A’ is the event that appeared earlier in the temporal sequence, and ‘Event B’ is the one presented later) within all 50 events. Finally, for each participant, and each event distance, two mean similarities for activation and connectivity pattern were calculated separately. Note that the number of available pairs decreases as the distance increases (e.g., events 1-50 are the only event pair with a distance of 49). To ensure a well-powered analysis for every event distance, we only compared event pairs with a distance less than or equal to 40, meaning at least 10 event pairs contributed to the event distance calculation. Analysis of all distances (d ≤ 49) can be found in the **Supplementary Materials**.

Next, we used linear regression to examine the relationship between pattern similarity and event distance. In addition, to investigate how the subsequent memory of the preceding event (event A) modulates the relationship between event distance and pattern similarity, we ran a two-way ANOVA (memory * event distance) using the memory performance (remembered or forgotten) of the preceding event and event distance (range from 1 to 40) as two independent variables. The relationship between memory and event distance was validated with the permutation test, in which memory labels (i.e., R and F) were shuffled randomly 1000 times to generate null comparisons between two kinds of events.

### 4. Statistical analysis

For parametric hypothesis tests involved in the fMRI data analyses, the significance level was set to *p* = 0.05 (two-tailed). For permutation tests, *p*-values were estimated by comparing real results with null distributions generated by shuffling event boundaries or memory labels, and their significance levels were also set to *p* = 0.05 (two-tailed). To account for the multiple comparisons problem that comes with multiple ROIs or brain regions, all reported *p* values in the main text were FDR-corrected (*p*_FDR_) (Genovese et al. 2002) unless otherwise stated (*p*_raw_). Specifically, this means correction was made for six tests in ROI analyses, and 1000 tests for the whole-brain analyses. All significant *p* values were reported together with the effect sizes (Cohen’s d or partial η^2^). The custom modified version of DABEST (https://github.com/ACCLAB/DABEST-python) was used to plot individual data points alongside otstrapping-based resampled distributions of the mean difference between conditions (Ho et al. 2019).

### 5. Data and code availability

ROI data are available at http://datasets.datalad.org/?dir = /workshops/mind-2017/sherlock. Whole-brain neuroimaging data are available at https://dataspace.princeton.edu/jspui/handle/88435/dsp01nz8062179. The replication dataset was stored at The Central Neuroimaging Data Archive (CNDA), Washington University, Saint Louis (https://cnda.wustl.edu/) and can be requested from Dynamic Cognition Laboratory (https://dcl.wustl.edu/people/jzacks/). Custom code used in this study will be publicly available via the Open Science Framework (OSF) (Link: https://osf.io/p68cv/?view_only=483703873dae4cfd8b36e9d6df6b8c92) upon publication. Further requests for scripts should be directed to the corresponding author.

## Results

### Subsequent memory performance measured by spoken recall

We first calculated recall accuracies for each participant. On average, 68.7% (*SD* = 12%, range 48% - 94%) of the 50 events (*Mean* = 34.4 events, *SD* = 6) were retrieved successfully (**Figure 1C**). Among these remembered events, we further defined *in-order* and *out-of-order* events based on whether they were recalled in the correct sequential order. On average, 58.8% (*SD* = 8%, range 40% - 71%) of the remembered events were *in-order* (**Figure 1D**).

### Neural pattern shifts were larger for between-event transitions compared to within-event transitions

Before linking neural pattern shifts to subsequent memory recall, we investigated how event boundaries modulated neural pattern shifts of both activation and connectivity patterns. More specifically, we compared the effect of *between-event* transitions compared to *within-event* transitions on neural pattern shifts (**Figure S6A**). Paired t-tests between *within-event* and *between-event* similarities revealed that in all six ROIs: (1) *within-event* activation similarities were significantly higher than *between-event* activation similarities; (2) higher connectivity pattern similarities were found for *within-event* transitions compared to *between-event* transitions (**Figure S6B; Table S2**). These results suggest that neural patterns are relatively stable within each event but shift significantly across events. In addition, we correlated activation pattern similarities with connectivity pattern similarities across event boundaries in both the discovery and replication datasets. Results revealed reliable region-specific negative correlations between two neural measures of pattern similarities for the early auditory cortex, PMC, and posterior parahippocampal gyrus (pPHG) (**Figure S7; Table S3**).

### Distinct activation pattern-mediated event segmentation is associated with subsequent retrieval success

We quantified neural similarities of event-specific *activation patterns* before and after event boundaries (i.e., two neighbouring events). Specifically, we generated a voxel-wise *activation pattern* per event by averaging over all time points in that event. This time-averaged *activation pattern* of all voxels within an ROI for an event was compared to the pattern of its subsequent event using Pearson’s correlation. A lower Pearson’s r indicates two more separateble activation patterns and thus more distinct neural representations for two distinct events. We investigated whether *activation pattern* similarities relate to memory formation by contrasting the pattern similarities of remembered with forgotten events in six ROIs. That is, pattern similarity between two events was compared to subsequent memory for the first of those events. We found that subsequently remembered events were associated with lower *activation pattern* similarities than subsequently forgotten events in early auditory cortex (*t* = −3.56, *p*_FDR_ = 0.007, Cohen’s d = 0.92, **Figure 3B**), hippocampus (*t* = −3.62, *p*_FDR_ = 0.007, Cohen’s d = 0.92, **Figure 3E**), mPFC (*t* = −2.79, *p*_FDR_ = 0.01, Cohen’s d = 0.80, **Figure 3C**) and pPHG (*t* = −2.85, *p*_FDR_ = 0.01, Cohen’s d = 0.89, **Figure 3F**). This finding suggests that distinct *activation patterns* for two sequential events are beneficial for the memory of the first event in that sequence. Early visual areas (*t* = −1.13, *p*_FDR_ = 0.27, Cohen’s d = 0.35, **Figure 3A**) and PMC (*t* = −1.91, *p*_FDR_ = 0.08, Cohen’s d = 0.65, **Figure 3D**) did not show this marked effect.

**Figure 3.**
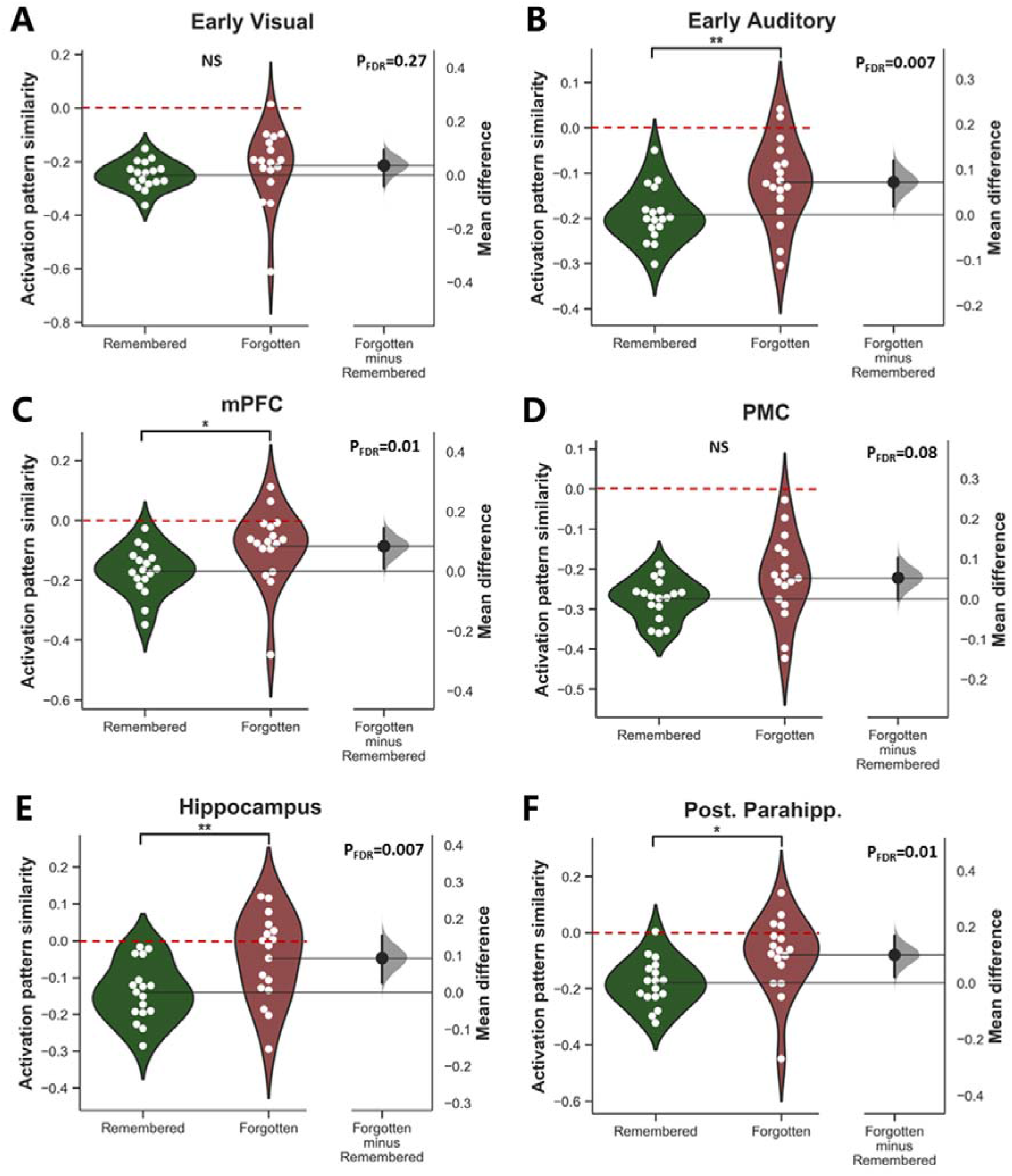
Association between activation pattern similarities of six ROIs and subsequent memory recall. We compared *activation pattern* similarities of sequential event pairs based on subsequent memory performance of the first event (*Remembered* vs. *Forgotten*) across six ROIs. For panel A-F, *activation pattern* similarities for *Remembered* events are displayed on the left (*green*), while similarities for *Forgotten* events are displayed on the right *(red).* For each comparison, a separate axis displays the *mean difference*. The curve (gray) indicates the resampled distribution of the *mean difference* generated via bootstrapping. The solid vertical line attached to the curve represents the *mean difference* as a 95% bootstrap confidence interval. We found significantly lower *activation pattern* similarity for *Remembered* vs. *Forgotten* event pairs in the early auditory area (*t* = −3.56, *p*_FDR_ = 0.007, Cohen’s d = 0.92; panel **B**), mPFC (*t* = −2.79, *p*_FDR_ = 0.01, Cohen’s d = 0.80; panel **C**), hippocampus (*t* = −3.62, *p*_FDR_ = 0.007, Cohen’s d = 0.92; panel **E**), and pPHG (*t* = −2.85, *p*_FDR_ = 0.01, Cohen’s d = 0.89; panel **F**). No significant differences were found in early visual areas (*t* = −1.13, *p*_FDR_ = 0.27, Cohen’s d = 0.35; panel **A**) and PMC (*t* = −1.91, *p*_FDR_ = 0.08, Cohen’s d = 0.65; panel **D**). NS=Not significant; **\*** *p*_FDR_<0.05; **\*\*** *p*_FDR_<0.01.

Beyond the main hypothesis-driven contrasts above (i.e., *paired t-tests*) between activation pattern similarity of remembered and forgotten events, we ran several additional exploratory statistical tests to further examine the relationship between activation pattern similarity and memory. Detailed methods and results from each ROI can be found in the *Supplementary Materials*.

1. In our main analyses above, we labeled an event pair as “*remembered*” if the first event of the pair was remembered. In control analyses, we probed the potential effects of the second event and/or the interaction between the first and second event: event pairs were divided into four categories based on memory (i.e., *both Forgotten (FF), first Forgotten and second Remembered (FR), first remembered and second forgotten (RF), and both remembered (RR))*), and compared neural similarities across these four categories. Consistent with main contrasts, hippocampal activation pattern similarities tended to be lower for RR pairs compared to FF pairs (t=-1.89, *p*_raw_=0.07). Significant effects were found for the early auditory area (t=-2.32, *p*_raw_=0.03), mPFC (t=-3.32, *p*_raw_=0.005), and pPHG (t=-3.36, *p*_raw_=0.004). Full comparations of four categories can be found in **Figure S8**.
2. To test whether the presented results only existed for the actual event structure, we generated shuffled event boundaries and re-ran the same contrasts on activation pattern similarity. Permutation tests demonstrated that presented subsequent memory effects only existed for the actual event boundaries, but not shuffled boundaries (**Figure S9**).
3. The percentage of remembered events was higher than forgotten events, leading to the potential power issue when comparing the two. To counter this, we evaluated the current statistical results by performing a second kind of permutation test: for each permutation, memory labels were shuffled across events, then activation pattern similarities were compared between shuffled memory labels instead of real labels. Similar to our main analyses, permutation tests found that three ROIs (early auditory cortex (*p*_corrected permutation_=0.012), hippocampus (*p*_corrected permutation_=0.006), and posterior parahippocampal gyrus (pPHG) (*p*_corrected permutation_=0.036) showed significant subsequent memory effects after the multiple comparison correction. mPFC demonstrated a significant effect (*p*_permutation raw_=0.014) but did not survive the correction (*p*_corrected permutation_=0.08) (**Figure S10**).
4. So far, within-participant comparisons between remembered and forgotten events revealed that differences in activation pattern similarities of several ROIs were related to subsequent memory. Next, we examined whether a similar relationship was evident across different events. Specifically, we investigated the relationship between the *event-specific recall rate* (the percentage of participants that successfully recalled a particular event) and the averaged *activation pattern* similarity for the corresponding event (the first one in the pair) across all participants. Consistent with our main contrasts, this analysis revealed that the recall rate negatively correlated with *activation pattern* similarity in the hippocampus (r = −0.292, *p*_raw_ = 0.042) and pPHG (r = −0.344, *p*_raw_ = 0.015), suggesting that events showing lower activation pattern similarity with the subsequent event were more likely to be recalled (**Figure S11**).
5. We further performed cross-participant individual differences analysis between activation pattern similarity and memory but found no significant associations. There was a trend for those participants with average lower activation pattern similarity in the early auditory (r=-0.37, *p*_raw_=0.13) and pPHG (r=-0.40, *p*_raw_=0.11) during movie watching performed better at the memory test (**Figure S12; Table S4**), which is consistent with results from our main contrast analyses.
6. Finally, we used a mixed-effects model for statistical analysis to examine the relationship between activation pattern similarity and memory, considering both participants and events as random effects. The relationship between memory and hippocampal activation pattern similarity (F=3.48, *p*_raw_=0.06, R^2^=0.004) failed to reach significance but demonstrated the same tendency as results from the paired t-test. In a second model, event-specific situational variables (e.g., *event duration, location, music…*) were further modeled as fixed effects to be controlled as confounds. Again, hippocampal activation pattern similarity (F=2.80, *p*_raw_=0.09) showed the same tendency of memory effects but failed to reach significance. In a third model, when low-level perceptual features were considered, the relationship between memory and hippocampal activation pattern similarity was significant (F=4.34, *p*_raw_=0.03) (**Table S5**).

### Similar connectivity pattern-mediated event integration is correlated with subsequent retrieval success

Next, we investigated the association between *connectivity patterns* – a different multivariate method to characterise neural representations – and subsequent memory retrieval. Within-region multi-voxel c*onnectivity patterns* were calculated by a voxel-by-voxel pairwise correlation matrix resulting from the correlations between time courses of all voxels within a given region. This represents the relative correlation structure between all voxels in a certain region during event processing. We first calculated the event-specific within-region *connectivity patterns* for two sequential events, and then we quantified the similarity between *connectivity patterns* across event boundaries also using Pearson’s r. Contrasting similarities of *connectivity patterns* of subsequently remembered and forgotten events allowed us to examine how transitions in *connectivity patterns* contribute to memory formation. We found higher *connectivity pattern* similarity for subsequently remembered compared to forgotten events in the early auditory area (*t* = 2.9, *p*_FDR_ = 0.02, Cohen’s d = 0.72, **Figure 4B**), visual areas (*t* = 3.34, *p*_FDR_ = 0.01, Cohen’s d = 0.74, **Figure 4A**), hippocampus (*t* = 3.39, *p*_FDR_ = 0.01, Cohen’s d = 0.73, **Figure 4E**), and PMC (*t* = 2.79, *p*_FDR_ = 0.02, Cohen’s d = 0.47, **Figure 4D**). The same contrast was not significant for mPFC (*t* = 1.22, *p*_FDR_ = 0.23, Cohen’s d = 0.25, **Figure 4C**) and pPHG (*t* = 1.36, *p*_FDR_ = 0.22, Cohen’s d = 0.30, **Figure 4F**).

**Figure 4.**
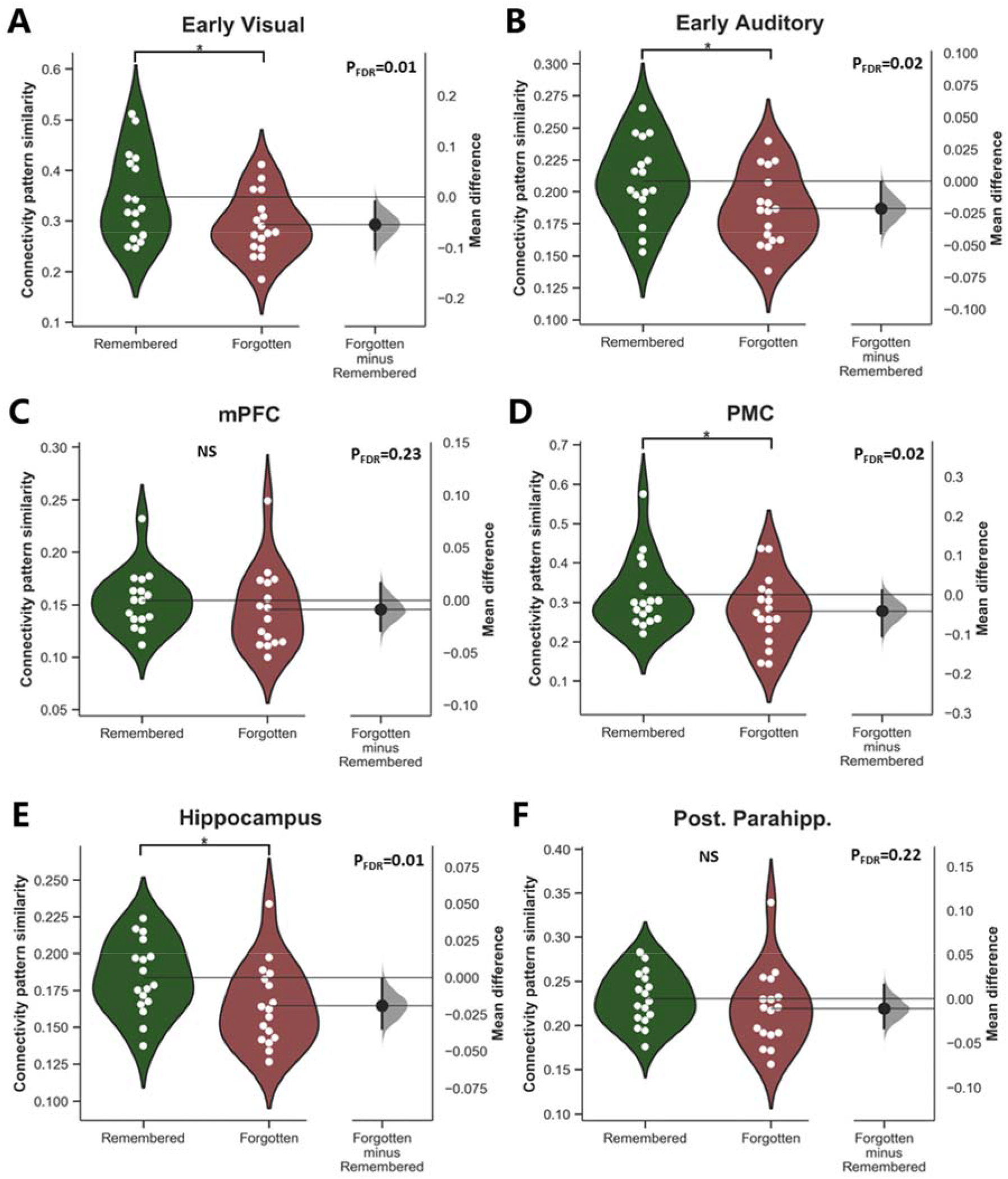
Association between connectivity pattern similarities of six ROIs and subsequent memory recall. We compared connectivity pattern similarities of sequential event pairs based on subsequent memory performance of the first event (*Remembered* vs. *Forgotten*) across six ROIs. For panel A-F, connectivity pattern similarities for *Remembered* events are displayed on the left (*green*), while similarities for *Forgotten* events are displayed on the right *(red).* For each comparison, a separate axis displays the *mean difference*. The curve (*gray*) indicates the resampled distribution of the *mean difference* generated via bootstrapping. The solid vertical line attached to the curve represents the *mean difference* as a 95% bootstrap confidence interval. We found significantly higher connectivity pattern similarity for *Remembered (green)* vs. *Forgotten (red)* event pairs in the early auditory area (*t* = 2.9, *p*_FDR_ = 0.02, Cohen’s d = 0.72, panel **B**), visual areas (*t* = 3.34, *p*_FDR_ = 0.01, Cohen’s d = 0.74, panel **A**), hippocampus (*t* = 3.39, *p*_FDR_ = 0.01, Cohen’s d = 0.73, panel **E**), and PMC (*t* = 2.79, *p*_FDR_ = 0.02, Cohen’s d = 0.47, panel **D**). No significant differences were found in mPFC (*t* = 1.22, *p*_FDR_ = 0.23, Cohen’s d = 0.25, panel **C**) and pPHG (*t* = 1.36, *p*_FDR_ = 0.22, Cohen’s d = 0.30, panel **F**). NS=Not significant; **\*** *p*_FDR_<0.05.

The same set of additional statistical tests was applied to the *connectivity pattern* analyses. (1) Event pairs were divided into four categories based on memory (i.e., *FF, FR, RF, RR*), and connectivity pattern similarities were compared between these four categories. Consistent with our main analyses, hippocampal connectivity pattern similarities are higher for RR pairs compared to FF pairs (t=3.85, *p*_raw_=0.002). This is also true for early auditory area (t=2.56, *p*_raw_=0.02), early visual area (t=3.70, *p*_raw_=0.002) and PMC (t=2.11, *p*_raw_=0.05) (**Figure S13**). (2) Permutation tests examined the specificity of subsequent memory effects to actual event boundaries (as opposed to randomly generated pseudo boundaries) (**Figure S14**). (3) When memory labels were assigned randomly to connectivity patterns, the reported subsequent memory effects disappeared. Permutation tests confirmed subsequent memory effects in four ROIs (early auditory cortex (*p*_corrected permutation_=0.012), hippocampus (*p*_corrected permutation_=0.012), PMC (*p*_corrected permutation_=0.03), and early visual cortex (*p*_corrected permutation_=0.012)) that were very similar to our main analyses (**Figure S15**). (4) The event-specific correlational analysis demonstrated that the recall rate positively correlated with *connectivity pattern* similarity in the early auditory area (r = 0.327, *p*_raw_ = 0.022), visual areas (r = 0.35, *p*_raw_ = 0.014), hippocampus (r = 0.301, *p*_raw_ = 0.036), PMC (r = 0.341, *p*_raw_ = 0.017), and pPHG (r = 0.341, *p*_raw_ = 0.017) (**Figure S16**). This supports our main findings, suggesting that events with higher connectivity pattern similarity with the subsequent event in these ROIs were more likely to be recalled. (5) Individual difference analyses revealed the same trends in the same direction as within-subject the within-subject contrast analyses: participants with higher connectivity pattern similarity in the early auditory (r=0.45, *p*_raw_=0.06) and hippocampus (r=0.40, *p*_raw_=0.10) were more likely to perform better at the memory test (**Figure S12**). (6) Mixed-effects models without (F=3.68, *p*_raw_=0.05, R^2^=0.003), with situational covariates (F=2.81, *p*_raw_=0.09), and with perceptual covariates (F=3.81, *p*_raw_=0.05) showed the same tendency that higher hippocampal pattern connectivity was associated with better memory (**Table S6**).

### Similar connectivity pattern-mediated event integration preserves sequential memory of events in later retrieval

So far, we have shown the opposite association between our two multivariate neural pattern measures and subsequent memory performance: distinct *activation patterns* but similar within-region *connectivity patterns* across events in the early auditory cortex and hippocampus predict retrieval success. This pattern of results suggests that the *connectivity pattern* may integrate events into a continuous sequence. To directly test this hypothesis, we examined the relationship between *connectivity pattern* similarity and sequential order of subsequent recall. We reasoned that if the *connectivity patterns* remain stable across event boundaries, events should tend to be recalled in the correct sequential order. We compared the mean *connectivity pattern* similarities for *in-order* and *out-of-order* events. Controlling for multiple comparisons, we found that *connectivity pattern* similarity in early visual cortex to be larger for *in-orde*r compared to *out-of-order* events (*t* = 3.16, *p*_FDR_ = 0.03, Cohen’s d = 0.47, **Figure 5A**). Similar trends that did not survive correction for multiple comparisons were detected in the hippocampus (*t* = −2.43, *p*_raw_ = 0.026, *p*_FDR_ = 0.08, Cohen’s d = 0.53, **Figure 5E**), auditory area (*t* = −2.08, *p*_raw_ = 0.053, *p*_FDR_ = 0.084, Cohen’s d = 0.46, **Figure 5B**) and posterior parahippocampal gyrus (*t* = −2.05, *p*_raw_ = 0.056, *p*_FDR_ = 0.084, Cohen’s d = 0.36, **Figure 5F**). No such effect was observed in the mPFC (*t* = −1.35, *p*_FDR_ = 0.19, Cohen’s d = 0.19, **Figure 5C**), and PMC (*t* = −2.05, *p*_FDR_ = 0.12, Cohen’s d = 0.33, **Figure 5D**). It is worth mentioning that our method cannot completely disentangle the neural effects of “*event memory*” and “*order memory*”. When we further fully control for the “*event memory*” by restricting analyses on two neighboring events that were both recalled, we did not find different connectivity pattern similarity levels between “*in-order*” and “*out-of-order*” events among six ROIs (***Supplemental Materials****-alternative order memory analysis*).

**Figure 5.**
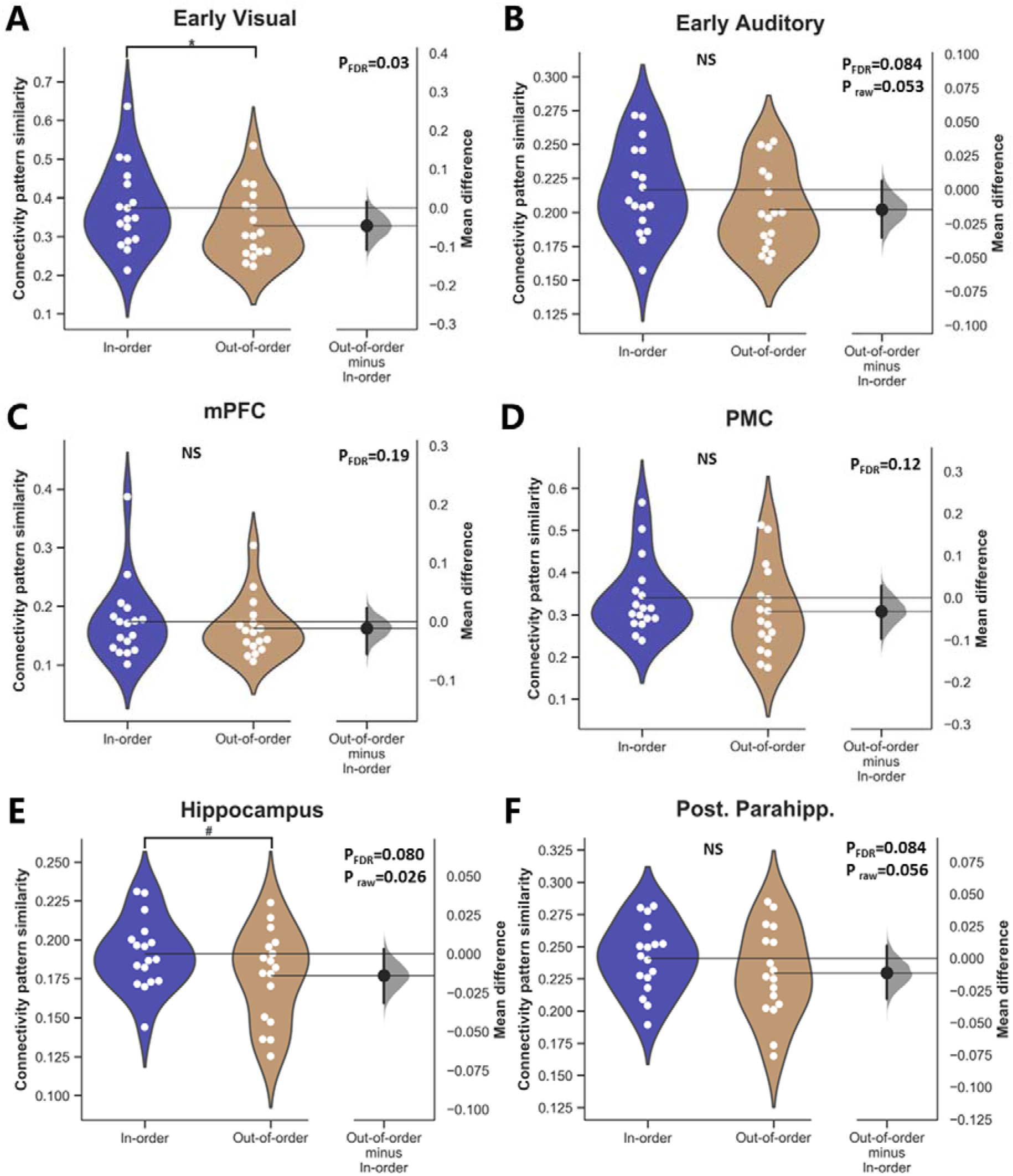
Association between connectivity pattern similarities of six ROIs and sequential order of memory recall. We compared connectivity pattern similarities of sequential event pairs (*In-order* vs. *Out-of-order*) based on sequential memory performance of the first event across six ROIs. For panel A-F, connectivity pattern similarities for *In-order* events are displayed on the left (*BLUE*), while similarities for *Out-of-order* events are displayed on the right *(BROWN)*. Early visual areas (*t* = 3.16, *p*_FDR_ = 0.03, Cohen’s d = 0.47, panel **A**) demonstrated higher connectivity pattern similarity for the *In-order* events compared to *Out-of-order* events. A similar trend was also detected in the hippocampus (*t* = −2.43, *p*_raw_ = 0.026, Cohen’s d = 0.53, panel **E**), but it did not survive FDR correction (*p*_FDR_ = 0.08). We also found modest, non-significant trends in the early auditory area (*t* = −2.08, *p*_raw_ = 0.053, *p*_FDR_ = 0.084, Cohen’s d = 0.46, panel **B**) and posterior parahippocampal gyrus (*t* = −2.05, *p*_raw_ = 0.056, *p*_FDR_ = 0.084, Cohen’s d = 0.36, panel **F**). No similar effects were detected in mPFC (*t* = −1.35, *p*_FDR_ = 0.19, Cohen’s d = 0.19, panel **C**), and PMC (*t* = −2.05, *p*_FDR_ = 0.12, Cohen’s d = 0.33, panel **D**). NS=Not significant; **\*** *p*_FDR_<0.05; **#** *p*_raw_<0.05.

### Hippocampal activation and connectivity patterns change differently with event distance

Among our six ROIs, we found converging evidence for a dissociation of event segmentation and integration in the hippocampus: lower *activation pattern* similarity but higher *connectivity pattern* similarity was beneficial for memory formation. Building on these findings, we hypothesized that hippocampal *activation patterns* of neighboring events should be less similar than events that occur far apart. By contrast, hippocampal *connectivity patterns* of close events should be more similar than events with a long interval in between. Thus, we calculated the *activation* and *connectivity pattern* similarity between all possible combinations of event pairs (‘Event A’ and ‘Event B’) within all 50 events (**Figure 6A** and **6D**). For all pairs of events with the same event distance (e.g., separated by four events), we calculated the mean similarity measure for *activation pattern* and *connectivity pattern* separately. This calculation was repeated for all possible event distances. To ensure reliable estimations of pattern similarities, we only present the similarities of distances with at least ten event pairs (d ≤ 40) in the main text. (Complete calculations can be found in **Figure S17**)

**Figure 6.**
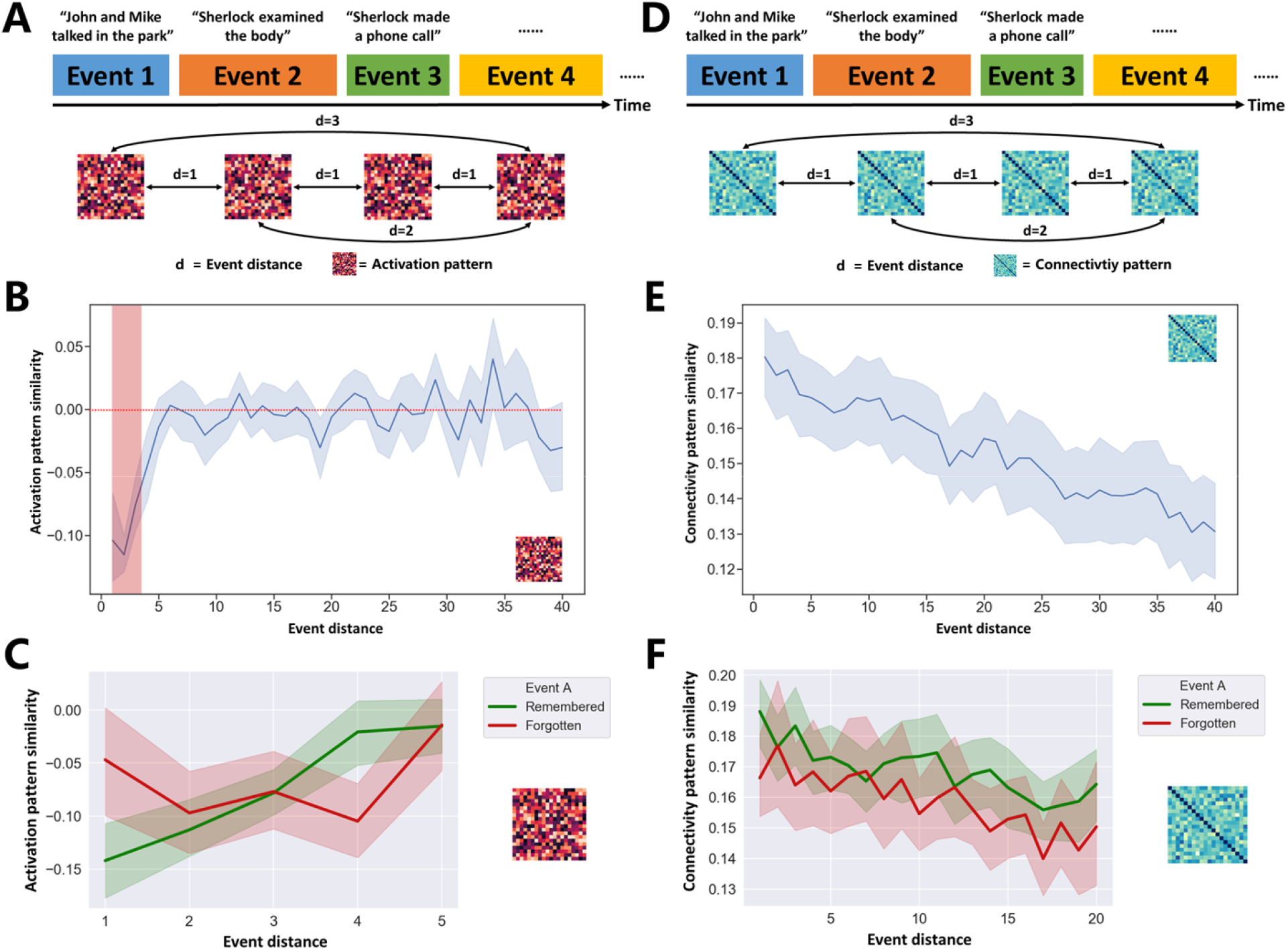
Hippocampal pattern similarity changes with event distance. **(A)** Hippocampal *activation patterns* were generated for all 50 events. We calculated *activation pattern* similarities between sequential events (event distance = 1) and all possible combinations of non-sequential event pairs (event distance > 1). **(B)** Hippocampal *activation patterns* between pairs of events were significantly dissimilar for events separated by a distance of less than 4 (red shadow). **(C)** Memory performance modulated the distance-*activation pattern* similarity relationship. If the first event (*Event A*) of the pair was successfully encoded, *activation pattern* similarities of the event pair increased with event distance (green line). **(D)** Hippocampal *connectivity patterns* were generated for all possible combinations of event pairs. **(E)** Event pairs with the shorter event distance had more similar hippocampus *connectivity patterns*. At the same time, similarities of hippocampus *connectivity patterns* are higher than 0 regardless of event distance. **(F)** Memory performance modulated distance-*connectivity pattern* similarity relationship. If the first event (*Event A*) of the event pair was successfully encoded, *connectivity pattern* similarities of the event pair are enhanced regardless of their event distance. For panel B-F, error bands (i.e., light shadow around the solid line) represent the 95% confidence interval of the mean.

We analysed the hippocampal *activation* and *connectivity patterns* separately. First, our *activation pattern* analysis found that the shorter the event distance, the more distinct the hippocampal *activation patterns* (r = 0.21, *p*_raw_ = 1.8 × 10^−8^; **Figure 6B** and **S16A**). This positive correlation was largely driven by the negative similarity values between events that occurred close to each other (i.e., *events with a distance smaller than four*). Furthermore, we found that subsequent memory recall of Event A modulated the relationship between event distance (d = 1 - 4) and *activation pattern* similarity (ANOVA with event A × distance interaction: *F* (3,48) = 10.1, *p* < 0.001; **Figure 6C**). That is, hippocampal *activation pattern* similarities increased as the event distance changes from 1 to 4, but only if event A was later recalled (*F*_remembered_ (3,48) = 9.54, *p* < 0.001; *F*_forgotten_ (3,48) = 1.35, *p* = 0.268).

Second, our *connectivity pattern* analysis found that the shorter the event distance, the more similar the hippocampal *connectivity patterns* (r = −0.439, *p*_raw_ = 1.8 × 10^−33^; **Figure 6E** and **S16B**). Furthermore, we found a significant interaction between event A recall and distance (*F* (19, 304) = 2.37, *p* = 0.001), and a significant main effect of event A (*F* (1, 16) = 7.53, *p* = 0.014). That is, if event A was recalled later, its hippocampal connectivity pattern was more similar to any other event in the sequence, compared to when event A was not successfully recalled (**Figure 6F**). This suggests that if *connectivity patterns* between pairs of events are more similar, for both short and long distances, then events are more likely to be successfully encoded.

Several time-dependent artifacts may contribute to signals in hippocampal event distance analyses (e.g., *temporal distance, temporal filtering*). These are unlikely to explain the subsequent memory effects we observed, but we ran several further analyses to limit their influence. First, (1) evaluate the effects of these potential artifacts (i.e., *temporal distance and temporal filtering*) on the event distance analysis (**Figure S19-S20**). We investigated how hippocampal pattern similar change with event distance when the temporal distance between events (i.e., *the number of TRs*) was controlled and when different cutoffs (i.e., *140s, 280s, 420s, 560s, 600s*) for high-pass filtering were applied to the time-series; Second, we performed a permutation test to validate the subsequent memory effects in the event distance analysis. We shuffled memory labels (i.e., *R and F*) randomly and performed the event distant analysis for each permutation. Third, event distance analysis was also applied to ROIs beyond the hippocampus to probe whether presented effects are hippocampal-specific. Results can be found in **Figure S21-S22**. These control analyses together demonstrated that (1) main findings were robust to artifacts, (2) relationship between event distance and neural similarity was indeed modulated by memory, (3) similar relationship between event distance and neural similarity were presented in other ROIs as well, but it interacted with subsequent memory in a region-specific manner.

### Subregions of the prefrontal cortex perform event segmentation and integration

Our hypothesis-driven ROI analyses found that (1) distinct hippocampal *activation patterns* were associated with better event memory; (2) similar hippocampal *connectivity patterns* were beneficial for event memory; (3) although not surviving multiple comparison correction, similar hippocampal *connectivity patterns* tended to preserve the sequential order of events (**Figure 7A**). To investigate whether these relationships are present in other brain regions beyond our six ROIs, we ran an exploratory region-based searchlight version of our pattern similarity analysis to identify overlapping event segmentation and integration computations across neocortical regions. In sum, we investigated three potential relationships between neural pattern similarity and subsequent retrieval separately. First, we identified brain regions whose lower *activation pattern* similarities across events were associated with retrieval success (**Figure S23A**). Next, we mapped the association between higher *connectivity pattern* similarities and retrieval success in each region (**Figure S23B**). Then, we identified the regions, which demonstrated a positive association between *connectivity pattern* similarities and order memory (**Figure S23C**).

**Figure 7.**
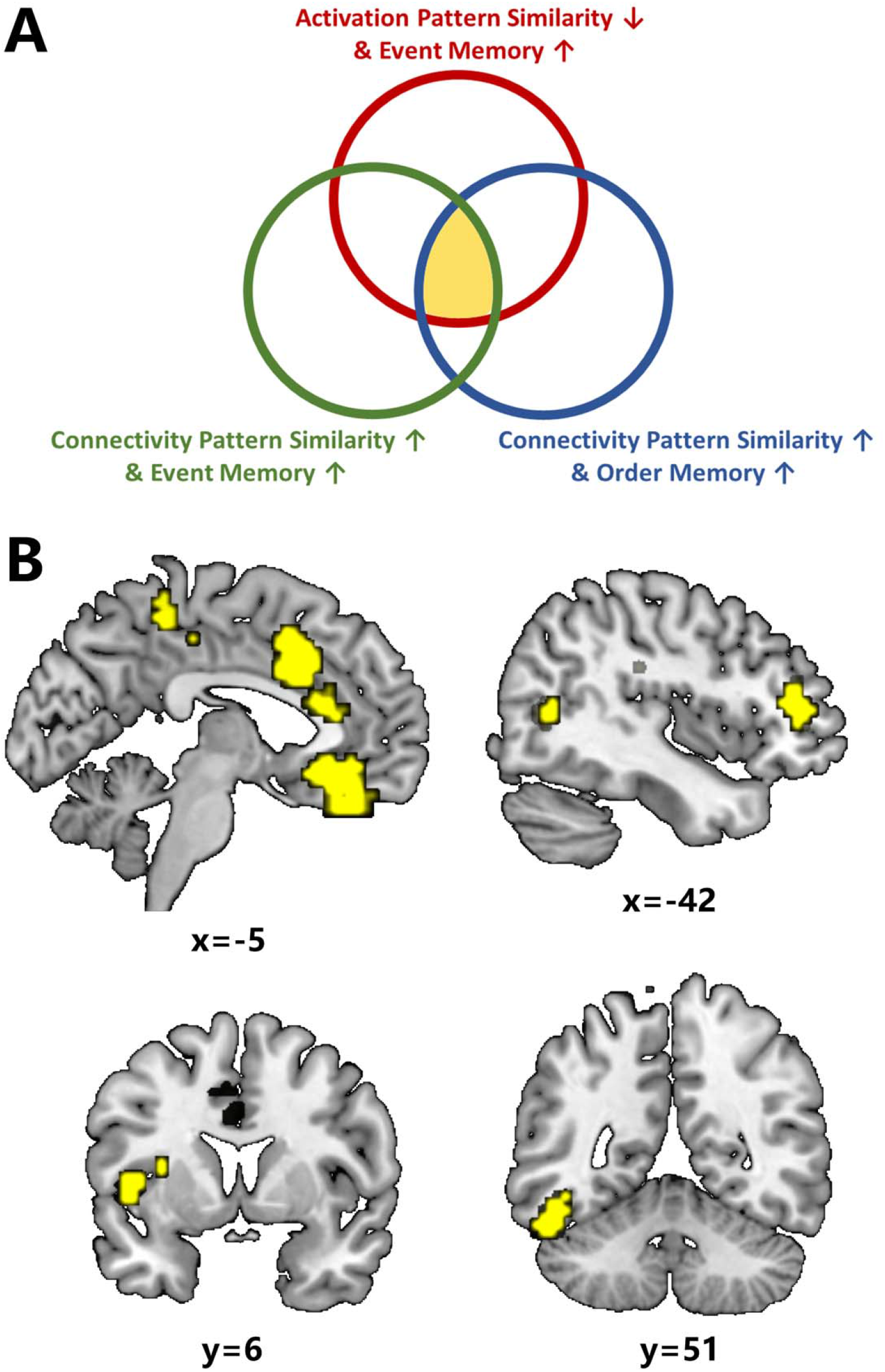
Identifying overlapping event segmentation and integration computations across the neocortex. **(A)** We identified three relationships between neural pattern similarity and subsequent memory in the hippocampus. **(B)** Similar to the hippocampus, overlapping event segmentation and integration computations were found in a network of brain regions including the medial prefrontal cortex (mPFC), right inferior frontal gyrus (IFG), anterior/middle cingulate cortex and supplementary motor area (SMA), left inferior temporal gyrus (ITG) and left insula (*p*_FDR_ < 0.05 across 1000 parcels, cluster size >= 50).

To identify brain regions that may support all three neural computations (*like the hippocampus*), we overlapped spatial patterns for these three effects (all *p*_FDR_ < 0.05). This revealed a set of brain regions including relatively large clusters (at least 50 voxels) in the mPFC, right inferior frontal gyrus (IFG), anterior/middle cingulate cortex and supplementary motor area (SMA), left inferior temporal gyrus (ITG), and left insula (**Figure 7B**). These results suggest that this network of cortical regions may use the same neural processes to perform event segmentation and integration as the hippocampus during continuous memory encoding.

### Hippocampal neural pattern similarities were correlated with memory formation in a replication dataset

We analyzed an independent movie watching-recall dataset (Kurby and Zacks 2018) (i.e., *replication dataset*) to conceptually replicate reported associations between neural similarity and memory formation in the *discovery dataset*. The discovery dataset and replication dataset differed in several aspects of data acquisition protocol (*See Methods*), and therefore, provide the opportunity to test generalization. Participants watched five short movies inside the MRI scanner and returned to the lab several days (mean=3.4 days) later for behavioral testing. During the behavioral session, participants also performed the segmentation task to generate participant-specific event boundaries for each movie clip (**Figure 8A**). These participant-specific boundaries were used as onsets/offsets of events to calculate event-specific fMRI activation patterns and connectivity patterns, and then neural similarities across boundaries within each movie clip (i.e., *“within-movie” method*). The estimation of neural similarities using the “*within-movie*” method is identical to the counterpart used in the *discovery dataset*. Furthermore, neural similarities were also calculated across different movie clips (i.e., “*between-movie*” method) (**Figure 8B**). It is notable that the *“within-movie”* method was used as the hypothesis-testing analysis, and then the “*between-movie*” method was used as the exploratory analysis.

**Figure 8.**
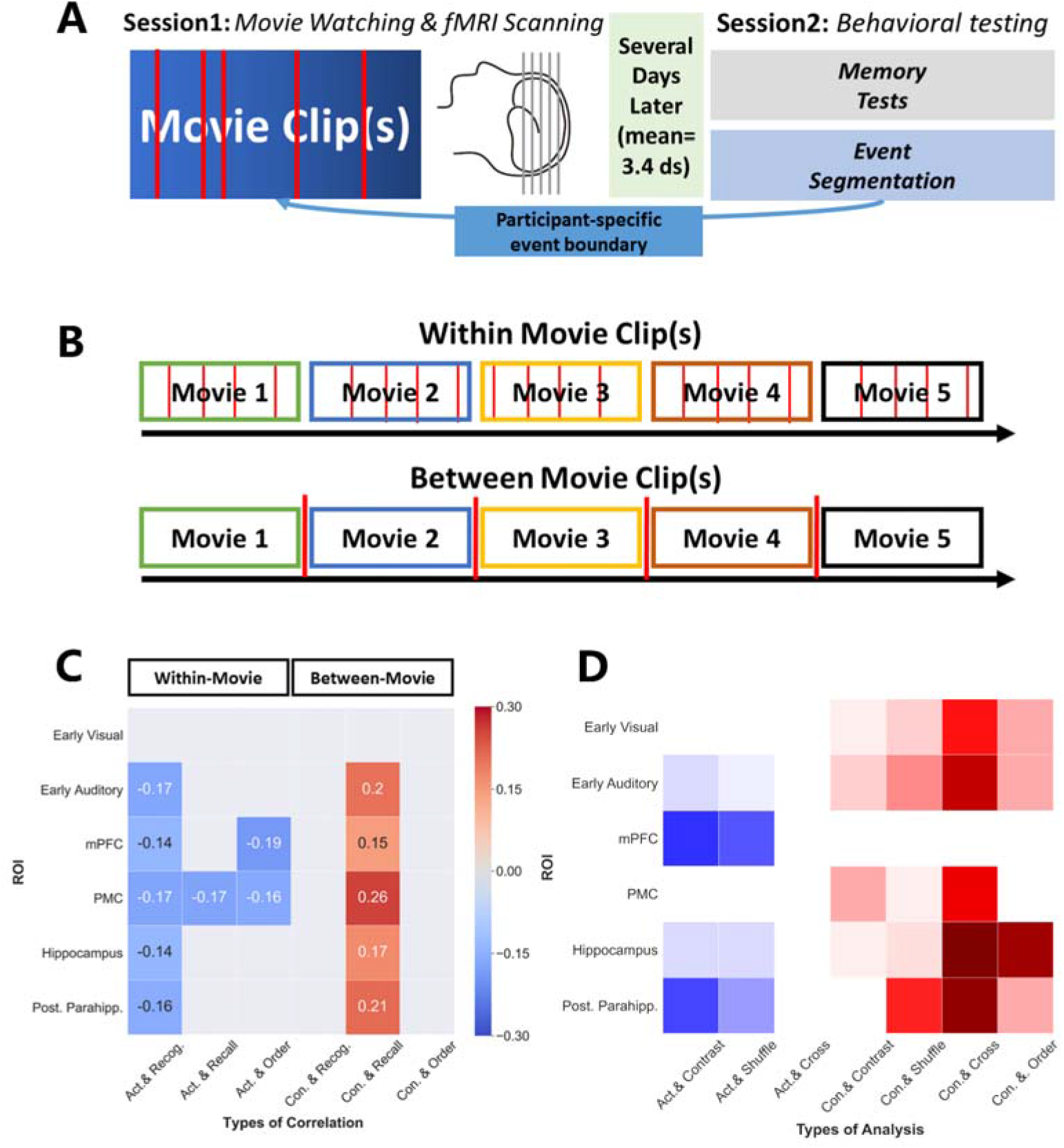
Experimental design as well as data analyses in the replication dataset and comparison between results from two datasets. **(A)** In the replication dataset, participants watched movie clips inside the MRI scanner (session1) and returned to the lab several days (mean=3.4 days) later for behavioral testing (session2), including both segmentation and memory tests. **(B)** Two methods (i.e., *within-movie and between-movie*) were used to estimate neural pattern similarity. For the within-movie method, participant-specific boundaries were used to define events within movies, while for the between-movie method, each movie clip was regarded as an individual event. **(C)** Correlations between ROI-specific neural pattern similarity and memory measures in the replication dataset. Correlation coefficients are presented for significant tests that survived FDR-correction. Full results from the replication dataset can be found in **Figure S24-S25** and **Table S7-S8**. **(D)** Significant tests in the discovery dataset, with solid blocks for those that survived FDR-correction. Colors represent directions of effects (blue=negative association; red=positive association). *Act.=Activation pattern; Con.=Connectivtiy pattern; Recog.=Recognition; Contrast=contrast between Remembered and Forgotten; Shuffle=test with shuffled event boundaries; Cross=cross-event analyses*.

We correlated clip-specific neural similarity measures with all three different memory measures (i.e., *recognition, recall, and order memory*). Results from the replication dataset that survived FDR-correction are summarized in **Figure 8C** alongside the discovery dataset results in **Figure 8D** (for full results, see **Figure S24-S25** and **Table S7-S8**).

Using the “*within-movie*” method, we found that lower activation pattern similarity in the hippocampus during movie watching correlated with better recognition memory (r=-0.14, *p*_raw_=0.028, *p*_FDR_=0.033). Similar correlations were found in early auditory cortex (r=-0.17, *p*_raw_=0.007, *p*_FDR_=0.021), mPFC (r=-0.14, *p*_raw_=0.025, *p*_FDR_=0.033), PMC (r=-0.17, *p*_raw_=0.006, *p*_FDR_=0.021), and pPHG (r=-0.16, *p*_raw_=0.01, *p*_FDR_=0.022). Thus, the relationship between subsequent retrieval success and lower activation pattern similarity in the hippocampus as well as early auditory cortex, mPFC, and pPHG was replicated across our two datasets (**Figure 8C-D**).

Activation pattern similarity in the mPFC and PMC also correlated with additional memory measures (mPFC: with order memory (r=-0.19, *p*_raw_=0.002, *p*_FDR_=0.012); PMC: with order memory (r=-0.16, *p*_raw_=0.009, *p*_FDR_=0.027) and recall (r=-0.17, *p*_raw_=0.006, *p*_FDR_=0.036)). However, connectivity pattern similarities estimated by the “*within-movie*” method did not associate with any memory measures. This may because durations were too short based on participants’ segmentation: event durations were around 60s in the *discovery dataset,* compared with 6-10s in the *replication dataset*. We propose that 3-5 TRs are insufficient to be viewed as “*encoding context*” and enable meaningful connectivity analyses. To explore further, we next regarded each movie clip (2 to 6 min) as an event and performed the calculation of connectivity patterns using the “*between-movie*” method. We found that higher connectivity pattern similarity across movie clips in the hippocampus (r=0.17, *p*_raw_=0.013, *p*_FDR_=0.019), early auditory cortex (r=0.2, *p*_raw_=0.003, *p*_FDR_=0.006), mPFC (r=0.15, *p*_raw_=0.036, *p*_FDR_=0.043), PMC (r=0.26, *p*_raw_<0.001, *p*_FDR_<0.006), and pPHG (r=0.21, *p*_raw_=0.002, *p*_FDR_=0.006) was associated with better recall performance. Therefore, the “*between-movie*” method showed that the relationship between retrieval success and higher connectivity pattern similarity in the hippocampus, PMC, and early auditory cortex was also replicated across two datasets (**Figure 8C**-**8D**).

Furthermore, we found modest evidence (i.e., *p*_raw_ value around 0.05) to support that higher connectivity pattern similarity in these ROIs tended to associated with better order memory performance (early auditory cortex (r=-0.137, *p*_raw_=0.048), mPFC (r=-0.12, *p*_raw_=0.08), PMC (r=-0.135, *p*_raw_=0.052) and pPHG (r=-0.12, *p*_raw_=0.08)). Across two datasets, connectivity pattern similarities of the early auditory cortex and pPHG were associated with the better order memory.

In sum, the relationships between hippocampal activation and connectivity pattern similarity and episodic memory formation were replicated across two datasets. However, higher hippocampal connectivity pattern similarity was associated with better order memory only in the *discovery dataset* but not in the *replication dataset*. Also, our “*within-movie*” connectivity analysis did not reveal the subsequent memory effect, but that could be due to experimental design (i.e., short events). When we analyzed connectivity patterns on a longer timescale with “*between-movie*” analysis, then memory effects got replicated.

## Discussion

To successfully form memories of our life experiences, we need to segregate continuous experience into events (Baldassano et al. 2017; Zacks 2020) and integrate those events across their boundaries into a coherent narrative (Griffiths and Fuentemilla 2020). Here we show that distinct hippocampal *activation patterns*, but similar hippocampal *connectivity patterns* across event boundaries, facilitate these two vital episodic memory functions. We propose that distinct *activation patterns* reflect event segmentation while similar *connectivity patterns* integrate separately represented events into a narrative. Supporting this role of *connectivity patterns* for event integration, we found that similar hippocampal *connectivity patterns* were relevant for the correct sequential order of subsequent retrieval. Our whole-brain analysis demonstrates that similar neurocomputations were performed by a network of cortical regions, in particular for the mPFC. Finally, these results were further validated using an independent movie-watching recall dataset, in which different stimuli and memory measures were utilized. Among the same set of ROIs, hippocampal activation and connectivity pattern similarity showed consistent relationships with memory across these two datasets. Overall, these results suggest that both hippocampal and medial prefrontal event segmentation and integration support memory formation of continuous experience.

Using multivoxel pattern analysis, we found that distinct local *activation patterns* across event boundaries in the early auditory area, mPFC, posterior parahippocampal gyrus, and hippocampus were associated with better subsequent memory, indexed by more distinct activation patterns between two adjacent events. The ability to segment continuous experience at the behavioral level has been linked to successful memory encoding (Sargent et al. 2013), and compelling evidence suggested that the hippocampus is activated around event boundaries (Ben-Yakov and Dudai 2011; Ben-Yakov et al. 2013; DuBrow and Davachi 2013; Ben-Yakov and Henson 2018). This hippocampal activity has been proposed to be associated with a hippocampal segmentation process, but how the hippocampus represents two separate events and whether the corresponding neural representations are relevant for memory remained unclear. Our findings suggest that the hippocampus and other brain regions (e.g., mPFC) segment events by representing them with two distinct patterns of activity. This is similar to the role of the hippocampus in pattern separation: when similar experiences need to be discriminated and encoded, the underlying hippocampal neural representations tend to be dissimilar (Bakker et al. 2008; Yassa and Stark 2011). This neural phenomenon has typically been studied to show how the brain separates perceptually similar stimuli (i.e., images). Here, our findings indicate that a similar pattern separation occurs for events during the continuous experience, and this determines subsequent memory for those events. That is to say, the episodic memory system may use ‘*orthogonalized’* neural representations to encode two events for the purpose of event segmentation. Further, we show these ‘*orthogonalized’* neural representations are potentially event-distance dependent: the hippocampus only generates consecutive dissimilar patterns when events occur relatively close in time. Taken together, this suggests the existence of a brain network (mainly hippocampus and mPFC) for the continuous segmentation of ongoing experience, and the degree of separate neural event representation for nearby events is relevant for memory formation.

Complementing this, we found that more similar within-region *connectivity patterns* of several regions across event boundaries, including again the early auditory area and hippocampus, were associated with better subsequent recall. Compared to local *activation patterns* (Cohen et al. 2017; Xue 2018), within-region *connectivity patterns* are a less used multivariate approach. Recently, Tambini and Davachi proposed that both *activation* and *connectivity patterns* could be used to capture neural states during memory encoding and reactivation, but *connectivity patterns* tend to encode contexts or states instead of particular perceptual inputs (Tambini and Davachi 2019). Our results support this notion: *activation patterns* were event-specific (Chen et al. 2017) and dissimilar for neighboring events, while *connectivity patterns* were relatively stable across event boundaries. Importantly, the stability of connectivity patterns positively associated with memory formation. Therefore, the function of similar *connectivity patterns* could be to integrate segmented and separately represented events into a coherent narrative. To provide further support for this idea, we examined the relationship between connectivity pattern similarity and sequential order during memory recall and found that higher cross-event connectivity pattern similarity (mainly in the hippocampus and pPHG) was associated with better retrieval of the sequential order of individual events. Results from the replication dataset, in which a more accurate order memory measure was adopted, still demonstrated the role of pPHG’s connectivity pattern in promoting order memory. In general, these results revealed that the multi-voxel *connectivity pattern* could be used to predict how temporal sequences are represented in the human medial temporal lobe memory system. Interestingly, our activation and connectivity pattern measures were both modulated by event boundaries in a similar way (ie., *greater within-event similarity compared with between-event similarity*). But the way they interact with memory was different/opposite, with dissimilar patterns being better for activation, and similar patterns being better for connectivity. In addition, contrary to our prediction, lower activation pattern similarities were associated with higher connectivity pattern similarities in certain ROIs across both discovery and replication dataset. This may suggest that, at least in the current dataset, there is no trade-off between event segmentation and integration. Instead, strong event segmentation and event integration appear to happen simultaneously. The local connectivity pattern has rarely been measured in human fMRI studies, therefore its precise cognitive function may reach beyond the representation of temporal context that we are proposing. Also, the question of whether activation- and connectivity patterns play complementary roles in other mental operations should be investigated in future studies.

Remembering the sequence of events is not only one of the critical features of episodic memory (Davachi and DuBrow 2015) but also highly relevant for other forms of sequence learning, for example, spatial memory and encoding of temporal information (Eichenbaum 2014; Bellmund et al. 2020). Animal studies have revealed the existence of hippocampal neurons that potentially encode temporal context via a gradual change in its neural activity patterns (Manns et al. 2007). More recently, *event-specific rate remapping cells* (Sun et al. 2020)) in the hippocampus were shown to be causally involved in representing temporally structured experience. Single-cell recordings in the hippocampus of patients with pharmacologically intractable epilepsy showed that as a result of repeated viewing of the same video clips, neuronal activity in successive time segments became gradually correlated, and this potential measure of temporal binding predicted subsequent recall (Paz et al. 2010). This study revealed how the temporal relationship between current hippocampal activity and hippocampal activity that follows in time could be used to link successive events in humans. Other recent human fMRI studies revealed the role of the human hippocampal-entorhinal region in representing the temporal sequence of experience across different paradigms and stimuli (Lositsky et al. 2016; Bellmund et al. 2019; Montchal et al. 2019; Thavabalasingam et al. 2019). Adding to this, our study tested the role of a new multivariate measure, *connectivity pattern* similarity, which could reflect the internal stability of neural states in temporal sequence coding. We provided preliminary evidence that *connectivity pattern* similarity across event boundaries in the medial temporal lobe is involved in sequential memory. Future studies are needed to further investigate the precise mnemonic functions of different neural measures (e.g., *activity pattern*, within-region *connectivity pattern*, and system-level interaction between regions) during memory formation, in particular for encoding temporal structure during the continuous experience (Tambini and Davachi 2019).

Our ROI analysis highlights the two functions of the hippocampus in the separate representation of segmented events and the binding function that linked events into a narrative, and region-based searchlight analysis identified the role of subregions of the prefrontal cortex (e.g., mPFC, IFG), insula, and inferior temporal gyrus in event segmentation and integration during memory formation. The role of the mPFC in event integration is particularly thought-provoking. The mPFC is generally implicated in encoding and retrieval of episodic memories (Kim 2010; Rugg and Vilberg 2013). Among its variety of functions in learning and memory (Fernández 2017), the online integration of events we observed here is consistent with its function in the facilitation of associative inference (Zeithamova et al. 2012; Preston and Eichenbaum 2013; Schlichting et al. 2014; Schlichting and Preston 2015; Spalding et al. 2018), accumulation of knowledge (Kumaran et al. 2009; Berkers et al. 2018), and integration of new and prior knowledge (van Kesteren et al. 2010, 2013, 2014). We propose that the general mnemonic function of mPFC is to establish links between separate elements across time and space. Taken together, we found that the hippocampus-mPFC circuit performs event segmentation and integration during memory formation of continuous experience. These findings demonstrate the contribution of two complementary event processing mechanisms and underlying neural representations in episodic memory formation. The hierarchical network model of event segmentation proposes that higher-order regions receive event representations from lower-order perceptual regions, and then transfer these representations to the hippocampus for storage (Hasson et al. 2015; Baldassano et al. 2017). Our study suggested that event integration is another key cognitive process involved in event memory by showing how distinct event representations are integrated by similar *connectivity patterns* of hippocampus and mPFC.

Beyond the hippocampus, the role of sensory regions in processing incoming continuous real-life-like experiences was also predicted by the hierarchical theory of event processing (Hasson et al. 2015) and supported by initial empirical evidence (Baldassano et al. 2017) and our analyses. Using a data-driven method, Baldassano and colleagues showed that event segmentation signals could exist in multiple brain regions that including different levels of sensory regions. Interestingly, continuous experience was segmented at different time scales by regions in the hierarchical structure (Baldassano et al. 2017). The high-level regions (e.g., *angular gyrus and PMC*) may represent the event structure that aligns best to human-labels, while lower-level sensory regions not only respond to this coarse segmentation but also create fine-scale segmentations within the individual event. Therefore, it is perhaps not surprising that, when we used human-labeled boundaries to analyses signals from sensory regions, we still observed associations between event segmentation measures and subsequent memory.

Our study, together with previous studies also combining human fMRI with naturalistic stimuli (Hasson et al. 2008; Baldassano et al. 2017; Chen et al. 2017), demonstrates the potential of this approach to advance our understanding of the human memory system, in particular for the formation of real-life memories. Similar paradigms and analyses can be easily adapted in clinical (e.g., memory and affective disorders) and developmental neuroimaging studies (e.g., children and older adults) to reveal changes related to disease or (mal)development. For example, fMRI-based event segmentation and integration measures could be used to probe how these processes are impaired in Alzheimer’s disease and mild cognitive impairment, how they develop from childhood to adulthood and diminish in normal aging. In addition, *connectivity patterns* have the potential to inform our understanding of other cognitive operations that require the integration of information, such as inferential reasoning (Preston and Eichenbaum 2013). However, due to the low temporal resolution of fMRI, the directionality of information flow between the neocortical regions of the ‘hierarchical memory system’ (Hasson et al. 2015) and the hippocampus remains unclear. Future application of deep-source magnetoencephalography (MEG) (e.g., Backus, Schoffelen, Szebényi, Hanslmayr, & Doeller, 2016) or intracranial electroencephalography (iEEG) (e.g., Jafarpour, Griffin, Lin, & Knight, 2019) with naturalistic memory paradigms may bridge this gap.

Despite the mentioned advantages of combining neuroimaging, naturalistic stimuli, and subsequent memory design to study episodic memory formation, there are also disadvantages worth noting, especially concerning the *discovery dataset*. For example, all participants viewed the same movie in the same order, therefore, memory for individual event and their sequential relationship cannot be isolated. Furthermore, in the discovery dataset, memory performance was relatively high, which means subsequent memory contrasts in some participants may not be well-powered. The inclusion of the replication dataset could potentially mitigate these concerns, revealing rather consistent neural effects in the hippocampus and parahippocampus across two datasets. Another disadvantage of naturalistic stimuli is that situational variables (e.g., *duration, music, location*) and low-level perceptual features (e.g., *visual similarity, luminance*) could influence how memorable each event is. Some of them were indeed positively correlated with memory (e.g., *event duration and arousal*). We investigated the effects of situational and low-level perceptual factors by including them in our mixed model analysis. After controlling for these confounds, hippocampal effects remained, although some effects were reduced to trend level (i.e., *p-values around the 0.05 threshold*). This pattern of results suggests that other factors beyond the hippocampal event segmentation and integration may also play an important role in forming continuous experience. But, in general, the results were consistent with our main analyses. Therefore, our proposed mnemonic functions of the hippocampus are not likely to be the result of external factors. Besides, it remains important to note that our key findings were replicated across two independent and distinct datasets, further strengthening our main conclusions.

In sum, we show that the hippocampus and mPFC may perform a dual function during naturalistic memory formation. Both regions segment events by representing them with distinct *activation patterns*, while also integrating those events by retaining similar *connectivity patterns* across events, enabling the representation of a coherent narrative. The ability to measure segmentation- and integration-related neural operations using fMRI opens new opportunities to investigate the mechanisms of memory encoding for real-life experience.

## Acknowledgment

We thank Dr. Janice Chen for sharing the dataset and ROI masks and suggestions on the searchlight analysis of this manuscript; Kirsten Rittershofer for proofreading this manuscript; Members of the Hasson and Norman laboratories for their efforts during data acquisition and data sharing. This work was supported by the Chinese Scholarship Council (PhD fellowship (201606990020)) to W.L), Ministerie van OCW (Sino-Dutch Bilateral Exchange Scholarship to YJS), and Nuffic Neso China (Orange Tulip Scholarship to YJS).

