## Supplementary Material for "Hippocampal-medial prefrontal event segmentation and integration contribute to episodic memory formation"

**This PDF file includes:**

Supplementary Text

Figure S1-S25

Table S1-S8

Correspondence:

Wei Liu

School of Psychology,

Central China Normal University (CCNU),

No. 152 Luoyu Road, Hongshan District, Wuhan 430079,

Hubei Province, Wuhan, China.

**1. Activity time courses across event boundaries are related to subsequent recall**

To identify regions in which the univariate neural responses to event boundaries were related to subsequent memory (i.e., *subsequent memory effect*), we directly compared the univariate activity time courses around boundaries for *remembered* (*R*) with *forgotten* (*F*) events. We were interested in the activity time courses for an event that was associated with successful memory (*R* vs. *F;* **Figure S2**, highlighted in blue shading), but also how the activation changed in the following event. This gave us a 2*2 design with *Memory* (*R* and *F)* and *Event (current* and *next event)* as independent variables (**Figure S3**). For each of the six predefined ROIs (**Figure S1**), BOLD signals around event boundaries (from 6 volumes before to 6 volumes after each boundary) were extracted from the time series during movie watching, labelled (*R* vs. *F*) based on the memory performance for the first of the two events, averaged across all the available boundaries, and transformed into z-scores for each participant.

The interaction between *Memory* and *Event* was significant in early visual cortex (*F* = 13.109, *p*_FDR_ = .006, η² = 0.45; **Figure S2A**), early auditory cortex (*F* = 10.852, *p*_FDR_ = .010, η² = 0.404, **Figure S2B**), hippocampus (*F* = 8.217, *p*_FDR_ = .017, η² = 0.339, **Figure S2E**), and posterior parahippocampal gyrus (pPHG) (*F* = 15.562, *p*_FDR_ = .006, η² = 0.493, **Figure S2F**), but not in mPFC (*F* = 3.036, *p*_FDR_ = .101, η² = 0.160, **Figure S2C**) and PMC (*F* = 4.718, *p*_FDR_ = .054, η² = 0.228; **Figure S2D**). For the current (first) event, we found greater activation for *remembered* compared to *forgotten* events in early visual cortex (*R*: 0.143 ± 0.125 (mean ± SD), *F*: -0.090 ± 0.145 (mean ± SD); *t* = 21.911, *p*_FDR_ < .001), hippocampus (*R*: 0.134 ± 0.079 (mean ± SD), *F*: -0.019±0.185 (mean ± SD); *t* = 11.801, *p*_FDR_ = .006) and pPHG (*R*: 0.118 ± 0.083 (mean ± SD), *F*: -0.028 ± 0.155 (mean ± SD); *t* = 13.264, *p*_FDR_ = .006). This effect was reversed in early auditory cortex for the next event, which showed greater activity for *forgotten* compared to *remembered* events (*R*: -0.217 ± 0.155 (mean ± SD), *F*: -0.034 ± 0.136 (mean ± SD); *t* = 10.803, *p*_FDR_ = .030). Consistent with previous subsequent memory studies^1-3^, enhanced activity in the timeframe of the current event was associated with successful subsequent recall. To further take into account the memory performance of the *next event*, the response curves were split into four (*R-R*; *R-F*; *F-R*; *F-F*) based on the memory performance of two consecutive events for demonstration (**Figure S3**). In sum, these results provided confirmative evidence for the role of hippocampus and pPHG’s univaraite activtiy in memory formation of continous experience and can be regarded as the data quality assessment.

**2. Analyse subsequent memory effects based on event pairs**

As shown in **Figure S7** and **Figure S12**, neighboring event pairs were divided into four categories based on memory (i.e., *both Forgotten (FF), first Forgotten and second Remembered (FR), first remembered and second forgotten (RF), and both remembered (RR))*, and we then compared neural similarities across these four categories. Notably, the number of event pairs within each category differed significantly based on individual memory performance. Therefore, presented contrasts may not be well-powered and that is the reason why some categories may have larger variations than others.

Two main statistical tests were applied to activation pattern similarity and connectivity pattern similarity separately: (1) we used a repeated 2-by-2 ANOVA to analyses potential group differences: first event (R vs. F) * second event (R vs. F). (2) we directly compared groups differences between FF and RR categories (to avoid considering the memory effect of only the first or second event)

Activation pattern similarity:

(1) For the hippocampus: there is no interaction between the factor memory of the first and second event and activation pattern similarity (F=0.169, p=0.68), and no effect of memory of the second event (F=4.351e-4, p=0.984). As already reported in the initial submission, the memory of the first event modulates the activation pattern similarity (F=32.88, *p*_raw_<0.001). The same analysis was applied to ROIs that demonstrated subsequent memory effects in our initial submission and revealed the same statistical patterns: subsequent memory effects were mainly driven by the first event (early auditory area (*p*_interaction_=0.68; *p*_first_=0.98; *p*_second_<0.001); mPFC (*p*_interaction_=0.11; *p*_first_=0.18; *p*_second_<0.001); pPHG (*p*_interaction_=0.65; *p*_first_=0.32; *p*_second_ =0.003)).

(2) Hippocampal activation pattern similarities tend to be lower for both remembered (RR) pairs compared to both Forgotten (FF) pairs (t=-1.89, *p*=0.07). This is also true for the early auditory area (t=-2.32, *p*=0.03), mPFC (t=-3.32, *p*=0.005), and pPHG (t=-3.36, *p*=0.004)

Connectivity pattern similarity:

(1) For the hippocampus: there is no interaction between the factor memory of the first and second event and connectivtiy pattern similarity (F=1.00, p=0.33). We found that both the memory of the first event (F_first_=12.47, *p*_first_=0.003) and the next event (F_second_=5.46, *p*_second_=0.03) modulates the connectivity pattern similarity. Except for the early auditory area (F_first_=9.50, *p*_first_=0.008; F_second_=1.726, F_second_=0.20), early visual area (F_first_=14.48, *p*_first_=0.002; F_seond_=8.78, *p*_second_=0.01) and PMC (F_first_=3.71, *p*_first_=0.07; F_second_=4.13, *p*_second_=0.06) demonstrated the same pattern of statistical results as the hippcocampus.

(2) Hippocampal connectivity pattern similarities are higher for both remembered (RR) pairs compared to both Forgotten (FF) pairs (t=3.85, *p*=0.002).This is also true for early auditory area (t=2.56, *p*=0.02), early visual area (t=3.70, *p*=0.002) and PMC (t=2.11, *p*=0.05)

In sum, by analyzing neural pattern similarity based on event pairs, we showed that (1) for the relationship between activation pattern similarity and memory, the memory effect was mainly driven by the effect of the first event while for the connectivity pattern similarity, subsequent memory effects can be detected for both the first and second event (*but there are no interaction/additional effects*); (2) subsequent memory effects were still evident in the same ROIs if we only compared “both-remembered (RR)” and “both-forgotten (FF)” event pairs. The latter result suggests that our results presented so far are not dependent on a certain way of memory labeling (i.e., l*abeling only the first event*).

**3. Event boundary permutation analysis**

To confirm that the subsequent memory effect on pattern similarity was only present for actual event boundaries (annotated by an independent rater based on the movie narrative), we shuffled the timestamps of the original annotated boundaries and tested whether the simulated boundaries were associated with memory. To retain the event structure (i.e., avoid cases where the pseudo events are too long/short), we held onto the duration of the 50 events and only scrambled their order, which led to randomly placed event boundaries across the sequence with the original set of event durations. Our calculation for *activation* and *connectivity patterns* was then performed on these permutated boundaries, yielding simulated *activation* and *connectivity pattern* similarities.

Paired *t*-tests were performed to test whether the pattern similarity indices were still associated with memory labels. This permutation procedure followed by the *t*-test was repeated 5000 times for both the *activation* and *connectivity patterns*, which gave us a null distribution of *t* and *p* values. The *p* value from the analysis using genuine event boundaries was subsequently corrected against this distribution by calculating the proportion of sampled permutations where the *p* values were smaller than or equal to the real observation. The regions that showed significant differences in *activation pattern* similarities are the same ones as in the previous analysis using real event boundaries: early auditory area (*p* = 0.001), hippocampus (*p* = 0.003), mPFC (*p* = 0.01), and pPHG (*p* = 0.006) (**Figure S8**). For *connectivity patterns*, the SME only exists for actual event boundaries in the early auditory area (*p* = 0.012), visual areas (*p* = 0.005), hippocampus (*p* = 0.005), and PMC (*p* = 0.012) (**Figure S13**). These results confirmed the specificity of the SME to actual event boundaries.

**4. Memory label permutation**

We ran the permutation test on memory labels (i.e. *R versus F*) to validate our main results because of the power issues in subsequent memory analyses (i.e., more R events compared to F events). For each permutation, memory labels were randomly shuffled within each participant, and then neural pattern similarities were calculated for shuffled R and F events separately. This permutation was repeated 1000 times, generating null distributions of two kinds of neural pattern similarities. The null distributions were visualized in the right panels of **Figure S9**, and **Figure S14**: (1) all ROIs showed inseparable distribution between shuffled R and F events in terms of (mean) similarity values (*right panels*) while similarity values of real R and F events differ in an ROI-specific manner (*middle panels*); (2) we indeed found the effects of the unbalanced number of trials between R and F events. But it affected the standard deviation instead of the mean values of the two distributions. Because there were significantly more R events than F events, the estimations of similarities are more stable (i.e., *lower standard deviation*) for R events compared to F events. Therefore, we can see that during permutations, the variances of similarities values for R events were smaller than F events. In sum, the ROI-specific subsequent memory effects can not be observed anymore when shuffled memory labels were analyzed. It suggests that although the power issue is exigent, presented subsequent memory effects are not the results of systematic errors that may be associated with a certain trial/recall structure.

**5. Cross-event correlation and cross-participant correlation**

Thus far we have examined the association between memory and neural pattern similarity in a within-participant fashion. Then, for each event, we examined whether the likelihood of an event being remembered among all participants correlated with its mean neural pattern similarity with the previous event (i.e., *Cross-event correlation*). The recall rate for an event was the proportion of participants that remembered it. At the same time, pattern similarities of both activation and connectivity patterns were calculated and averaged across all participants, generating the neural transition indices across participants. Recall rates for all 49 events (the last event is not available) and their corresponding pattern transition measures were then correlated, providing a further indication of how subsequent memory related to pattern transitions across boundaries (**Figure S10** and **Figure S15**).

As exploratory analyses, we also applied *cross-participant analyses* to all ROIs. More specifically, for each ROI, we asked whether subjects with higher/lower neural similarity are better/worse at subsequent memory recall. However, individual differences analysis between neuroimaging measures and behaviors requires a relatively larger sample size to be reliable^4^. Considering the small sample size (N=17) of the discovery dataset, our results can be only seen as preliminary analyses. As shown in **Figure S11** and **Table S3**, individual difference analyses revealed a similar statistical pattern as results from the within-subject contrast analyses in the hippocampus, early auditory area, and posterior parahippocampal gyrus (*although some of the effects are not statistically significant due to power issue*): subjects with average lower activation pattern similarity during movie watching tend to perform better at the memory test, as suggested by higher recall rate (early auditory: r=-0.37, p=0.13; Post. Parahipp.: r=-0.40, p=0.11). By contrast, subjects with higher connectivity pattern similarity were more likely to perform better at the memory test (early auditory: r=0.45, p=0.06; hippocampus: r=0.40, p=0.10). Notably, the hippocampus and early auditory area were the two regions among six regions that demonstrated significant relationships between neural similarity and memory recall during within-subject paired t-test.

**6. Analyse subsequent memory effects using mixed-effect models**

We further used the mixed-effects model to examine the relationship between neural similarity and memory, considering both participants and events as random effects^5^. We closely followed the methodology used by a previous study^6^ which also used the mixed-effects model in the data analyses of fMRI data during movie watching. In brief, mixed-effects models were fitted using packages in R (i.e., *lme4, lmerTest, MuMIn*), with the following formulas:

① neural similarity^note1^ ~ memory^note2^ + (1|event) + (1|participant)

*Note1: neural similarity could be activation pattern similarity or connectivity pattern similarity.*

*Note2: memory is the subsequent recall of the first event in an event pair (i.e., R or F)*

Beyond the raw model (i.e., formula ①) above, we ran a second model ② incorporating multiple event-specific situational variables (e.g., *event duration, location, music*…).

② neural similarity ~ memory + duration + place + music + arousal + valence + (1|event) + (1|participant)

To control for the effects of low-level perceptual features on our subsequent memory analysis, we ran a third model including perceptual variables (i.e., *corrs, dist, and lum*).

③ neural similarity ~ memory + corrs + dist + lum + (1|event) + (1|participant)

We firstly focused on the results of the hippocampus and early auditory area because they were two regions that showed subsequent memory effects in both previous activation and connectivity pattern similarity. Mixed-effects analyses based on formula ① revealed that the relationship between memory and hippocampal activation pattern similarity (F=3.48, *p*=0.06, R^2^=0.004) or connectivity pattern similarity (F=3.68, *p*=0.05, R^2^=0.003) failed to reach significance, but suggested the same tendency as results from paired t-tests (i.e., *lower hippocampal activation pattern similarity, but higher connectivity pattern similarity is associated with better memory*). For the early auditory area, lower activation pattern similarity showed the tendency to correlate with memory (F=3.07, *p*=0.08, R^2^=0.003), while there is no memory effect for the connectivity pattern similarity (F=1.27, *p*=0.25, R^2^=0.001).

Next, we assessed how the above results are affected by situational covariates. When covariates were included in the model (i.e., *formula* ②), hippocampal activation pattern similarity (F=2.80, *p*=0.09), and connectivity pattern similarity (F=2.81, *p*=0.09) continued to show the same tendency of memory effects but failed to reach significance. For the early auditory area, lower activation pattern similarity showed the tendency to correlate with memory (F=3.00, *p*=0.08), while there is no memory for the connectivity pattern similarity (F=0.58, *p*=0.44). By the way, we found the significant fixed effects of two variables (i.e., *duration and arousal*) in some models (also see **Figure S5**).

In addition, we investigated how low-level perceptual features affected our subsequent memory analyses. In the third model (i.e., *formula* ③), hippocampal activation pattern similarity (F=4.34, *p*=0.03), and connectivity pattern similarity (F=3.81, *p*=0.05) continued to show significant memory effects. For the early auditory area, lower activation pattern similarity correlated with memory (F=4.06, *p*=0.04), while there is no memory for the connectivity pattern similarity (F=1.21, *p*=0.27).

Results from all ROIs can be found in **Table S4-S5** below. In sum, modeling participants as the random effect and adding covariates into the model did not change the overall pattern for the activation pattern analyses (*although the exact values of significance changed*). However, these steps had impacts on the connectivity pattern analyses: in the mixed-effect models, (1) the hippocampus is the only ROI that remained to show the association between connectivity pattern similarity and memory; (2) the connectivity pattern similarity of early visual, early auditory area, and PMC did not show memory effects anymore, even though subsequent memory effects of these ROIs can be detected by the paired t-tests.

**7. Alternative order memory analysis**

In the main text, our method of computing “*order memory*” confounds with “*event memory*”. We reanalyzed the “*order memory*” controlling for the effect of (first) event recall. More specifically, we first identified all remembered neighboring events regardless of their order and then compared event pairs which were recalled in the correct order (e.g. event i -> event i+1) and incorrect order (e.g. event -> and event i). This method can give us a more “pure” measure of order memory compared to our old method.

The new method revealed limited effect of “*order memory*” on connectivity pattern similarity among six ROIs (early visual cortex: t =-0.02 , p_raw_ =0.97 , Cohen’s d =-0.007; hippocampus: t =1.27 , p_raw_ =0.22 , Cohen’s d =0.31; early auditory area: t =-0.89 , p_raw_ =0.38 , Cohen’s d =-0.22; posterior parahippocampal gyrus: t =0.76, p_raw_ =0.45, Cohen’s d =0.19; mPFC: t =0.65 , p_raw_ =0.52 , Cohen’s d =0.16; PMC: t =1.06, p_raw_ =0.52, Cohen’s d =0.166). These non-significant results are hard to interpret, but one of the reasons could be limited power: no enough event (“trial”) pairs can be extracted for the certain contrast when analyses were restricted to the remembered event pairs (in-order pairs: mean=19.47, SD=5.03; out-order pairs: mean=5, SD=5.03).

**8. Control analyses for event distance analyses**

We ran several control analyses for event distance analyses (1) to evaluate the effects of potential artifacts (i.e., *temporal distance* and *temporal filtering*) on the event distance analysis; (2) the relationship between event distance and memory was tested with the permutation test, in which memory labels (i.e., R and F) were shuffled randomly; (3) event distance analysis was also applied to ROIs beyond the hippocampus.

(1) *Temporal distance*. As depicted in **Figure S18**, **panel A and B** showed how hippocampal pattern similarity changes with event distance when there is no TR adjustment applied. Controlling for different numbers of TRs between events, **panel C and D** illustrated how these adjusted hippocampal pattern similarities change with event distance. One big difference before and after the adjustment is that most of the variances between different event pairs (with the same event distance) were controlled. We fitted the linear regression model to these adjusted hippocampal pattern similarities and found the same relationship between pattern similarities and event distances which were reported in our initial submission. For the activation pattern similarity, the shorter the event distance, the more distinct the hippocampal activation patterns (r=0.24, *p*<0.001, **Panel E**). This positive correlation was largely driven by the negative correlations between events when the event distance is smaller than 5. For the connectivity pattern similarity, the shorter the event distance, the more similar the hippocampal connectivity patterns (r=-0.51, *p*<0.001, **Panel F**).

*Temporal filtering*. In the main text, high-pass filtering (140 s cutoff) was applied to fMRI data. Of note, the 140s cutoff was used in several previous fMRI studies with movie stimuli (e.g. Lerner et al., 2011) including the original study^7^ that used the same Sherlock dataset. We further used different cutoffs for high-pass filtering and re-ran the hippocampal event distance analysis. **Figure S19** illustrated how hippocampal neural similarity (after controlling for different numbers of TRs) changes with event distance under different high-pass filtering cutoffs: [1] different cutoffs affected the exact number of activation pattern similarity when the event distance is smaller than 4, but all of them were negative and the general pattern (i.e., the shorter the event distance, the more distinct the activation patterns) existed regardless of the specific cutoff used (**Figure S19A**); [2] different cutoffs did not affect the connectivity pattern similarity, as suggested by inseparable lines (**Figure S19B**).

(2) We investigated how memory modulates the relationship between event distance and pattern similarity in all ROIs and validated the memory relevance using the permutation (i.e., *shuffle*) method. As shown in the right panels (i.e., *data=shuffle*) of **Figure S20** and **Figure S21**, when we randomly shuffled memory labels (for 1000 times in our analysis), the relationship between event distance and neural similarity was not modulated by memory anymore and demonstrated almost identical effects across all ROIs.

(3) Critically, we found the region-specific modulation of memory on event distance analysis (*see* *left panels; data=real*). More specifically, data patterns that were shown in these figures were largely consistent with reported SME effects in our initial submission (i.e., *Figure3 and Figure4 of the main text*): activation pattern similarities of the early visual area and PMC did not associate with subsequent memory recall (*Figure 3A* and *Figure 3D*). In the current event distance analysis, memory has limited effects on distance X similarity interaction in the early visual area and PMC (**Figure S20**). Similarly, connectivity pattern similarities of the mPFC and pPHG did not correlate with subsequent memory (*Figure 4C and Figure 4F*). And when we plot lines for remembered and forgotten events separately, the two lines (i.e., *R and F*) of these two regions are also largely overlapping (**Figure S20**).

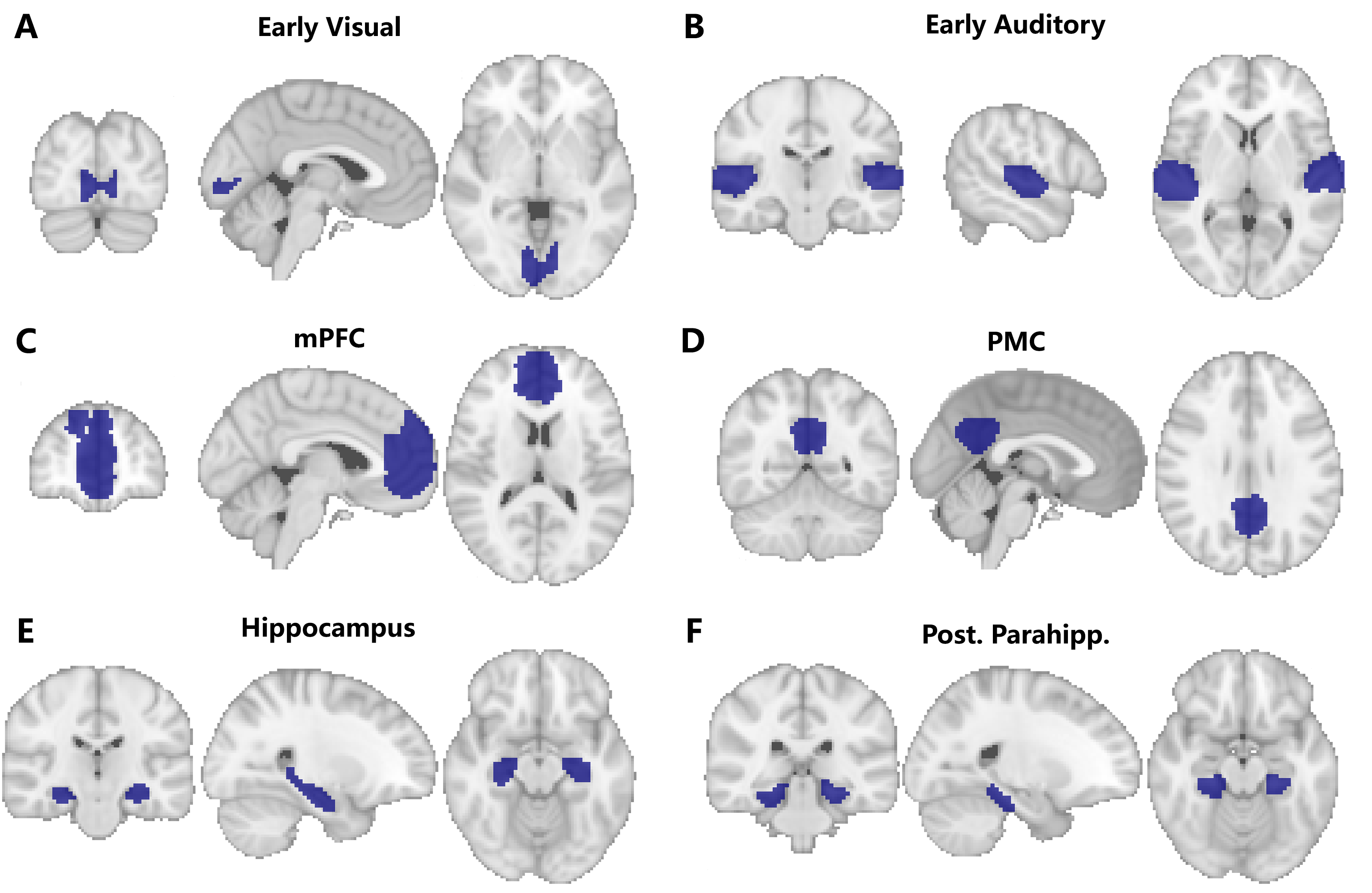

**Figure S1. Six predefined regions-of-interest (ROIs) used. (A) (B)** Early visual and auditory cortex were functionally defined in the literature using inter-subject correlation analysis. **(C) (D)** The medial prefrontal cortex (mPFC) and primary motor cortex (PMC) were defined in the functional atlas of the resting-state default mode network. **(E) (F)** The hippocampus and posterior parahippocampal gyrus were anatomically defined from the probabilistic *Harvard-Oxford Subcortical Structural Atlas* using the threshold of 50%.

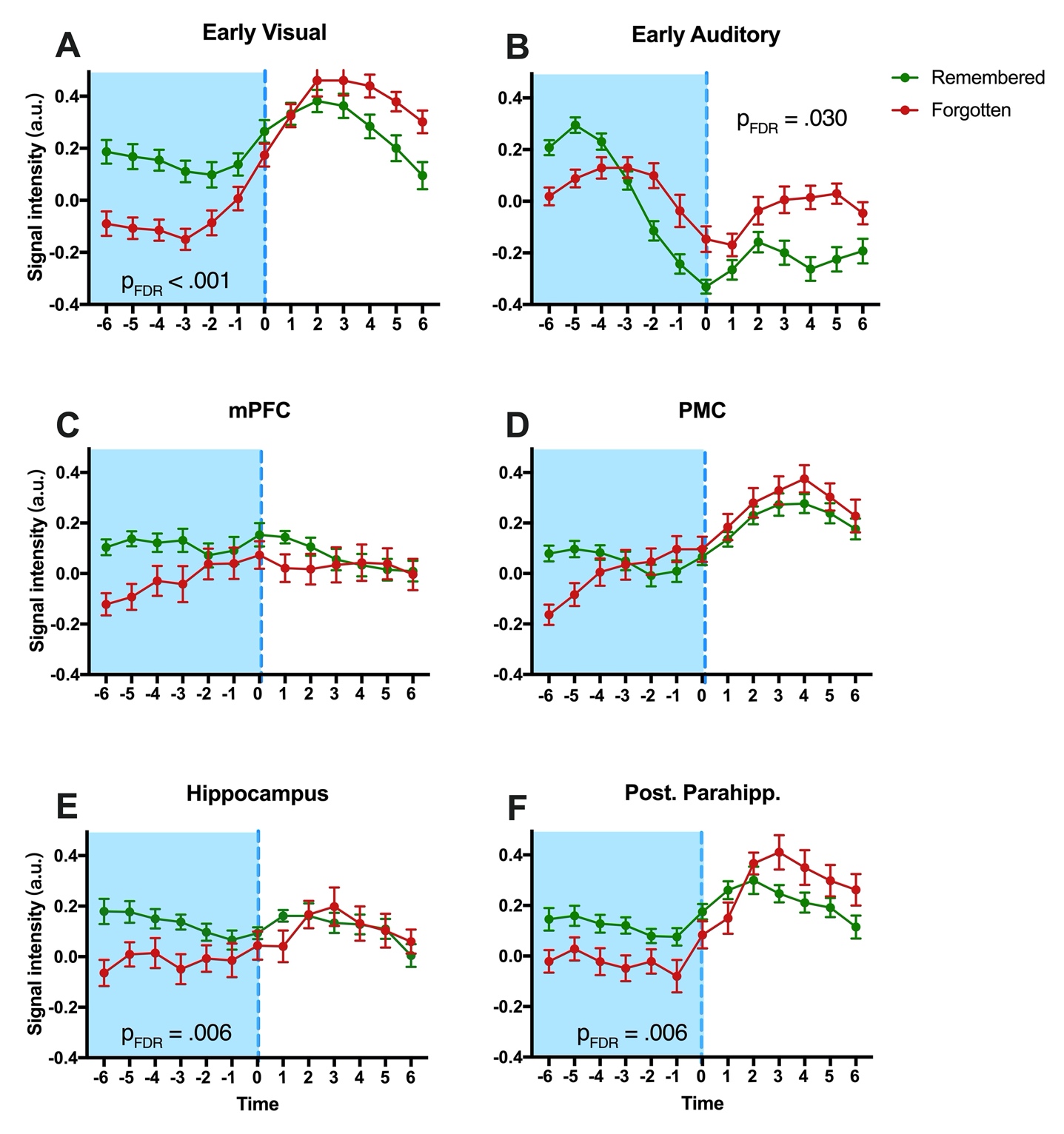

**Figure S2. Activity time courses of *remembered* and *forgotten* events around event boundaries in six ROIs.** We compared activity time courses of sequential event pairs (*Remembered* vs*. Forgotten*) based on subsequent memory performance of the first event across six ROIs using repeated two-way ANOVA (*memory*; *event*). The left side (blue) shows activity for an event that was later *remembered* or *forgotten*, while the right side (white) shows what happened to the signal after those events. The early visual cortex (*F* = 13.109, *p*_FDR_ = .006, η² = 0.45, panel **A**), hippocampus (*F* = 8.217, *p*_FDR_ = .017, η² = 0.339, panel **E**), and pPHG (*F* = 15.562, *p*_FDR_ = .006, η² = 0.493, panel **F**) demonstrated a significant interaction between *event* and *memory*. Post-hoc comparison revealed greater activation for *remembered* compared to *forgotten* events during the *current event*. Similar patterns were seen in PMC (*F* = 4.718, *p*_FDR_ = .054, η² = 0.228, panel **D**) and mPFC (*F* = 3.036, *p*_FDR_ = .101, η² = 0.160, panel **C**), but did not survive correction for multiple comparisons. The reverse effect (higher activation for *forgotten* vs. *remembered* events) was found in early auditory cortex (*F* = 10.852, *p*_FDR_ = .010, η² = 0.404, panel **B**). *Note: figures shows the mean and the standard error.*

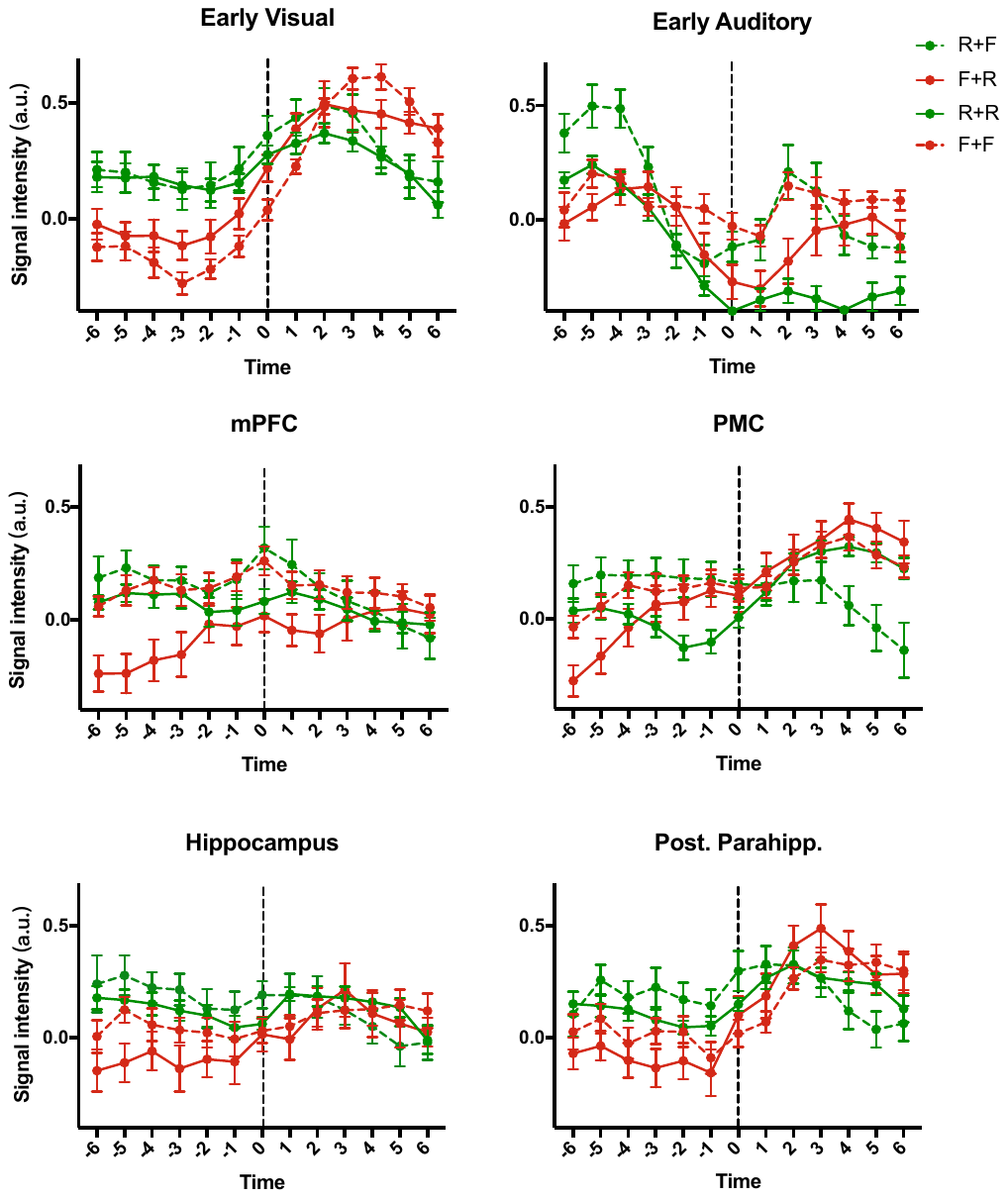

**Figure S3. Univariate response curves based on memory performance of two consecutive events.** BOLD signals around the event boundaries (6 TRs before and after event offset; 13 time points including the boundary itself) were extracted from the time series, averaged across all voxels, and z-scored within each region. Considering the memory label of the two consecutive events simultaneously, the time course segments were categorised into four types: R (green)/F (red) + R (solid)/F (dashed), and were plotted against the time (in TRs) relative to the event boundary. *Note: figures shows the mean and the standard error.*

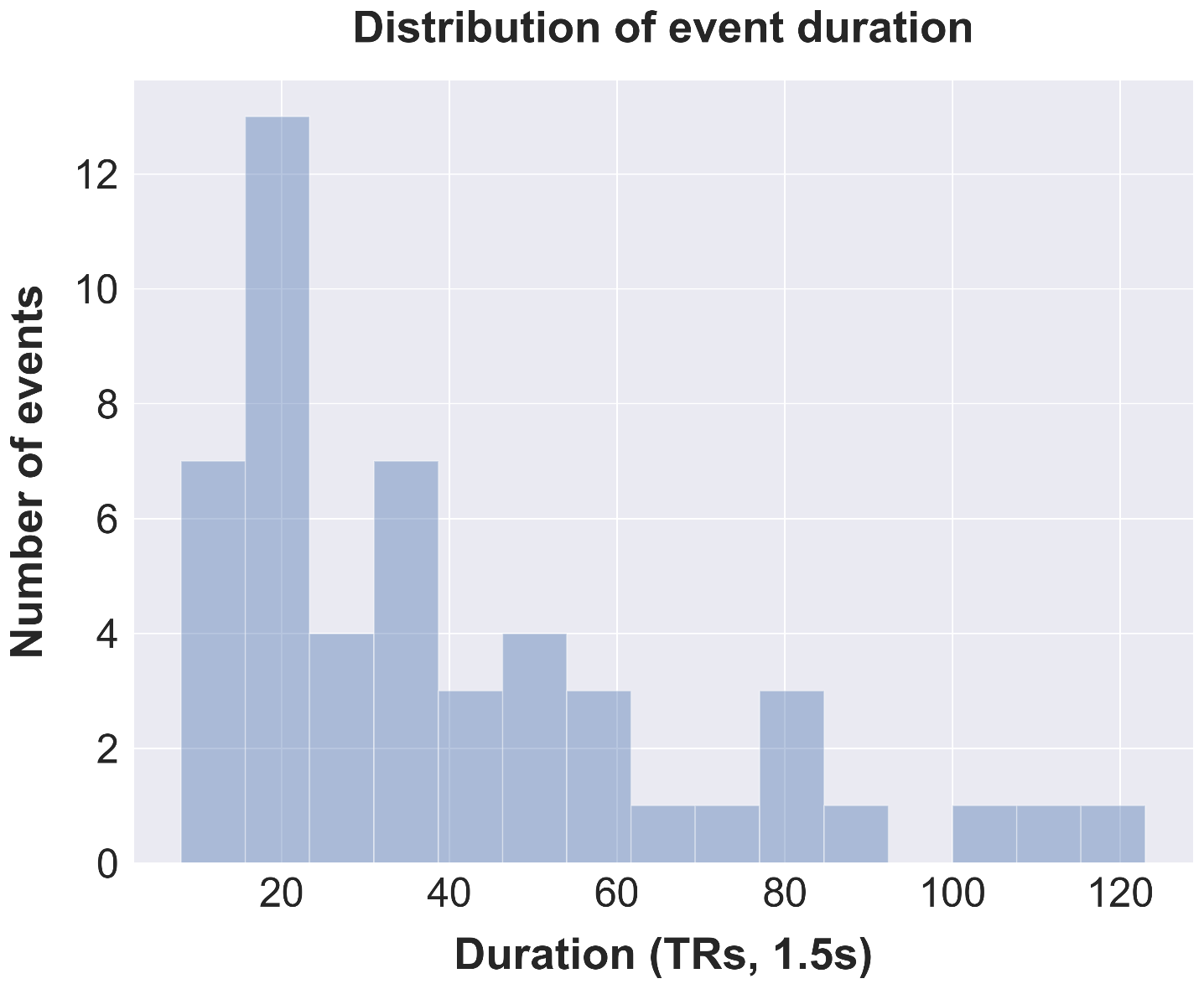

**Figure S4 Distribution of event durations (TRs) in the discovery dataset (total number of events=50).**

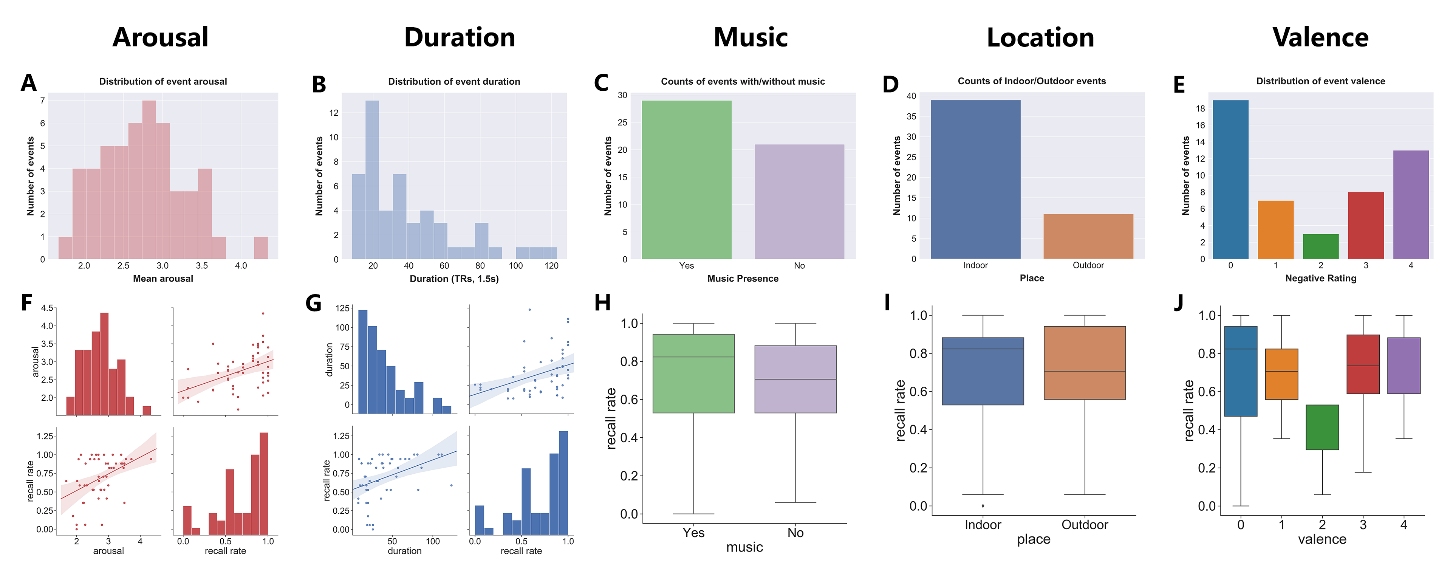

**Figure S5** **Event-specific situational variables and their relationship with memory**. (A-E) distributions of five variables (i.e., *arousal, duration, music, location, and valence*). (F-J) Association between five variables and memory recall rate. *Arousal* and *duration* positively correlated with the overall recall rate (%) while *music*, *location*, and *valence* did not associate with subsequent recall. *Note: valence was defined as how many of the total four raters labeled the event as the negative event. The higher the valence value is, the more negative the event*.

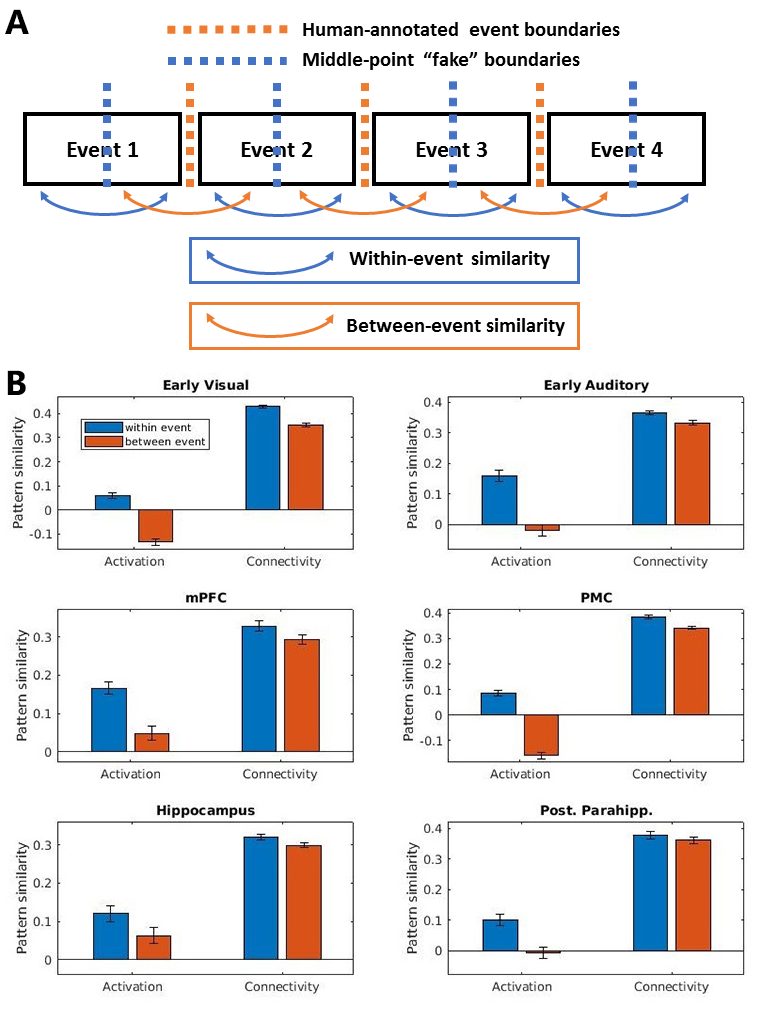

**Figure S6. Comparisons between within-event and between-event neural pattern shifts. (A)** The methodological framework of the comparisons. Both human-annotated event boundaries and middle-point “fake” boundaries were used to segment the fMRI time series. **(B)** Results in 6 ROIs. For both activation and connectivity pattern similarities, neural pattern shifts are larger for between-event transitions compared to within-event transitions. *Note: Error bar represents the standard error (SE).*

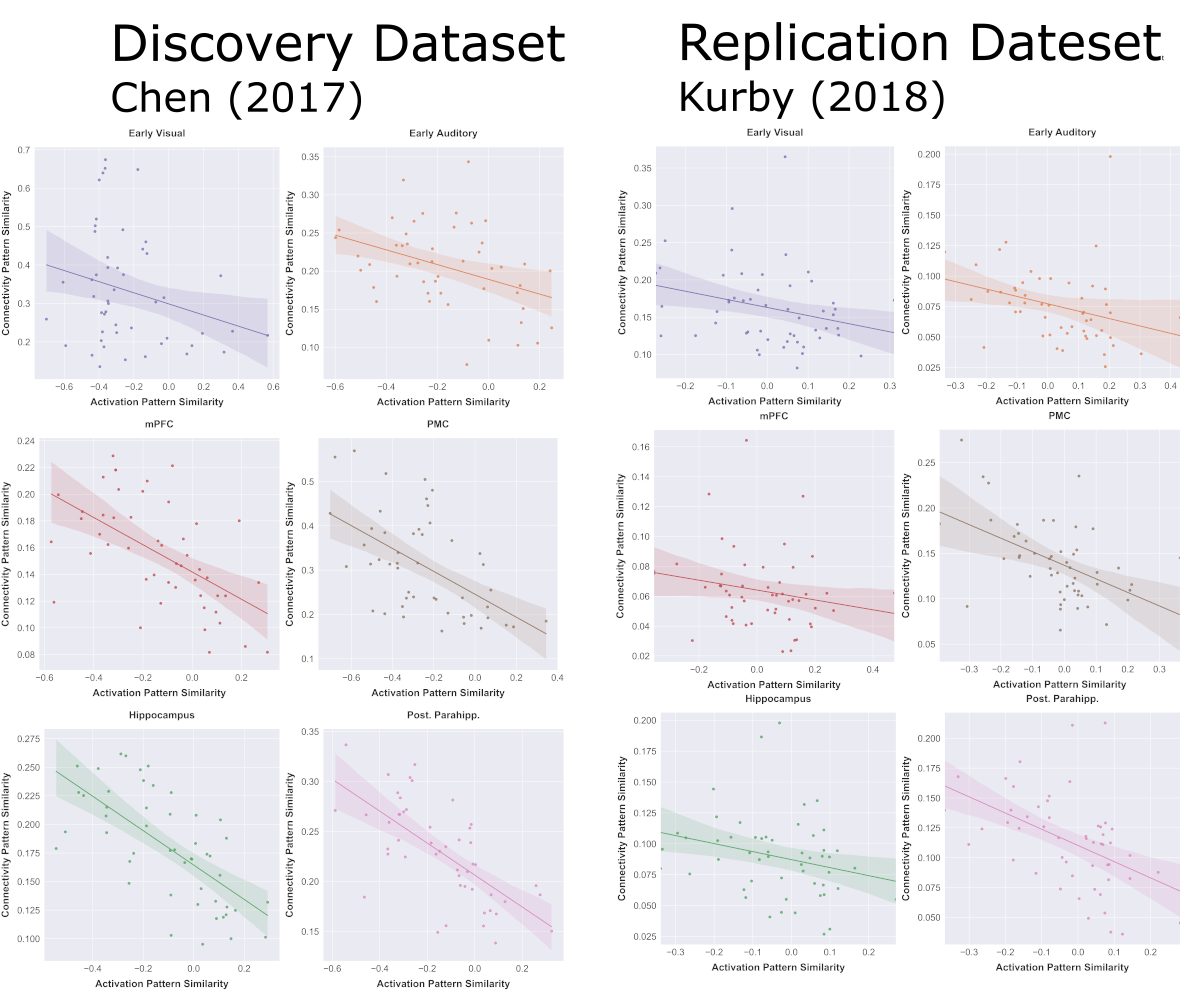

**Figure S7 Correlations between activation pattern similarities and connectivity pattern similarities across event boundaries in both the discovery and replication datasets.** Results revealed reliable region-specific negative correlations between two neural measures of pattern similarities for the early auditory cortex, PMC, and posterior parahippocampal gyrus (pPHG). Detailed statistics of correlation analyses can be found in **Table S3**.

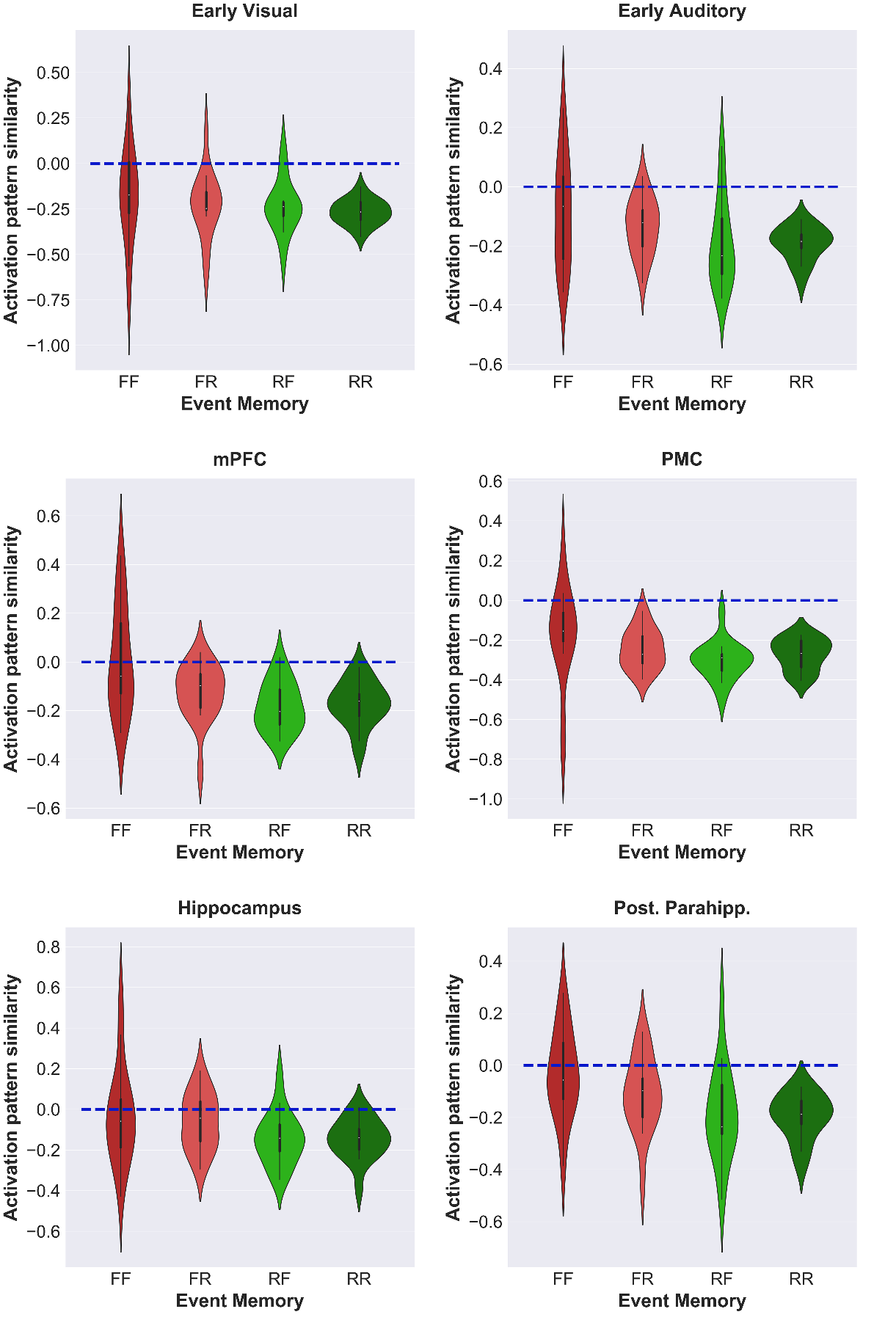

**Figure S8**. **Subsequent memory analyses of activation pattern similarity based on memory labels of event pairs**. *Both Forgotten (FF), first Forgotten and second Remembered (FR), first remembered and second forgotten (RF), and both remembered (RR))*

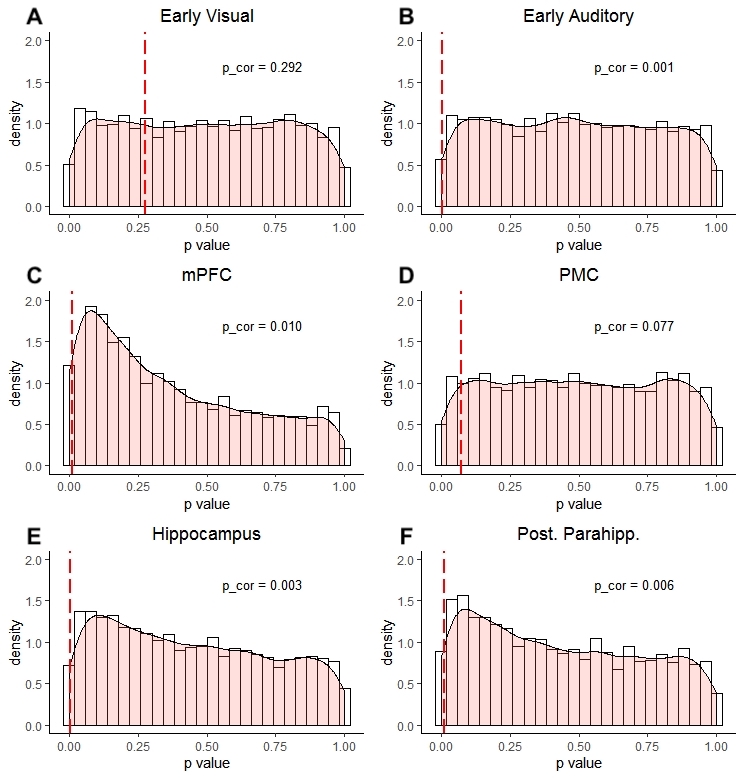

**Figure S9. Event boundary permutation analysis for *activation* *pattern* similarity.** The *p* value obtained from the true event boundary similarity analysis (red dashed line) is corrected based on the null distribution (histogram) generated by random permutation of event intervals (5000 times). A smoothed density estimate calculated by *stat_density* with *ggplot2* and *R* is indicated by the semitransparent region. The corrected values for each ROI are shown in the corresponding subplots.

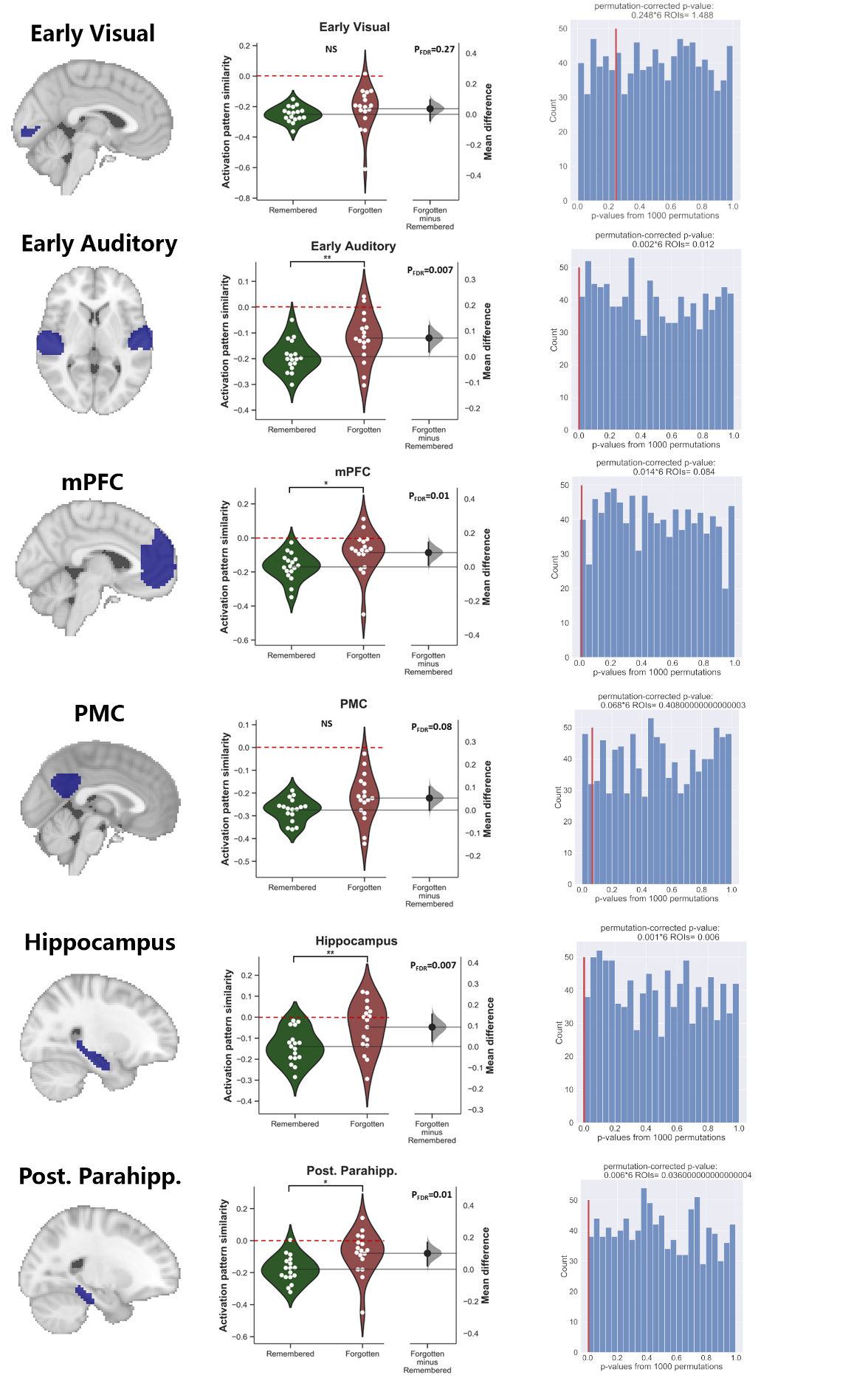

**Figure S10** **Comparisons of activation pattern similarities between Remembered and Forgotten events.** Comparisons based on real memory labels (*middle*). Estimation of permutation-based p-value based on shuffled memory labels (*right*).

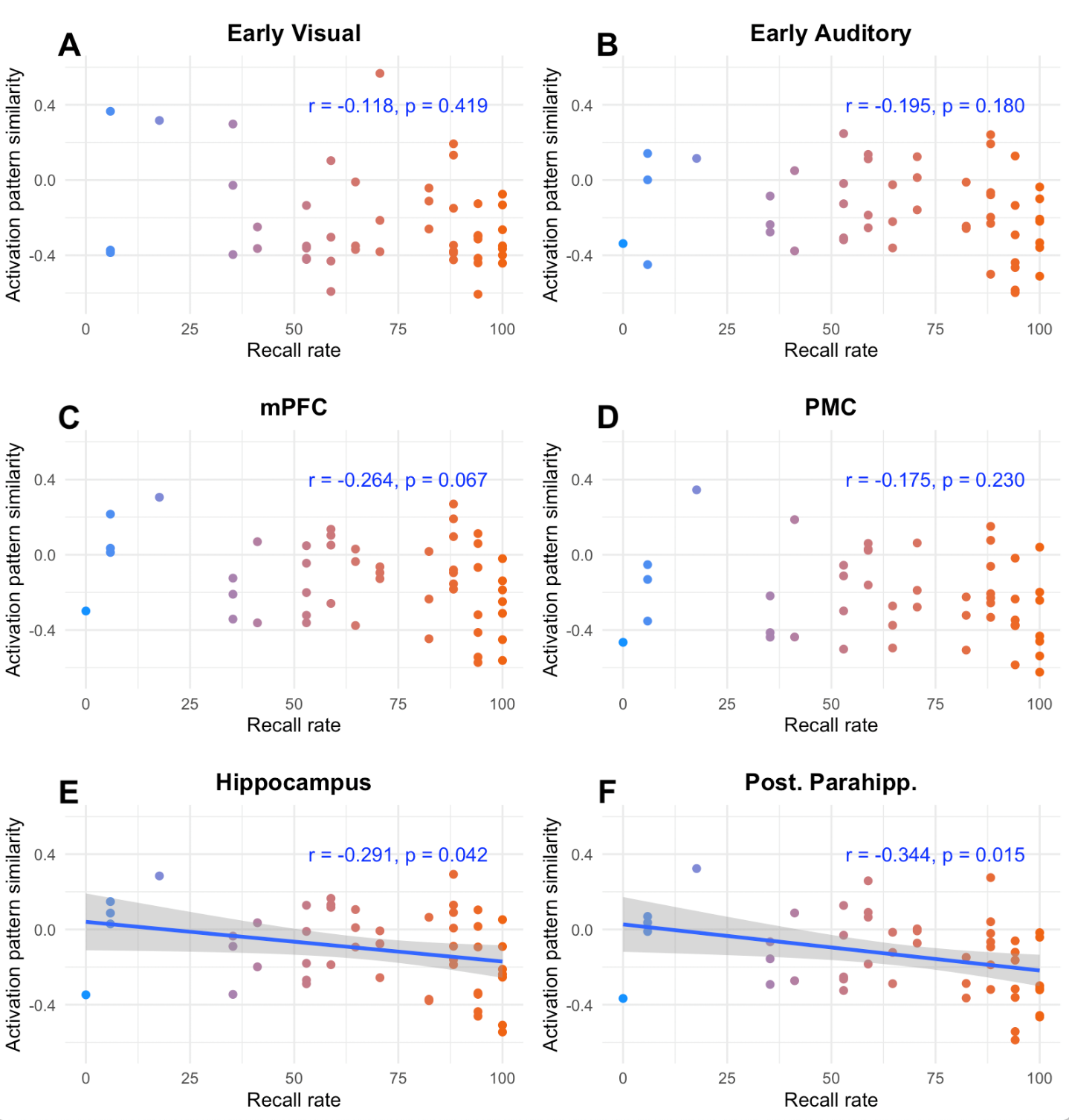

**Figure S11. Event-specific correlational analysis between *activation* *pattern* similarity and memory performance.** Memory performance per event is calculated by averaging the number of participants that successfully recalled the event divided by the total number of participants (recall rate). Pearson’s correlation coefficient *r* and *p*-value (uncorrected) are shown for each ROI. For the regions that demonstrated a significant correlation (i.e., hippocampus and pPHG), a linear model-based regression line (blue) is fitted to the data points with the confidence interval shown in gray. To better visualize the correlation, the color of the data points are set to change in the direction of the first principal component of the data.

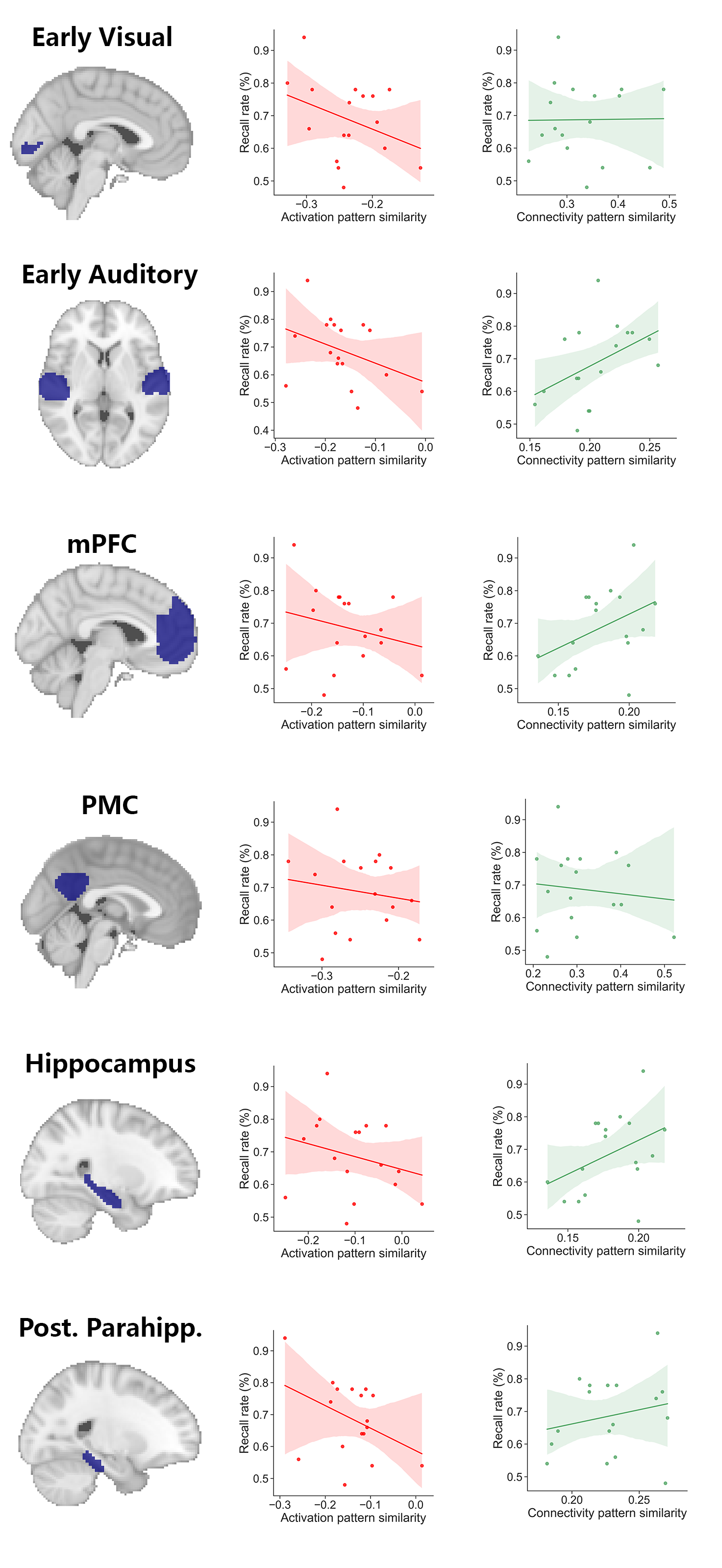

**Figure S12 Cross-participant correlation between neural pattern similarity and memory performance.** *Correlation with activation pattern similarity (Left; Red). Correlation with connectivity pattern similarity (Right; Green)*

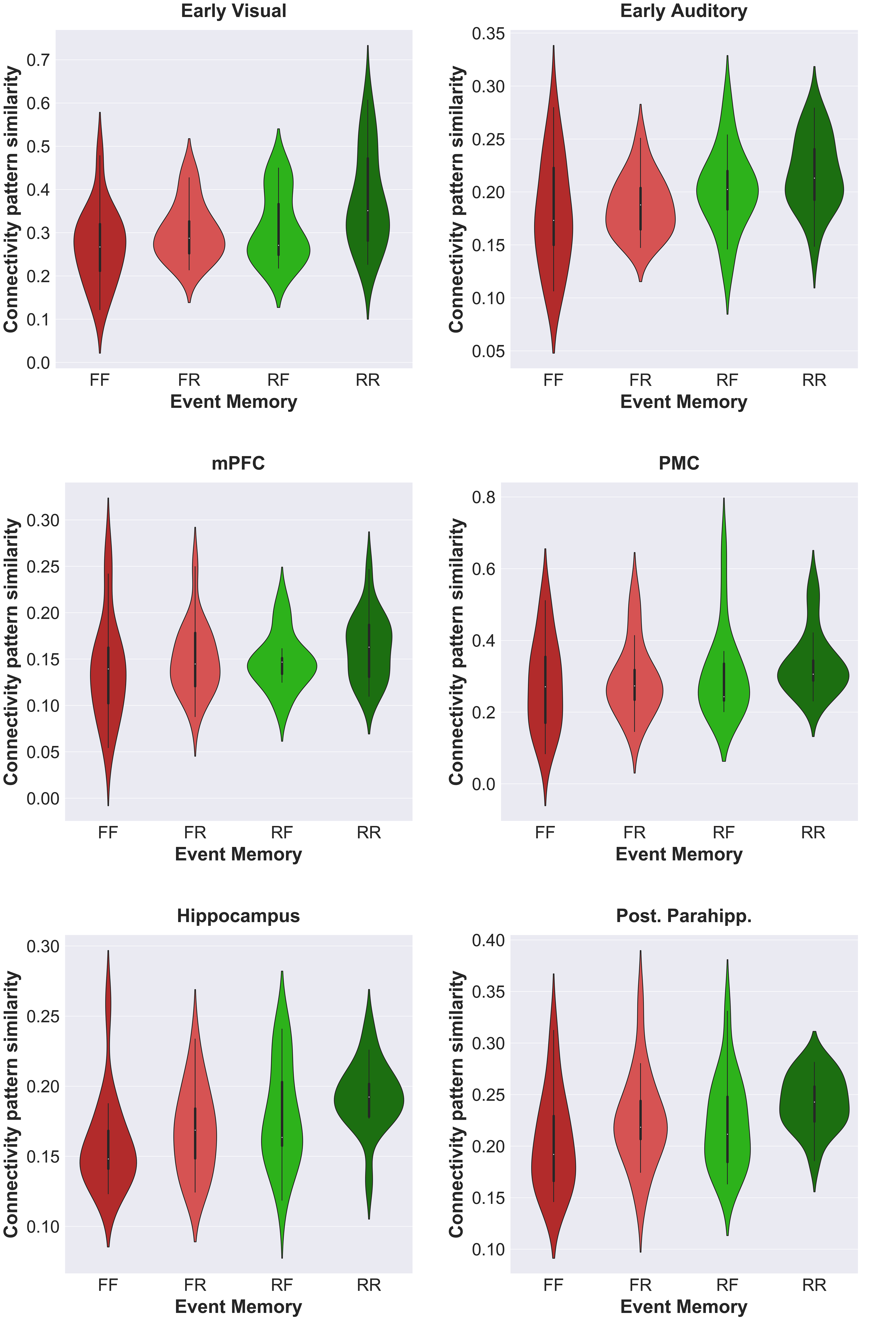

**Figure S13**. **Subsequent memory analyses of connectivity pattern similarity based on memory labels of event pairs**. *Both Forgotten (FF), first Forgotten and second Remembered (FR), first remembered and second forgotten (RF), and both remembered (RR))*

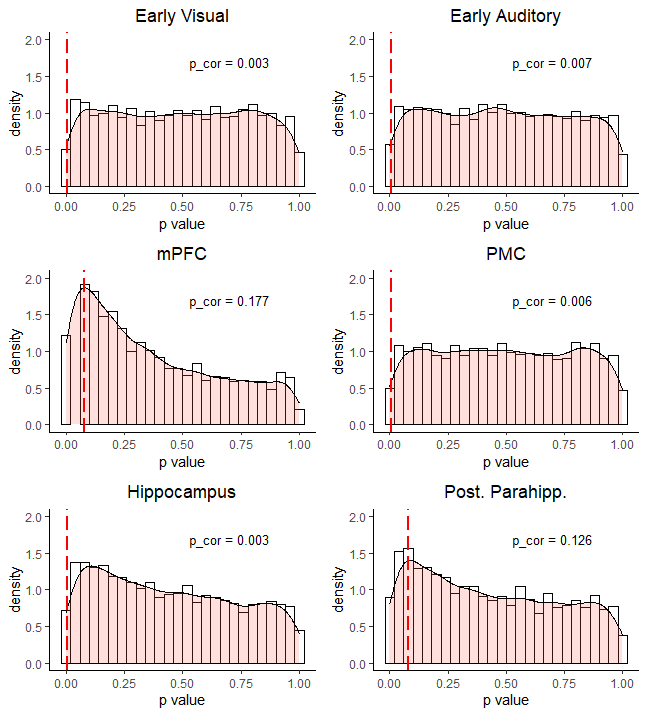

**Figure S14. Event boundary permutation analysis for *connectivity pattern* similarity.** Analysing *connectivity pattern* similarity with “scrambled” event boundaries. The *p* value obtained from the true event boundary similarity analysis (red dashed line) is corrected based on the null distribution (histogram) generated by random permutation of event intervals (5000 times). A smoothed density estimate calculated by *stat_density* with *ggplot2* and *R* is indicated by the semitransparent region. The corrected values for each ROI are shown in the corresponding subplots.

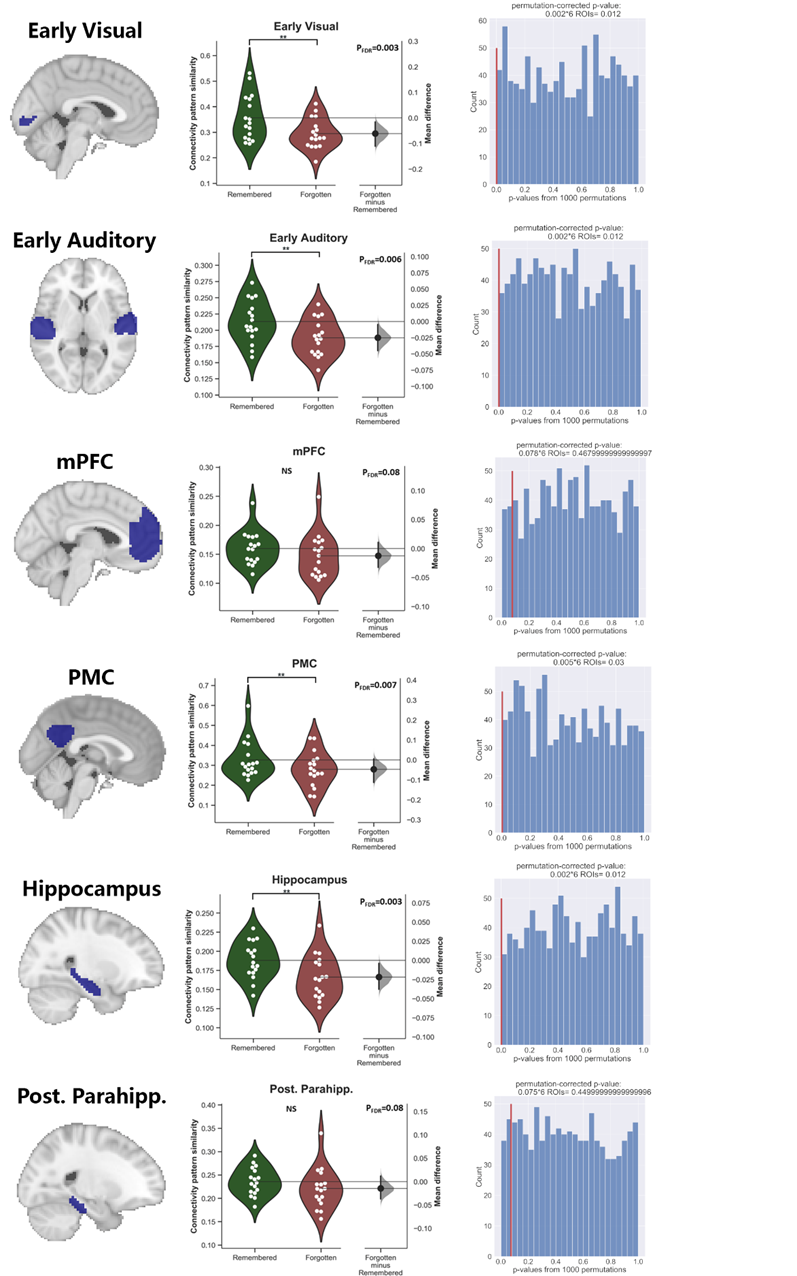

**Figure S15 Comparisons of connectivity pattern similarities between Remembered and Forgotten events.** Comparisons based on real memory labels (middle). Estimation of permutation-based p-value based on shuffled memory labels (right)

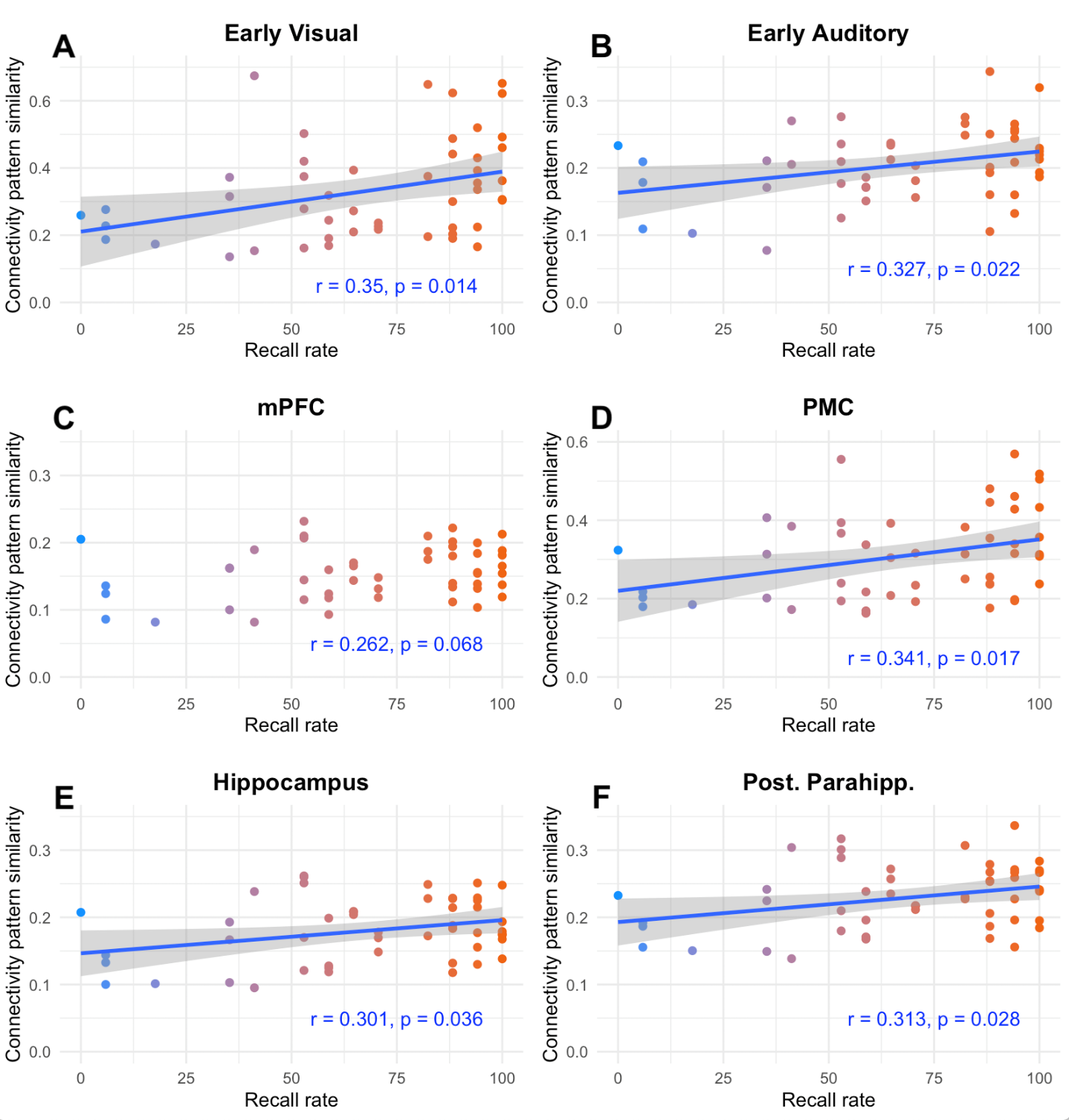

**Figure S16. Event-specific correlational analysis between *connectivity* *pattern* similarity and memory performance.** Memory performance per event is calculated by averaging the number of participants that successfully recalled the event divided by the total number of participants (recall rate). Pearson’s correlation coefficient *r* and *p* value (uncorrected) are shown for each ROI. For the regions that demonstrated a significant correlation (i.e., all six regions except for mPFC), a linear model based regression line (blue) is fitted to the data points with the confidence interval shown in gray. To better visualise the correlation, the color of the data points are set to change in the direction of the first principal component of the data.

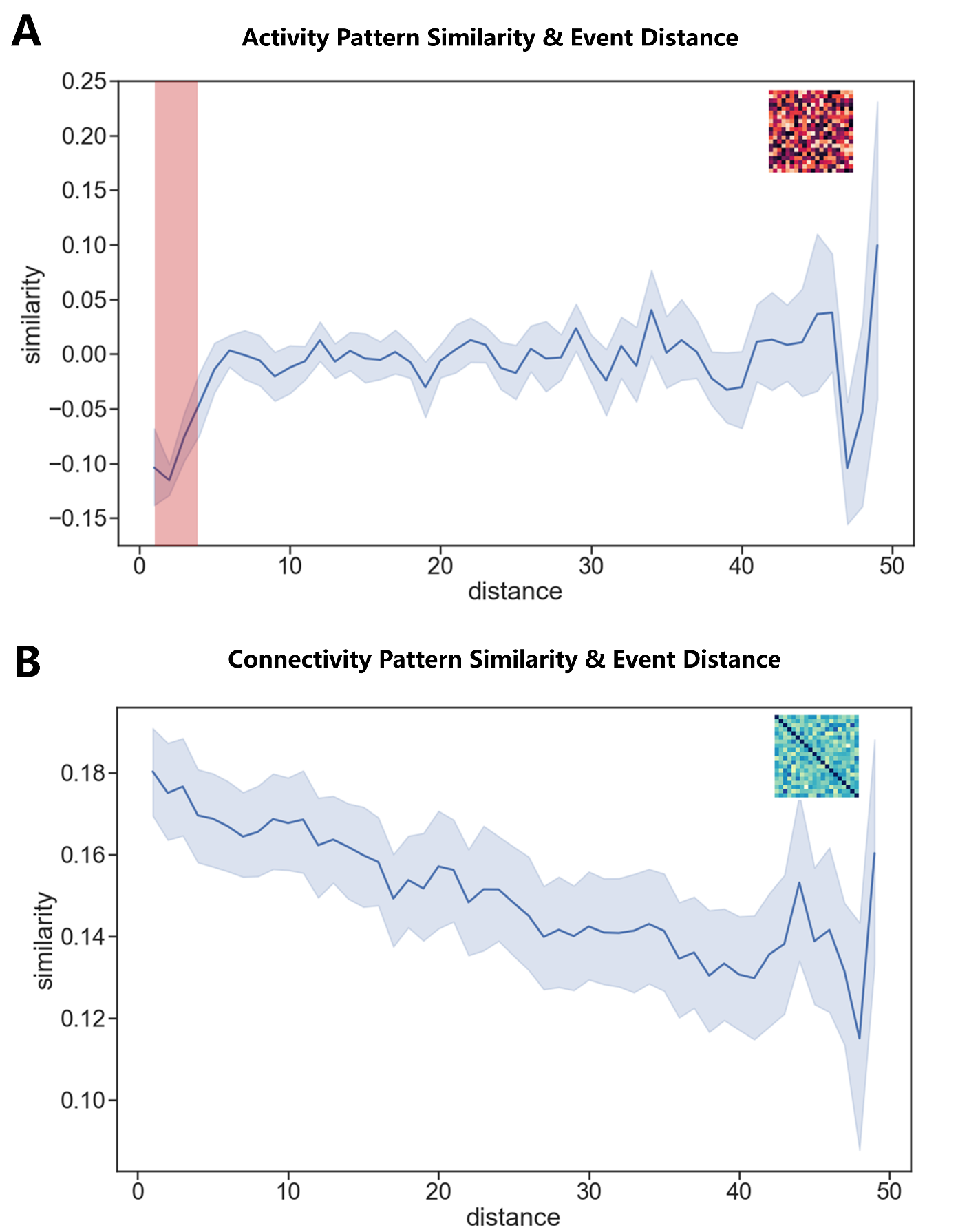

**Figure S17. Relationship between all possible event distance and hippocampal neural similarities.** (**A**) Relationship between hippocampal *activation pattern* similarities and event distance. Event distance ranges from 1 to 49. The shallow red bar indicates that the similarity is significantly lower than 0 after false discovery rate (FDR) correction. (**B**) Relationship between event distance and hippocampal *connectivity pattern* similarities. Event distance ranges from 1 to 49.

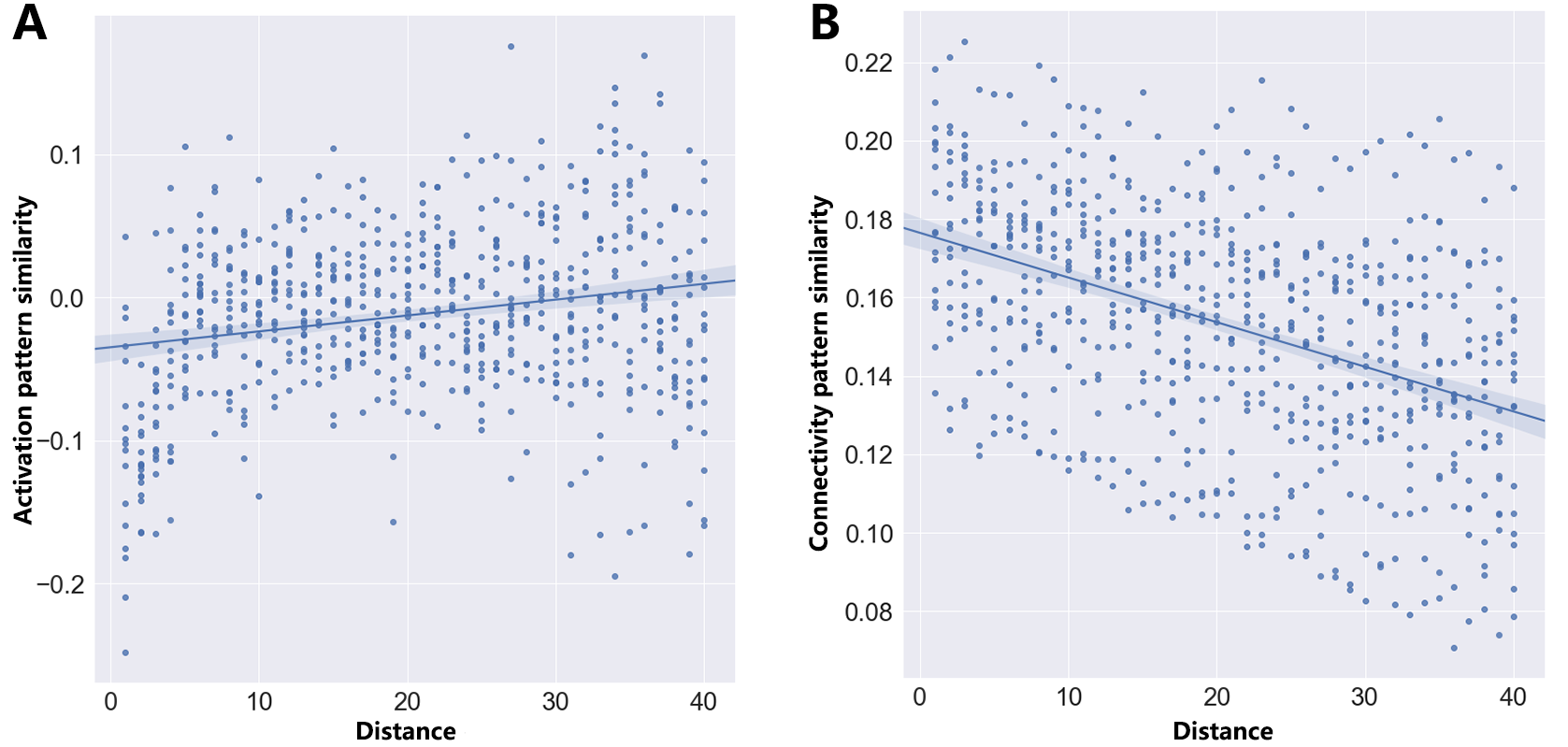

**Figure S18 Relationship between event distance and hippocampal *activation* and *connectivity pattern* similarities.** **(A)** Positive correlation between event distance (ranging from 1 to 40) and hippocampal *activation pattern* similarities (*r* = 0.21, *p*_raw_ = 1.8 × 10^-8^). **(B)** Negative correlation between event distance (ranging from 1 to 40) and hippocampal *connectivity pattern* similarities (*r* = -0.439, *p*_raw_ = 1.8 × 10^-33^).

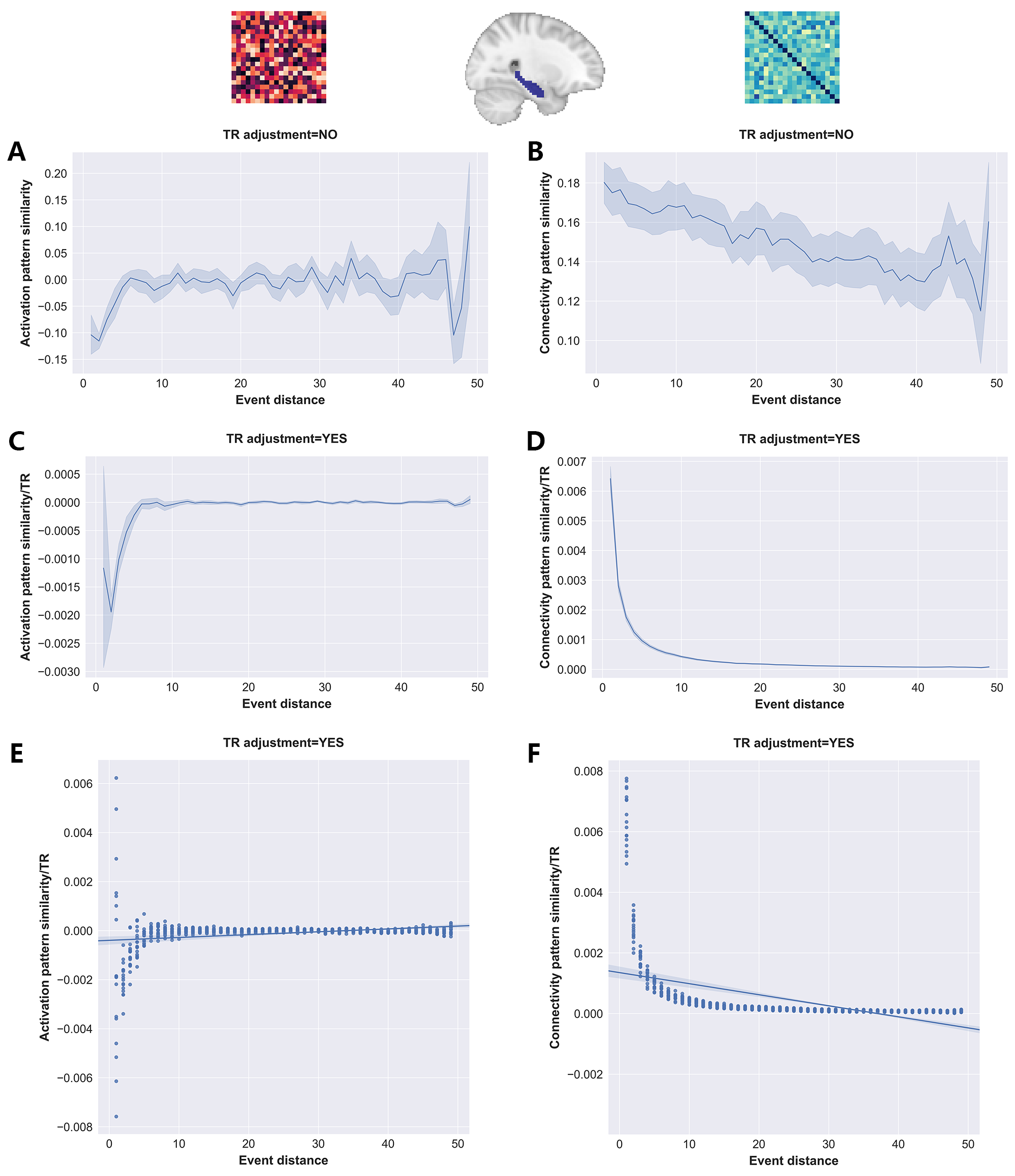

**Figure S19 The effect of controlling for temporal distance (i.e., *number of TRs*) on the hippocampal event distance analysis.** (A)(C)(E) presented results of hippocampal activation pattern similarity while (B)(D)(F) showed results of hippocampal connectivity pattern similarity. TR adjustment=whether the temporal distance between events were controlled.

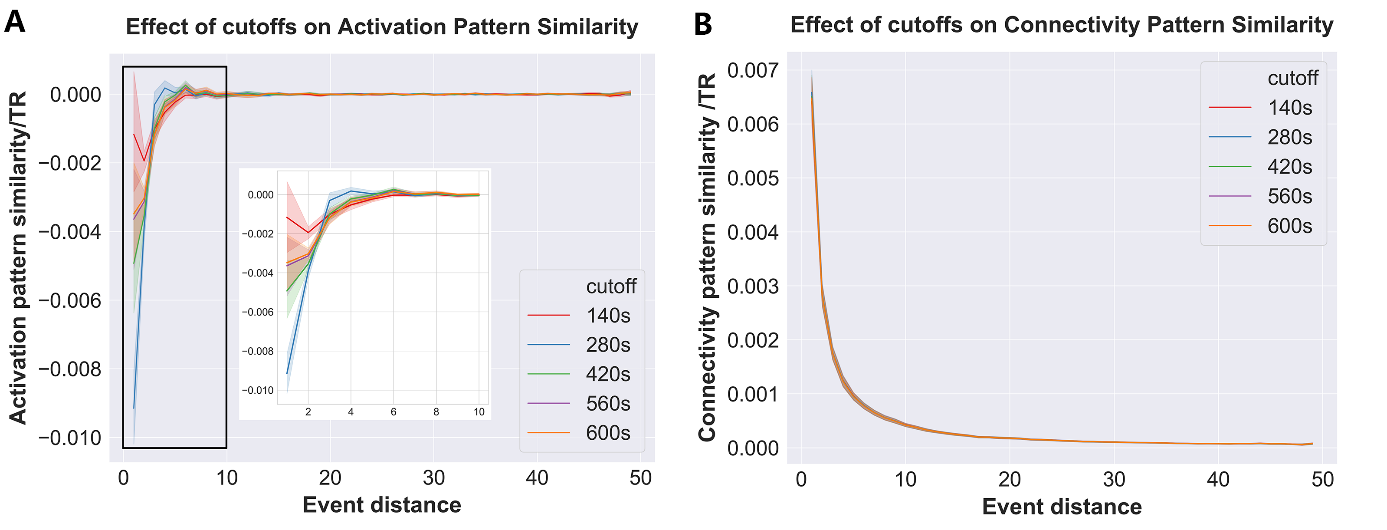

**Figure S20 The effect of different high-pass filtering cutoffs on the hippocampal event distance analysis.** Five different cutoffs (i.e., 140s, 280s, 420s, 560s, 600s) were used, and the relationship between hippocampal pattern similarity and event distance was demonstrated across five cutoffs. 140s is the cutoff used by the original study^7^ and us.

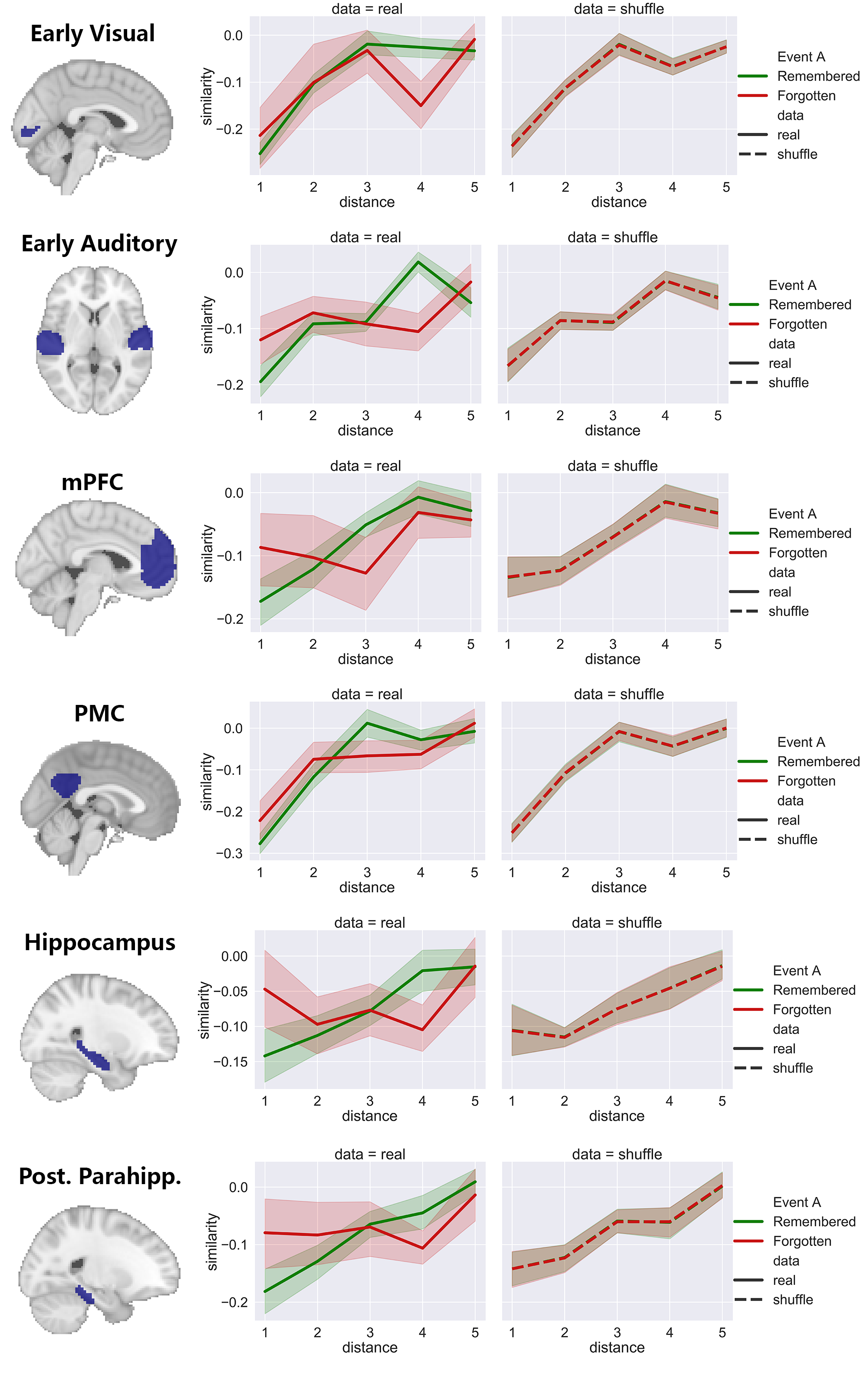

**Figure S21 Relationship between event distance, activation pattern similarity, and memory in all ROIs.** Left panels showed the relationship between similarity and distance using the real data. Right panels demonstrated the same relationship but based on the data which memory labels were shuffled with each participant 1000 times.

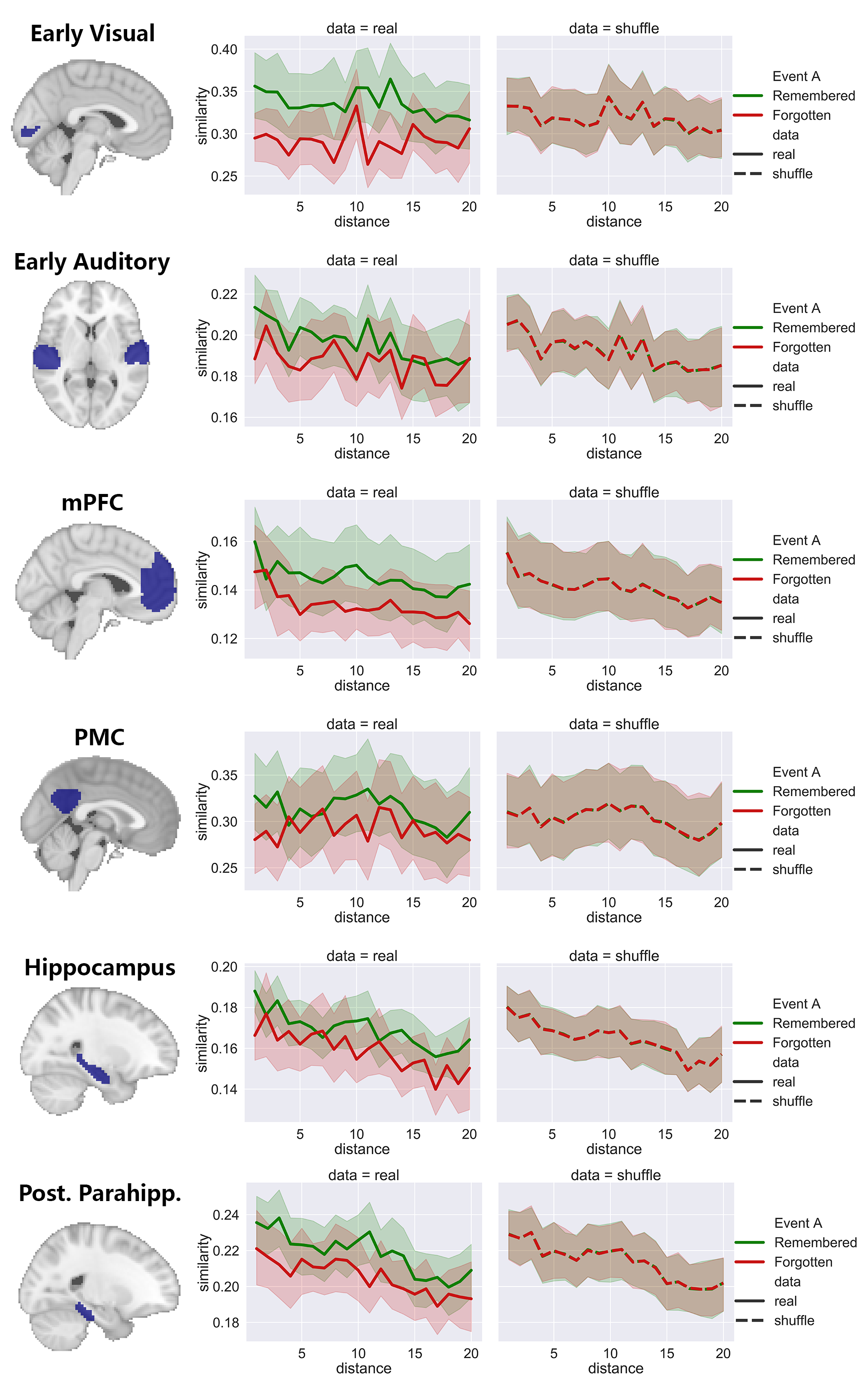

**Figure S22 Relationship between event distance, connectivity pattern similarity, and memory in all ROIs.** Left panels showed the relationship between similarity and distance using the real data. Right panels demonstrated the same relationship but based on the data which memory labels were shuffled with each participant 1000 times.

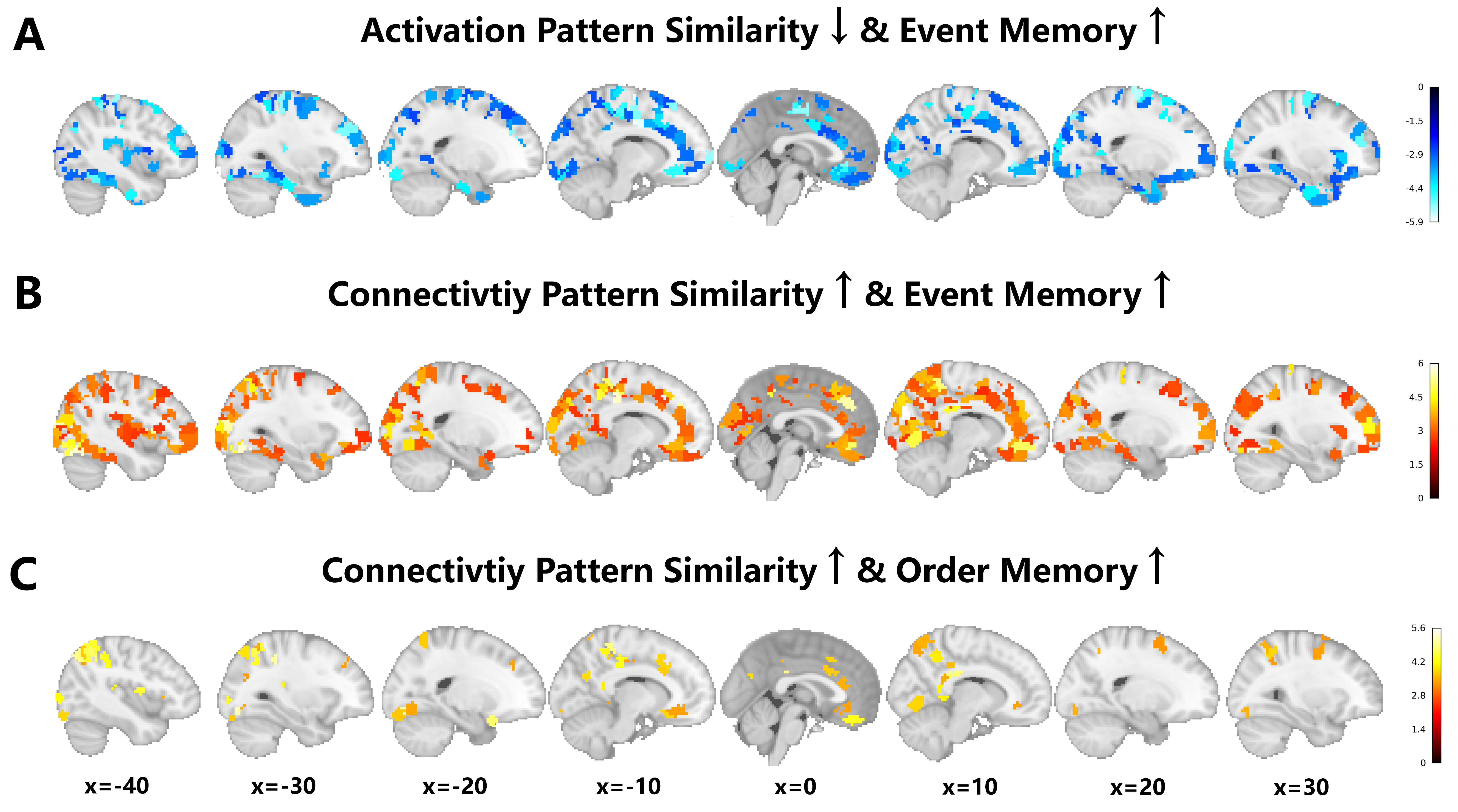

**Figure S23 Identifying event segmentation and integration computations separately across the neocortical regions.** (**A**) Distinct *activation patterns* of these brain regions across event boundaries correlate with better event memory. (**B**) Similar *connectivity patterns* of these brain regions across event boundaries relate to better event memory. (**C**) Similar *connectivity patterns* of these brain regions across event boundaries link to better order memory. All results are displayed at *p*_FDR_ < 0.05 across 1000 brain regions.

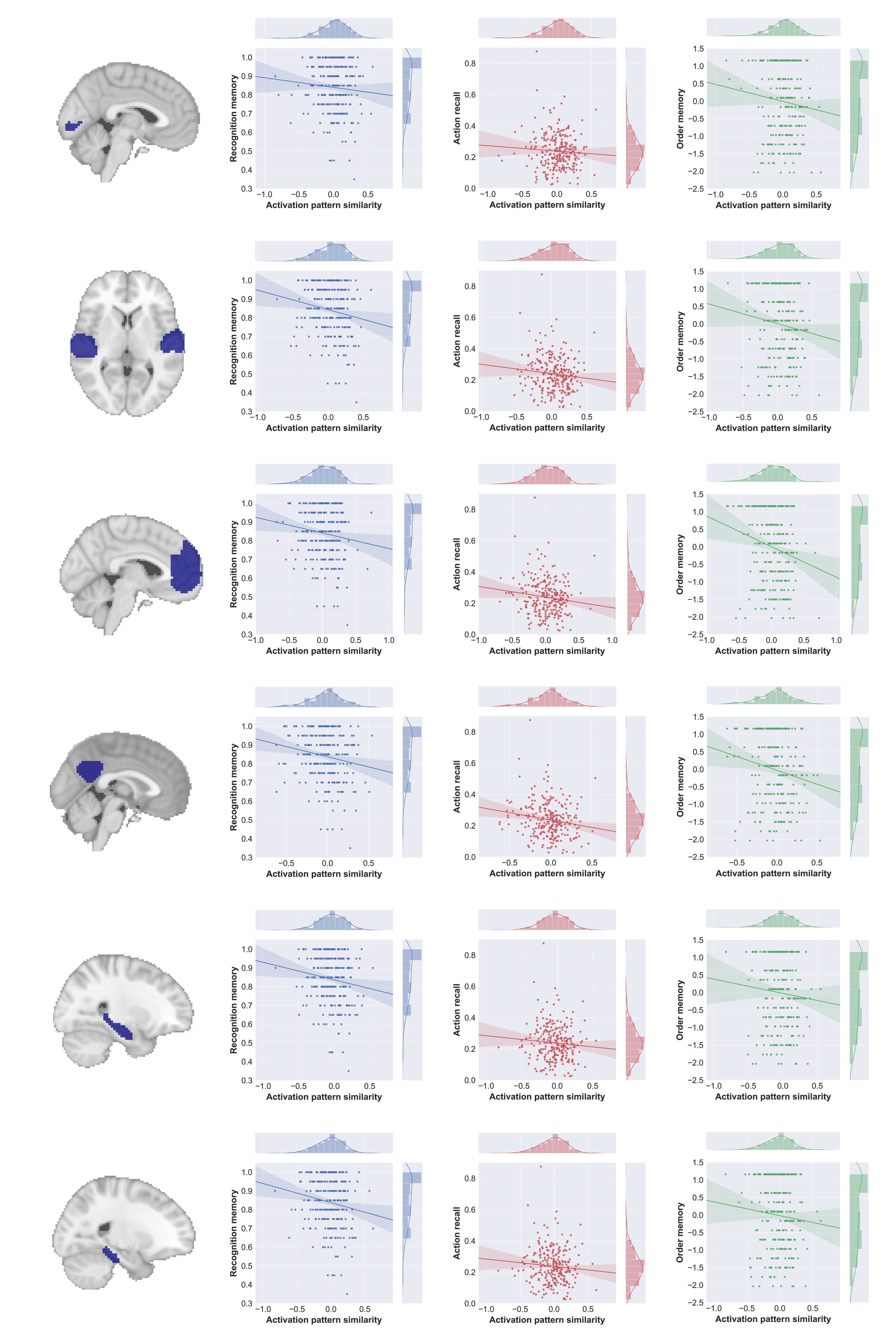

**Figure S24 Association between activation pattern similarity and three memory measures in the replication dataset.** *Correlation coefficients and corresponding p-values can be found in Table S5.*

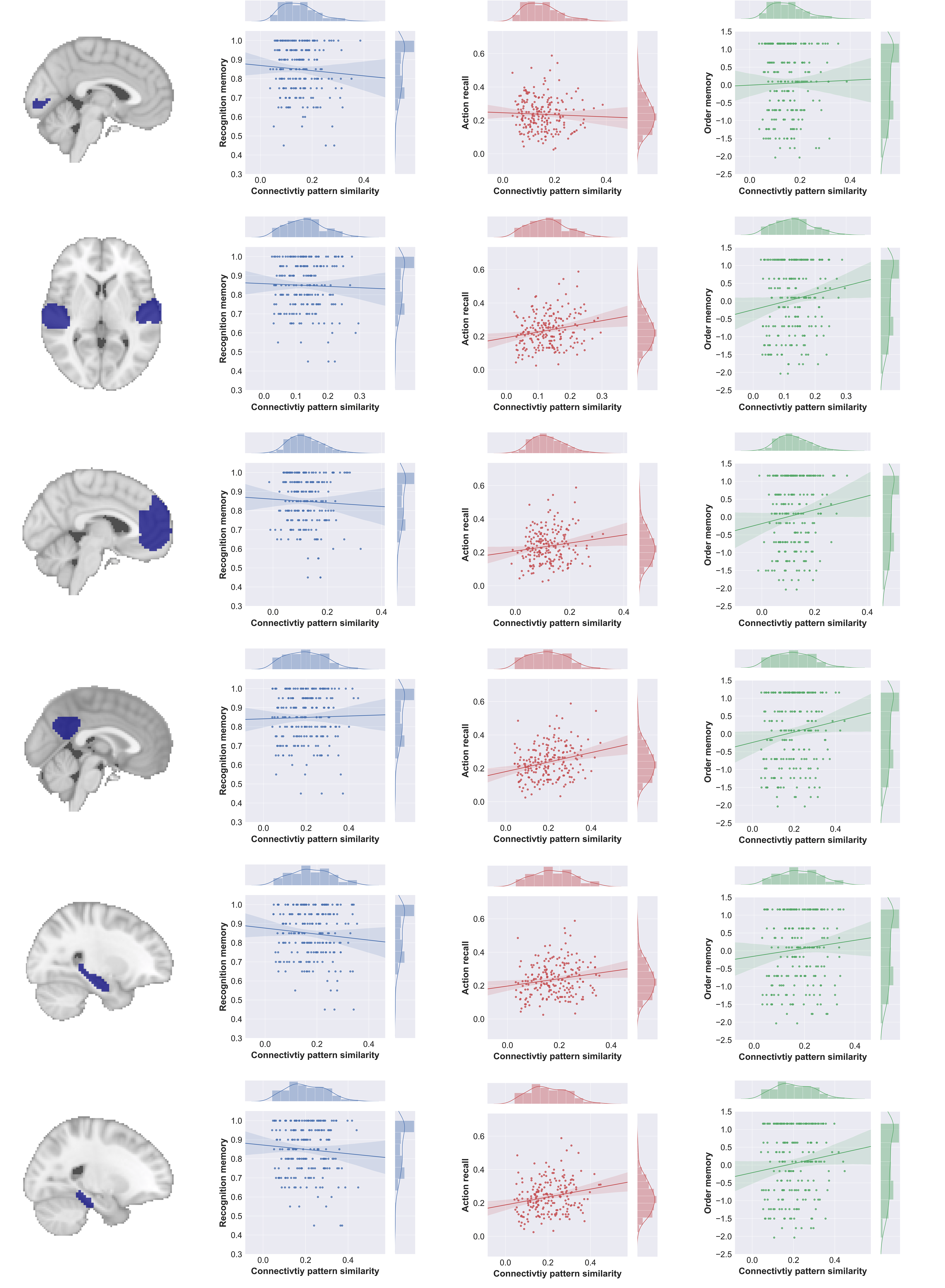

**Figure S25 Association between connectivity pattern similarity and three memory measures in the replication dataset.** *Correlation coefficients and corresponding p-values can be found in Table S5.*

| **Event_ID** | **Duration** | **Place** | **Music** | **Arousal** | **Valence** | **Recall rate (%)** |
| --- | --- | --- | --- | --- | --- | --- |
| **1** | 26 | Indoor | Yes | 2 | 0 | 0.058824 |
| **2** | 9 | Outdoor | No | 4.340909 | 4 | 0.941176 |
| **3** | 20 | Indoor | No | 3.45 | 4 | 0.882353 |
| **4** | 31 | Indoor | No | 1.673077 | 3 | 0.647059 |
| **5** | 23 | Indoor | No | 2.28125 | 3 | 0.882353 |
| **6** | 22 | Outdoor | Yes | 2.795455 | 2 | 0.058824 |
| **7** | 13 | Indoor | No | 2.09375 | 1 | 0.588235 |
| **8** | 12 | Indoor | Yes | 3.4 | 4 | 0.882353 |
| **9** | 17 | Indoor | Yes | 2.9375 | 4 | 0.705882 |
| **10** | 20 | Outdoor | Yes | 2.75 | 4 | 0.529412 |
| **11** | 12 | Indoor | Yes | 3.5 | 4 | 0.882353 |
| **12** | 22 | Indoor | Yes | 2.704545 | 3 | 0.647059 |
| **13** | 86 | Indoor | No | 3.025 | 3 | 0.823529 |
| **14** | 49 | Outdoor | No | 2.0625 | 1 | 0.941176 |
| **15** | 37 | Indoor | Yes | 2.988095 | 0 | 0.882353 |
| **16** | 107 | Indoor | Yes | 2.613636 | 0 | 1 |
| **17** | 20 | Indoor | No | 1.888889 | 3 | 0.176471 |
| **18** | 8 | Indoor | Yes | 3.5 | 4 | 0.352941 |
| **19** | 81 | Indoor | No | 2.135714 | 0 | 1 |
| **20** | 19 | Indoor | No | 2.270833 | 0 | 0.058824 |
| **21** | 8 | Indoor | Yes | 2.6875 | 3 | 0.411765 |
| **22** | 38 | Indoor | Yes | 2.86 | 3 | 1 |
| **23** | 18 | Indoor | No | 2.204545 | 1 | 0.352941 |
| **24** | 50 | Indoor | Yes | 2.693182 | 0 | 0.941176 |
| **25** | 123 | Outdoor | Yes | 2.996094 | 0 | 0.588235 |
| **26** | 20 | Outdoor | No | 2.222222 | 0 | 0.352941 |
| **27** | 55 | Outdoor | No | 2.690789 | 1 | 0.705882 |
| **28** | 26 | Indoor | Yes | 2 | 0 | 0 |
| **29** | 43 | Indoor | Yes | 2.533333 | 2 | 0.529412 |
| **30** | 60 | Indoor | Yes | 3.547619 | 0 | 0.941176 |
| **31** | 16 | Indoor | No | 2.840909 | 1 | 0.705882 |
| **32** | 48 | Indoor | Yes | 2.5 | 2 | 0.529412 |
| **33** | 82 | Indoor | No | 2.869048 | 1 | 0.529412 |
| **34** | 66 | Indoor | Yes | 3.721154 | 0 | 0.941176 |
| **35** | 71 | Outdoor | Yes | 2.422414 | 3 | 0.941176 |
| **36** | 77 | Outdoor | Yes | 3.398438 | 4 | 1 |
| **37** | 34 | Outdoor | Yes | 2.683333 | 1 | 1 |
| **38** | 111 | Indoor | Yes | 3.135 | 4 | 1 |
| **39** | 38 | Indoor | Yes | 3.238095 | 4 | 0.882353 |
| **40** | 57 | Indoor | Yes | 3.163793 | 4 | 0.823529 |
| **41** | 18 | Indoor | Yes | 2.321429 | 4 | 0.588235 |
| **42** | 10 | Indoor | Yes | 2.25 | 4 | 0.588235 |
| **43** | 32 | Outdoor | Yes | 2.019231 | 0 | 0.647059 |
| **44** | 25 | Indoor | No | 2.557692 | 0 | 0.941176 |
| **45** | 45 | Indoor | No | 2.473214 | 0 | 1 |
| **46** | 18 | Indoor | No | 2.769231 | 0 | 0.529412 |
| **47** | 36 | Indoor | No | 2.945652 | 0 | 0.411765 |
| **48** | 47 | Indoor | No | 2.916667 | 0 | 0.882353 |
| **49** | 39 | Indoor | No | 3.092105 | 0 | 0.823529 |
| **50** | 30 | Indoor | Yes | 3.473684 | 0 | 0.823529 |

**Table S1. Situational variables of movie events in the discovery dataset.** *These situational variables were used in the mixed effect models to control for the item effects.*

| **ROIs** |  | ***t value*** | ***p* value** |
| --- | --- | --- | --- |
| **Early visual** | Activation | 11.44 | 4.1e-09 |
|  | Connectivity | 11.46 | 4.0e-09 |
| **Early auditory** | Activation | 9.15 | 9.3e-08 |
|  | Connectivity | 7.28 | 1.8e-06 |
| **mPFC** | Activation | 9.69 | 4.3e-08 |
|  | Connectivity | 9.40 | 6.4e-08 |
| **PMC** | Activation | 16.94 | 1.2e-11 |
|  | Connectivity | 11.82 | 2.6e-09 |
| **Hippocampus** | Activation | 3.92 | 1.2e-03 |
|  | Connectivity | 7.04 | 2.8e-06 |
| **Post. Parahipp.** | Activation | 9.15 | 9.3e-08 |
|  | Connectivity | 7.28 | 1.8e-06 |

**Table S2 Paired t-tests results for *within-event* and *between-event* activation and connectivity pattern similarities**. *Within-event* pattern similarities were higher than *between-event* pattern similarities in all ROIs investigated. Note. Bonferroni corrected significance threshold: 0.05 / 6 ROIs = 8.3e-03.

|  | **Discovery Dataset (Chen (2017))** | | **Replication Dataset (Kurby (2018))** | |
| --- | --- | --- | --- | --- |
| **ROI** | **r** | ***p*_raw_** | **r** | ***p*_raw_** |
| **Early Visual** | -0.25 | 0.07 | -0.27 | 0.04 |
| **Early Auditory** | -0.38 | 0.005 | -0.31 | 0.02 |
| **mPFC** | -0.57 | <0.001 | -0.20 | 0.15 |
| **PMC** | -0.54 | <0.001 | -0.49 | <0.001 |
| **Hippocampus** | -0.67 | <0.001 | -0.26 | 0.06 |
| **Post. Parahipp.** | -0.67 | <0.001 | -0.47 | <0.001 |

**Table S3 Correlations between activation pattern similarities and connectivity pattern similarities across event boundaries in both the discovery and replication datasets.**

| **ROIs** | **Activation pattern analysis** | | | **Connectivity pattern analysis** | | |
| --- | --- | --- | --- | --- | --- | --- |
|  | **r** | ***p****_correlation_^1^* | ***p****_paired-t test_^2^* | **r** | ***p****_correlation_* | ***p****_paired-t test_* |
| **Early visual** | -0.34 | 0.17 | 0.27 | 0.01 | 0.96 | 0.01 |
| **Early auditory** | -0.37 | 0.13 | 0.007 | 0.45 | 0.06 | 0.02 |
| **mPFC** | -0.23 | 0.37 | 0.01 | 0.24 | 0.34 | 0.23 |
| **PMC** | -0.15 | 0.55 | 0.08 | -0.11 | 0.66 | 0.02 |
| **Hippocampus** | -0.25 | 0.32 | 0.007 | 0.40 | 0.10 | 0.01 |
| **Post. Parahipp.** | -0.40 | 0.11 | 0.01 | 0.21 | 0.40 | 0.22 |

**Table S4** **Comparison between cross-participant and within-participant analyses of neural similarity and memory**. *1. P-value of the cross-participant Pearson correlation. 2. P-value of the within-participant paired t-test (i.e., main results in the main text).*

| **ROIs** | **Without covariates^1^** | | **With situational covariates^2^** | | **With perceptual covariates^3^** | | ***P*_paired t-tests_** | **Similar Pattern^4^?** |
| --- | --- | --- | --- | --- | --- | --- | --- | --- |
|  | **F** | ***p*** | **F** | ***p*** | **F** | ***p*** |  |  |
| **Early visual** | 0.93 | 0.33 | 0.73 | 0.39 | 1.03 | 0.30 | 0.27 | YES |
| **Early auditory** | 3.07 | 0.08 | 3.00 | 0.08 | 4.06 | 0.04 | 0.007 | YES |
| **mPFC** | 3.67 | 0.05 | 2.97 | 0.08 | 5.27 | 0.02 | 0.01 | YES |
| **PMC** | 1.35 | 0.24 | 1.07 | 0.30 | 2.54 | 0.11 | 0.08 | YES |
| **Hippocampus** | 3.48 | 0.06 | 2.80 | 0.09 | 4.34 | 0.03 | 0.007 | YES |
| **Post. Parahipp.** | 3.72 | 0.05 | 2.48 | 0.11 | 4.10 | 0.04 | 0.01 | YES |

**Table S5 Relationship between activation pattern similarity and memory based on mixed-effects models.** 1. Results from the raw model (i.e., formula ①). 2. Results from the second model (i.e., formula ②) where event-specific situational variables (e.g., *event duration, location, music…*) were modeled as fixed effects. 3. Results from the third model (i.e., formula ③) where perceptual variables (i.e., *corrs, dist, and lum*) were modeled as fixed effects. 4. Whether the result from the mixed –effect models showed the similar pattern as the result from the paired t-tests.

**Table S6** **Relationship between connectivity pattern similarity and memory based on mixed-effects models**. 1. Results from the raw model (i.e., formula ①). 2. Results from the second model (i.e., formula ②) where event-specific situational variables (e.g., *event duration, location, music…*) were modeled as fixed effects. 3. Results from the third model (i.e., formula ③) where perceptual variables (i.e., *corrs, dist, and lum*) were modeled as fixed effects. 4. Whether the result from the mixed –effect models showed the similar pattern as the result from the paired t-tests.

| **ROIs** | **Without covariates^1^** | | **With situational covariates^2^** | | **With perceptual covariates^3^** | | ***P*_paired t-tests_** | **Similar Pattern^4^?** |
| --- | --- | --- | --- | --- | --- | --- | --- | --- |
|  | **F** | ***p*** | **F** | ***p*** | **F** | ***p*** |  |  |
| **Early visual** | 0.12 | 0.72 | 0.03 | 0.85 | 0.03 | 0.85 | 0.01 | NO |
| **Early auditory** | 1.27 | 0.25 | 0.58 | 0.44 | 1.21 | 0.27 | 0.02 | NO |
| **mPFC** | 0.22 | 0.63 | 0.007 | 0.93 | 0.35 | 0.55 | 0.23 | YES |
| **PMC** | 0.18 | 0.67 | 0.06 | 0.80 | 0.01 | 0.90 | 0.02 | NO |
| **Hippocampus** | 3.68 | 0.05 | 2.81 | 0.09 | 3.81 | 0.05 | 0.01 | YES |
| **Post. Parahipp.** | 0.03 | 0.86 | 0.025 | 0.96 | 0.29 | 0.58 | 0.22 | YES |

| **ROIs** | **Activation Pattern Similarity** | | | | | |
| --- | --- | --- | --- | --- | --- | --- |
|  | **Recognition** | | **Recall** | | **Order** | |
|  | r value | *p*_raw_ | r value | *p*_raw_ | r value | *p*_raw_ |
| **Early Visual** | -0.08 | 0.18 | -0.06 | 0.30 | -0.10 | 0.10 |
| **Early Auditory** | -0.17 | 0.007 | -0.11 | 0.07 | -0.12 | 0.06 |
| **mPFC** | -0.14 | 0.025 | -0.13 | 0.03 | -0.19 | 0.002 |
| **PMC** | -0.17 | 0.006 | -0.17 | 0.006 | -0.16 | 0.009 |
| **Hippocampus** | -0.14 | 0.028 | -0.08 | 0.19 | -0.07 | 0.21 |
| **Post. Parahipp.** | -0.16 | 0.01 | -0.08 | 0.18 | -0.08 | 0.19 |

**Table S7 Correlation between activation pattern similarity and three memory measures in the replication dataset.**

| **ROIs** | **Connectivity Pattern Similarity** | | | | | |
| --- | --- | --- | --- | --- | --- | --- |
|  | **Recognition** | | **Recall** | | **Order** | |
|  | r value | *p*_raw_ | r value | *p*_raw_ | r value | *p*_raw_ |
| **Early Visual** | -0.04 | 0.55 | -0.07 | 0.31 | 0.03 | 0.73 |
| **Early Auditory** | -0.03 | 0.64 | 0.20 | 0.003 | 0.14 | 0.05 |
| **mPFC** | -0.05 | 0.52 | 0.15 | 0.04 | 0.12 | 0.08 |
| **PMC** | 0.02 | 0.71 | 0.26 | 0.001 | 0.14 | 0.05 |
| **Hippocampus** | -0.09 | 0.17 | 0.17 | 0.01 | 0.09 | 0.20 |
| **Post. Parahipp.** | -0.08 | 0.26 | 0.21 | 0.002 | 0.12 | 0.08 |

**Table S8 Correlation between connectivity pattern similarity and three memory measures in the replication dataset.**
